# Duplication of *NRAMP3* gene in poplars generated two homologous transporters with distinct functions

**DOI:** 10.1101/2021.12.04.471152

**Authors:** Mathieu Pottier, Van Anh Le Thi, Catherine Primard-Brisset, Jessica Marion, Michele Bianchi, Cindy Victor, Annabelle Déjardin, Gilles Pilate, Sébastien Thomine

**Author notes:** Author for correspondence Sébastien Thomine.

## Abstract

Transition metals are essential for a wealth of metabolic reactions, but their concentrations need to be tightly controlled across cells and cell compartments, as metal excess or imbalance has deleterious effects. Metal homeostasis is achieved by a combination of metal transport across membranes and metal binding to a variety of molecules. Gene duplication is a key process in evolution, as emergence of advantageous mutations on one of the copies can confer a new function. Here, we report that the poplar genome contains two paralogues encoding NRAMP3 metal transporters localized in tandem. All *Populus* species analyzed had two copies of *NRAMP3*, whereas only one could be identified in *Salix* species indicating that duplication occurred when the two genera separated. Both copies are under purifying selection and encode functional transporters, as shown by expression in the yeast heterologous expression system. However, genetic complementation revealed that only one of the paralogues has retained the original function in release of metals stored in the vacuole previously characterized in *A. thaliana*. Confocal imaging showed that the other copy has acquired a distinct localization to the Trans Golgi Network (TGN). Expression in poplar suggested that the copy of NRAMP3 localized on the TGN has a novel function in the control of cell-to-cell transport of manganese. This work provides a clear case of neo-functionalization through change in the subcellular localization of a metal transporter as well as evidence for the involvement of the secretory pathway in cell-to-cell transport of manganese.

## INTRODUCTION

Several transition metals are essential cofactors for a wealth of metabolic reactions in all living organisms. Iron (Fe) and copper (Cu) are for example needed in large amounts for the respiratory electron transfer chains and ATP production in bacteria and mitochondria. Transition metals are also important for DNA synthesis, proteolysis and the control of reactive oxygen species (ROS). Photosynthetic organisms have an additional specific requirement for manganese (Mn) for light energy conversion and water splitting (Shen 2015). Although they are essential, transition metal concentrations need to be tightly controlled across cells and cell compartments, as excess or imbalance between different metals has deleterious effects. Metal homeostasis is achieved by a combination of metal transport across membranes and metal binding to a variety of molecules, including proteins, small peptides, amino acids, organic acids and specialized metabolites, such as phytochelatins or nicotianamine in plants (Seregin and Kozhevnikova 2021).

Many families of transporters are implicated in the maintenance of metal homeostasis, either for metal uptake, distribution of metals to organs within organisms and to organelles within a cell, or for removal and sequestration of excess metal. For example, in *Arabidopsis thaliana*, Mn is taken up by AtNRAMP1 (Natural Resistance Associated Macrophage Protein 1) in the roots and distributed within cells by AtNRAMP2 (Cailliatte et al. 2010; Alejandro et al. 2017; Gao et al. 2018). AtMTP8 (Metal Tolerance Protein 8) and AtMTP11, which belong to a different transporter family, are responsible for loading Mn from the cytosol into the vacuole or the Trans Golgi Network (TGN), respectively (Delhaize et al. 2007; Peiter et al. 2007; Eroglu et al. 2016). The vacuole is used to store Mn excess and prevent its toxicity. However, when this element becomes scarce, other NRAMP family members, namely AtNRAMP3 and AtNRAMP4, allow the retrieval of Mn from the vacuole (Lanquar et al. 2010). Mn is needed in the secretory system as a cofactor of glycosyl transferases involved in protein glycosylation (Alejandro et al. 2020). It also plays important roles as a cofactor of superoxide dismutase in mitochondria and peroxisomes (Alejandro et al. 2020). Moreover, Mn is essential for oxygenic photosynthesis as a component of the Mn_4_CaO_5_ cofactor of the water splitting complex, which is bound to photosystem II (PS II) at the inner side of the thylakoid membranes (Shen 2015). CMT1 (Chloroplast Manganese Transporter 1) and PAM71 (Photosynthesis-affected mutant 71), two transporters belonging to the GDT1 family, have been shown to allow the import of Mn across the inner membrane of the chloroplast envelope and the thylakoid membrane, respectively (Schneider et al. 2016; Eisenhut et al. 2018; Zhang et al. 2018). Recently, another member of the GDT1/UPF0016 family was shown to play a crucial role in loading Mn in the Golgi apparatus, where it is needed as a cofactor of glycosyl transferases involved in cell wall formation (Yang et al. 2021). The networks of transporters that mediate uptake, storage and distribution of other essential metals, such as Fe, Zn and Cu, have also been described. Interestingly, these networks are interconnected, as some transporters, as well as ligands, are able to transport a broad range of metal cations (Pottier, Oomen, et al. 2015; Seregin and Kozhevnikova 2021).

This is well illustrated when looking at the functions of transporters of the NRAMP family. This family was first identified in the context of resistance to intracellular pathogens, such *Mycobacterium tuberculosis*, in mammals. Murine NRAMP1 was shown to limit the growth of intracellular pathogens by depleting essential metals from the phagosomes where they reside (Vidal et al. 1993; Wessling-Resnick 2015). Mammalian NRAMP2 plays a central role in Fe absorption in the intestine. In yeast, the NRAMP members SMF1 and SMF2 are involved in Mn absorption and distribution, similar to *A. thaliana* NRAMP1 and NRAMP2 (Portnoy et al. 2000; Cailliatte et al. 2010; Alejandro et al. 2017; Gao et al. 2018), while SMF3 allows the release of Fe from the vacuole, similar to *A. thaliana* NRAMP3 and NRAMP4 (Portnoy et al. 2000; Lanquar et al. 2005). In *A. thaliana*, NRAMP1 does not only allow high affinity Mn uptake but also plays a role in low affinity Fe uptake (Castaings et al. 2016), clearly illustrating that these transporters connect Fe and Mn homeostasis.

While many studies have addressed the molecular mechanisms of metal homeostasis in *A. thaliana* and rice, there are only a limited number of reports on this topic in poplar. Poplars are both model trees for which the genome of several species has been sequenced (Tuskan et al. 2006; Lin et al. 2018; Zhang et al. 2019), and an industrially important crop for wood production. Poplars display outstanding growth yield among tree species and their wood is mostly used by the peeling industry to produce light packaging and plywood. Moreover, poplars are often used in the rehabilitation of polluted areas because they are highly tolerant to heavy metals and other pollutants (Krämer 2005a; Pottier, García de la Torre, et al. 2015). Poplar MTP family members have been functionally investigated. In the first sequenced poplar species *Populus trichocarpa* (Tuskan et al. 2006), it has been shown that PotriMTP1 and PotriMTP11 could be the functional homologues of AtMTP1 and AtMTP11 and are involved in Zn loading into the vacuole and Mn loading into the TGN, respectively (Blaudez et al. 2003; Krämer 2005b; Peiter et al. 2007). Copper homeostasis has also been investigated in the context of photosynthetic efficiency (Ravet et al. 2011). In addition, over-expression of genes involved in Zn and Cd chelation and homeostasis has been undertaken in an attempt to increase tolerance and accumulation of these metals (Adams et al. 2011; He et al. 2015; Wang et al. 2019). Several studies have mined poplar genomic data and established lists of metal transport proteins in this species, analyzed the expression pattern of the corresponding genes and sometimes demonstrated the transport function using yeast complementation (Migeon et al. 2010; Li et al. 2015; Gao et al. 2020). These studies have often highlighted the presence of duplication in metal homeostasis genes, which is a prevalent feature in poplar genome (Tuskan et al. 2006). However, they have not investigated in detail the function of the duplicated copies.

Gene duplication is a key process in evolution. It occurs through two major processes: either whole genome duplication or local duplication by unequal crossover or transposition (Conant and Wolfe 2008). Gene duplication underlies several key events in evolution such as variation in gene copy number, the generation of new regulatory networks and the appearance of novel functions. After gene duplication occurs, relaxation of selective pressure opens the door to several scenarios. Most of the time, one of the copies undergoes non-functionalization through accumulation of deleterious mutations due to the lack of selective pressure on this copy. In other cases, having multiple copies of the same gene provides advantages and several functional identical genes are therefore actively maintained (Hanikenne et al. 2013). Often the copies can also undergo sub-functionalization: the preduplication function is maintained but partitioned between the two copies. Typically, the expression pattern of the ancestral gene is covered by the two copies which are expressed in different organs and involved in distinct regulatory networks (Tuskan et al. 2006). Finally, in rare cases, emergence of advantageous mutations on one of the copies can also confer a new function, which is commonly known as neo-functionalization (Moriyama et al. 2016).

In this study, we have investigated the function of poplar *NRAMP3*. We found that the poplar genome contains two paralogues of *NRAMP3* in tandem. One of the paralogues has conserved the original function in release of metals stored in the vacuole characterized in *A. thaliana*, whereas the other paralogue has acquired a distinct localization to the TGN. Analysis of the function of this gene in transgenic poplars suggests that it has a novel function in the control of cell-to-cell transport of Mn. Therefore, the functional analysis of the two paralogues of *PotriNRAMP3* provides a clear case of neo-functionalization through change in the subcellular localization of a transporter and evidence for the involvement of the secretory pathway in cell-to-cell transport of Mn.

## RESULTS

### *PotriNRAMP3*.*1* and *PotriNRAMP3*.*2* are a tandem gene pair encoding homologous proteins

The eleven NRAMPs retrieved from *P. trichocarpa* genome V4.1 distribute into the three different phylogenetic groups of plant NRAMPs defined according to their protein sequence identities and exon-intron structures (Migeon et al. 2010, table S1; fig. S1A). PotriNRAMP1, PotriNRAMP6.1 and PotriNRAMP6.2 as well as PotriNRAMP7.1, PotriNRAMP7.2 and PotriNRAMP7.3 belong to group I, which also includes AtNRAMP1 and AtNRAMP6. PotriNRAMP2, PotriNRAMP3.1 and PotriNRAMP3.2 are in group II, which includes AtNRAMP2, AtNRAMP5, AtNRAMP3 and AtNRAMP4. PotriEIN2.1 and PotriEIN2.2 are located in the group III as AtEIN2 (fig. S1A).

Interestingly, *PotriNRAMP3*.*1* and *PotriNRAMP3*.*2* genes are localized in close vicinity in *P. trichocarpa* genome. They are positioned in a 32 kb area of chromosome 7 and encode 88.2% identical proteins (fig. S1B-C). Dot plot analyses performed on genomic DNA including *PotriNRAMP3*.*1* and *PotriNRAMP3*.*2* genes show sequence conservation specifically between *NRAMP3* loci, more precisely between their coding sequences and their 3’UTR, while no conservation is observed between introns (fig. S1C-D).

### *NRAMP3.1* and *NRAMP3*.*2* are present in all sequenced poplar species but not in closely related species

To determine how widespread the duplication of *NRAMP3* observed in *P. trichocarpa* genome is, sequences similar to *PotriNRAMP3*.*1* and *PotriNRAMP3*.*2* were retrieved using reciprocal BLASTs in nine genomes and four transcriptomes of *Populus* species, covering the five main *Populus* sections, *i*.*e*., Tacahamaca, Populus, Leucoïdes, Turanga, Aigeiros and Abaso (Zhang et al. 2019; Wang et al. 2020). We could unequivocally identify two distinct sequences similar to *PotriNRAMP3*.*1* or *PotriNRAMP3*.*2*, in all investigated *Populus* genomes, *i*.*e*., *P. alba, P. cathayana, P. simonii, P. lasiocarpa, P. maximowiczii, P. euphratica, P. ussuriensis, P. nigra, P. deltoides, P. tremula, P. tremuloides* and *P. grandidentata* (table S1 and supplementary data S1) (Zhang et al. 2019). Evidence for two distinct sequences similar to *PotriNRAMP3*.*1* or *PotriNRAMP3*.*2* was also observed in *P. mexicana*, the single living species of the ancestral poplar section Abaso, even though its genome is not fully sequenced yet (table S1 and supplementary data S2; Wang et al. 2020). This result suggests that distinct *PotriNRAMP3*.*1* and *PotriNRAMP3*.*2* homologues are present in all *Populus* species. In contrast, blasting *PotriNRAMP3*.*1* and *PotriNRAMP3*.*2* on the genomes of three Salix species (*S. purpurea, S. suchowensis* and *S. brachista*) and on the transcriptomes of five other Salix species (*S. viminalis, S. sachalinensis, S. eriocephala, S. fargesii* and *S. dasyclados*) that belong to the closest phylogenetic group to *Populus* genus (Chen et al. 2019) identified a single *NRAMP3* sequence in each species (table S1 and supplementary data S1). The reciprocal BLASTs performed on *P. trichocarpa* genome provided *PotriNRAMP3*.*2* as best hit. These results suggest that the duplication that gave rise to *NRAMP3*.*1* and *NRAMP3*.*2* genes coincided with the divergence between *Populus* and *Salix* about 52 million years ago (Ma) (Hou et al. 2016). However, the chromosomal rearrangements that distinguish *Populus* and *Salix* genus did not affect chromosome 7, which carries the *NRAMP3* loci (Hou et al. 2016). Moreover, gene collinearity is maintained around *NRAMP3* loci in *P. trichocarpa* and *S. purpurea* (fig. 1; fig. S2). Using the corresponding protein sequences of all the *Populus* and *Salix* NRAMP3 homologues identified during this analysis (supplementary data S3), we constructed a phylogenetic tree (fig. 2). This analysis showed that *Populus* NRAMP3.1 and NRAMP3.2 form distinct phylogenetic groups (fig. 2). The *Salix* homologues clearly cluster together with *Populus* NRAMP3.2 indicating that it corresponds to the ancestral copy, whereas *Populus* NRAMP3.1 sequence diverged.

**Fig. 1.**
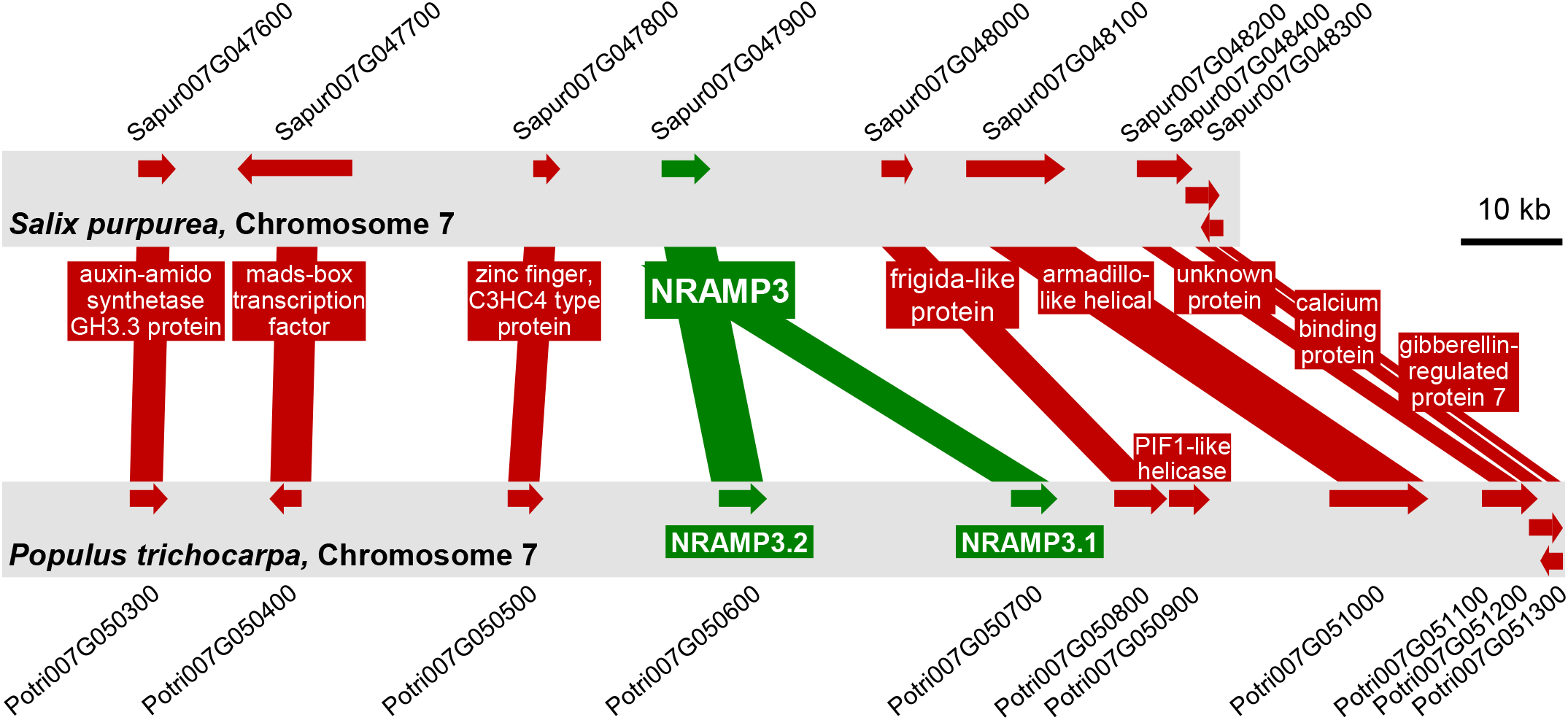
Gene collinearity between *Salix purpurea* and *Populus trichocarpa* genomes is maintained in the area surrounding *NRAMP3* locus which is specifically duplicated in poplars. Schematic representation of the genomic sequence around *NRAMP3* loci in *P. trichocarpa* and *S. purpurea*. Gene collinearity and poplar-specific NRAMP3 duplication are supported by dot-plots analysis performed with *S. purpurea* and *P. trichocarpa* genomic sequences surrounding *NRAMP3* loci (fig. S2).

**Fig. 2.**
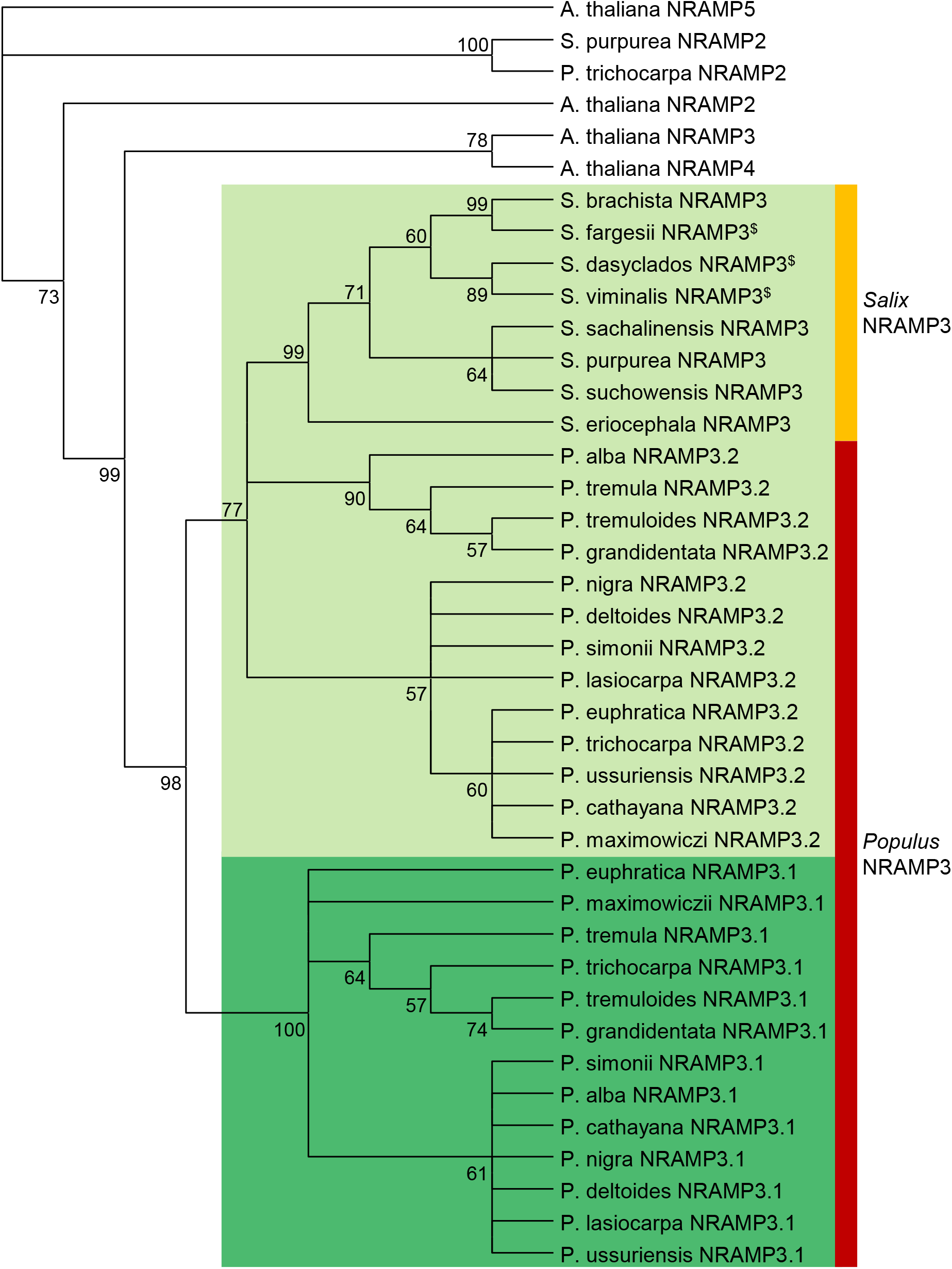
Phylogenetic tree of NRAMP3 homologues in *Populus* and *Salix* species. *A. thaliana* NRAMP2 (AT1G47240.1), NRAMP3 (AT2G23150.1), NRAMP4 (AT5G67330.1) and NRAMP5 (AT4G18790 .1) as well as *P. trichocarpa* NRAMP2 (Potri002G121000.1) and *S. purpurea* NRAMP2 (Sapur.002G098200.1) are shown as an outgroup. The other protein sequences used for this tree are listed in supplementary data S3. ^$^ indicates incomplete protein sequences. Phylogenetic analyses were conducted as described in Materials and methods.

To further analyze the evolutionary history of *Populus* NRAMP3.1 and NRAMP3.2 sequences, we calculated the ratio of non-synonymous (dN) vs synonymous codons (dS), between all NRAMP3.1s, between all NRAMP3.2s, and between all NRAMP3.1s and NRAMP3.2s together. Low global dN/dS around 0.2 were obtained for NRAMP3.1 and NRAMP3.2 indicating that both genes are under purifying selection. A sliding window analysis revealed that the low global dN/dS values obtained among NRAMP3.1 or NRAMP3.2 sequences result from homogeneously low ratio values along their open reading frames (fig. S3). In contrast, comparing NRAMP3.1s with NRAMP3.2s revealed heterogeneous values along the open reading frame, with ratios above or close to 1 in the N and C termini, as expected from divergent sequences. These results indicate that relaxation of selective pressure led to sequence diversity after the duplication event, but that both genes are now under purifying selection. Fixed Effects Likelihood (FEL) method was then employed to investigate site specific selective pressures specifically applied to either NRAMP3.1s or NRAMP3.2s. In this way, 21 and 4 residues under purifying selection (p < 0.05) were identified in NRAMP3.1s and NRAMP3.2s, respectively (tables S2 and S3). These residues are highlighted on an alignment between the consensus sequence of *Populus* NRAMP3.1 and that of *Populus* NRAMP3.2 generated from all the *Populus* NRAMP3.1 and NRAMP3.2 sequences retrieved in this study (fig. 3; supplementary data S3 and fig. S4). Note that the four residues under purifying selection in NRAMP3.2s are also under purifying selection in NRAMP3.1s. Moreover, with the exception of V491, amino acids under purifying selection in NRAMP3.1s are conserved in NRAMP3.2s. These results suggest essential roles of these residues in the basal NRAMP3 function, common to both NRAMP3.1 and NRAMP3.2. In contrast, distinct residues were found to be under positive selection in both NRAMP3.1 (positions 3 and 437, p < 0.05) and NRAMP3.2 (positions 161 and 265, p < 0.05) (fig. 3; tables S2 and S3). These analyses indicate that NRAMP3.1 and NRAMP3.2 are not under non-functionalization; both sequences continue to diversify, even though they are globally under strong purifying selection. Moreover, transcript level analysis showed that *NRAMP3*.*1* and *NRAMP3*.*2* are both expressed in roots, stems, buds and leaves, arguing against subfunctionalization by partitioning their expression in distinct organs of poplar (fig. S5).

**Fig. 3.**
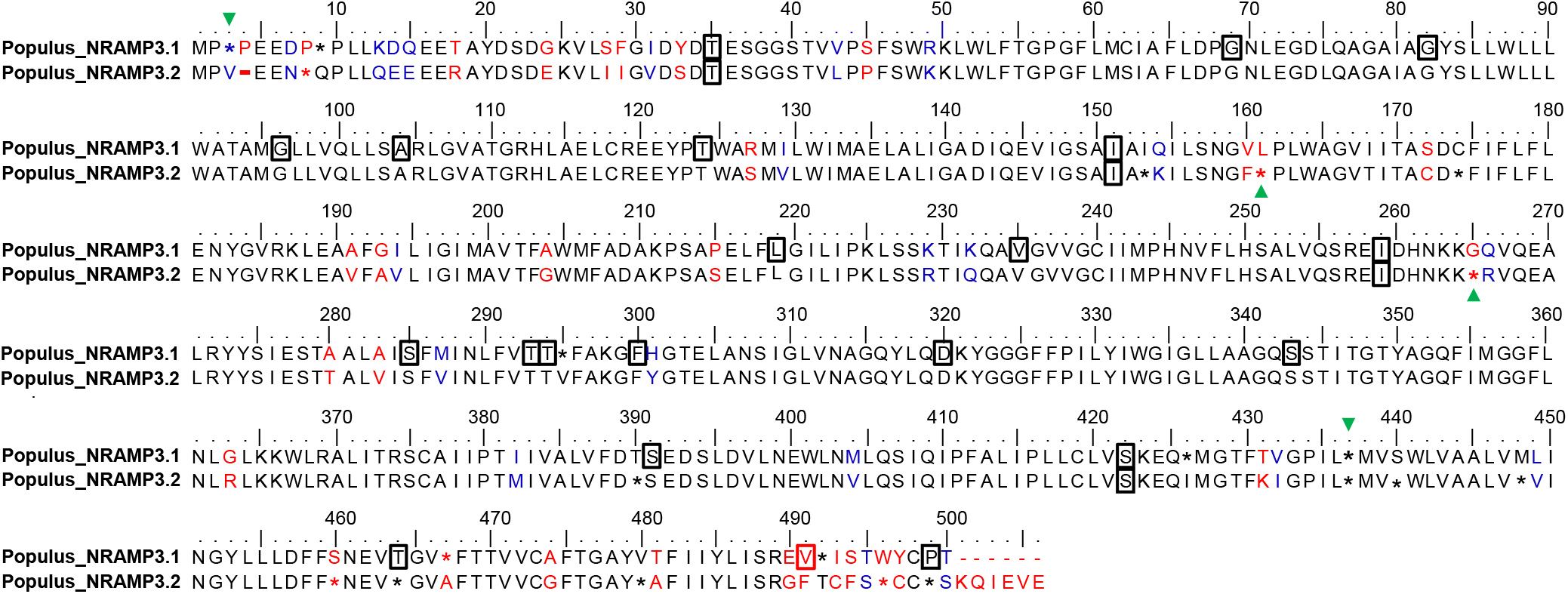
Alignment of the consensus sequences of *Populus* NRAMP3.1 and NRAMP3.2. The consensus sequences were determined from the NRAMP alignments shown in supplementary figure S3. Asterisk indicates less than 90% of conservation within *Populus* NRAMP3.1 or NRAMP3.2 cluster. Identical, similar and different residues between the two consensus sequences are indicated in black, blue, and red, respectively. Frames indicate residues under purifying selection and green arrows indicate positive selection in either *Populus* NRAMP3.1 or NRAMP3.2 cluster according to the FEL method (p < 0.05). Note that NRAMP3.1 sequences contain an insertion of residue at position 4, leading to a gap (-) in NRAMP3.2 sequences.

### Both PotriNRAMP3.1 and PotriNRAMP3.2 encode functional metal transporters

To examine the functions of the two paralogues, we cloned *P. trichocarpa NRAMP3*.*1* and *NRAMP3*.*2* cDNAs and expressed them in yeast. Because several plant NRAMP were previously shown to function in Mn homeostasis, we investigated PotriNRAMP3s ability to transport this metal. For this purpose, we tested whether they could complement the s*mf1* and *smf2* Mn transporter yeast mutants, which are unable to grow on low Mn condition (Cohen et al. 2000). AtNRAMP1 and AtNRAMP2, which are the functional homologues of Smf1p and Smf2p, respectively, were used as positive controls (Supek et al. 1996; Luk and Culotta 2001; Cailliatte et al. 2010; Alejandro et al. 2017). The β-glucuronidase enzyme (GUS) that has no transport activity was used as a negative control. We found that the expression of *PotriNRAMP3*.*2* restored *smf1* growth on low Mn condition to the same extent as *AtNRAMP1* (Thomine et al. 2000), whereas the expression of *PotriNRAMP3*.*1* allowed only a partial complementation (fig. 4A). In contrast, the expression of *PotriNRAMP3*.*1, PotriNRAMP3*.*2* or *AtNRAMP2* fully complemented *smf2* growth defect in this condition (fig. 4B). We then took advantage of the low Mn concentration in *smf2* mutant cells to investigate the effect of *PotriNRAMP3*.*1* and *PotriNRAMP3*.*2* expression on Mn accumulation. We used AtNRAMP4 as a positive control, as expression of this homologue of PotriNRAMP3s was previously shown to enhance Mn accumulation in yeast (Pottier, Oomen, et al. 2015). We found that expression of *PotriNRAMP3*.*1, PotriNRAMP3*.*2* or *AtNRAMP4* significantly increased Mn concentration in the yeast mutant (fig. S6). Mn concentration was 14 and 8 times higher in *PotriNRAMP3*.*1* and *PotriNRAMP3*.*2* expressing *smf2* strains than in *smf2* GUS control (fig. S6). Thus, both copies of PotriNRAMP3 have retained metal transport ability. However, differences in complementation efficiency and metal accumulation suggest differences in transport capacity or localization between these two transporters. Besides, complementation assays of the *fet3fet4* yeast strain deficient for both low-and high-affinity Fe uptake systems indicated that PotriNRAMP3.1 and PotriNRAMP3.2 are able to transport Fe in addition to Mn (fig. S7), as previously shown for AtNRAMP3 and AtNRAMP4 (Thomine et al. 2000).

**Fig. 4.**
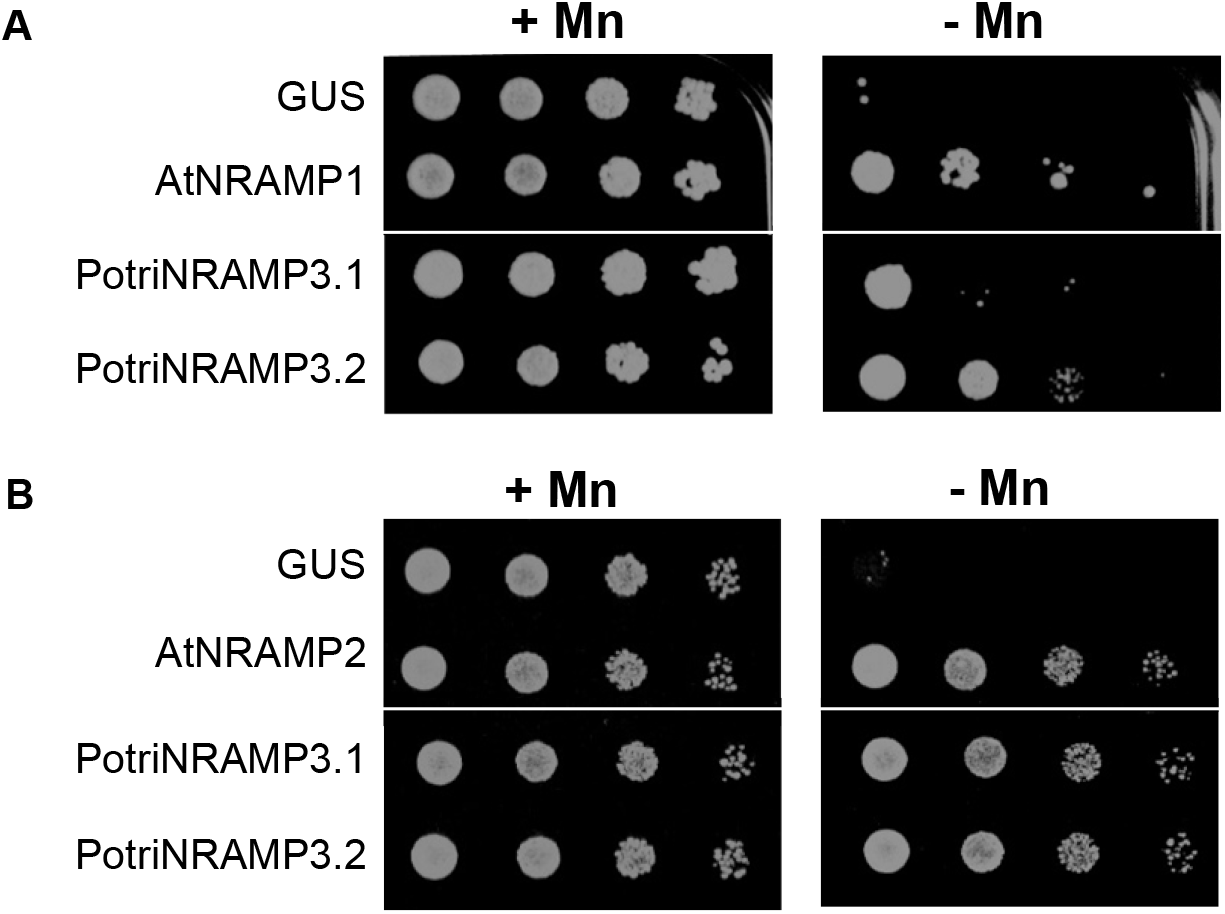
*PotriNRAMP3*.*1* and *PotriNRAMP3*.*2* encode functional Mn transporters. Functional complementation of *smf1* (A) and *smf2* (B) yeast mutants. Yeast cells expressing *GUS* (negative control), *AtNRAMP1* (positive control), *AtNRAMP2* (positive control), *PotriNRAMP3*.*1* or *PotriNRAMP3*.*2* were grown overnight. The cultures were diluted to ODs of 1 to 10^−3^ and spotted on synthetic dextrose -ura plates. Transformed *smf1* (A) strains were grown on medium supplemented with 5 mM EGTA and 100 µM MnSO_4_ (+ Mn) or with 5 mM EGTA without MnSO_4_ (-Mn). Transformed *smf2* (B) strains were grown in medium supplemented with 10 mM EGTA and 100 µM MnSO_4_ (+ Mn) or with 10 mM EGTA without MnSO4 (-Mn). The plates were incubated at 30°C for 5 days (*smf1*) or 2 days (*smf2*) before photography. White lines indicate cropping.

### *PotriNRAMP3*.*2*, but not *PotriNRAMP3*.*1*, complements the *nramp3nramp4* double mutant of *A. thaliana*

Because both PotriNRAMP3s share high protein sequence identity with AtNRAMP3 and AtNRAMP4, we tested whether they could perform the same function *in planta*. AtNRAMP3 and AtNRAMP4 have redundant functions in Fe remobilization from vacuoles during seed germination. As a consequence, *A. thaliana nramp3nramp4* double mutants are sensitive to Fe starvation during their early development (Lanquar et al. 2005). We expressed *PotriNRAMP3*.*1* and *PotriNRAMP3*.*2* under the *Ubiquitin 10* (*pUb10*) promoter in the *A. thaliana* Columbia 0 (Col-0) *nramp3nramp4* mutant background (Grefen et al. 2010; Bastow et al. 2018), and selected 3 independent homozygous T3 lines expressing transgenes at various levels for further experiments (fig. S8). Lines transformed with *pUb10:PotriNRAMP3*.*2* exhibited full complementation of the double mutant phenotype on Fe deficient medium: root length and cotyledon greening were indistinguishable from wild-type (fig. 5). Complementation was observed irrespective of the expression level of the transgene, indicating that even low levels are sufficient to restore the wild-type phenotype. In contrast, the expression of *PotriNRAMP3*.*1* did not improve the growth of the *A. thaliana nramp3nramp4* double mutant under Fe deficient conditions, even in lines showing high expression of the transgene (fig. 5, S8). Similar results were obtained when expressing *PotriNRAMP3*.*1-GFP* and *PotriNRAMP3*.*2-GFP* under the CaMV 35S promoter (*p35S*) to study their subcellular localization in *A. thaliana* and poplar (see below), except that *p35S:PotriNRAMP3*.*1-GFP* partially improved *nramp3nramp4* growth under Fe deficiency (fig. S9). Even though PotriNRAMP3.1 is able to transport Fe (fig. S7), it is thus not able to restore Fe remobilization during seed germination.

**Fig. 5.**
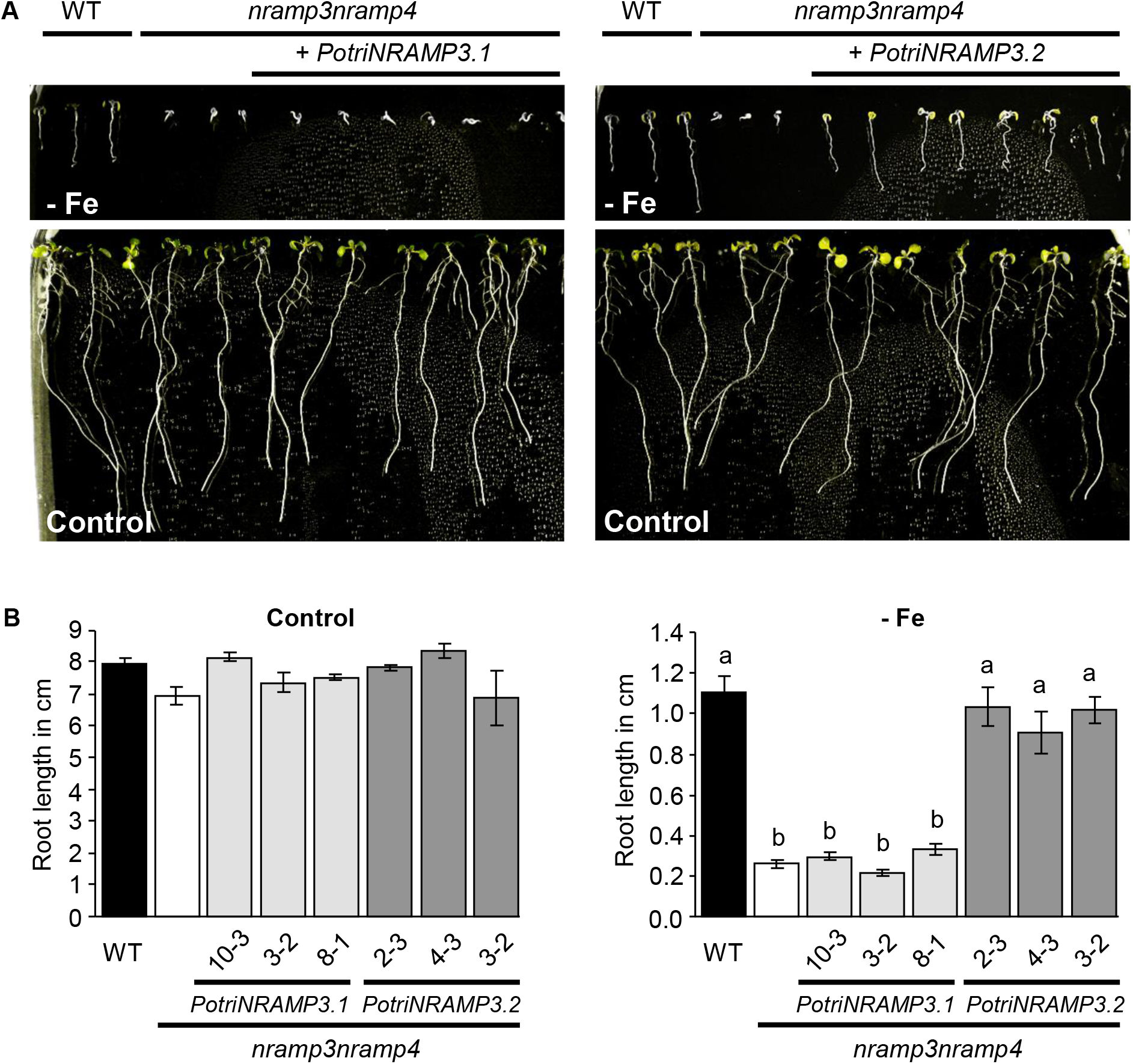
*PotriNRAMP3*.*2* but not *PotriNRAMP3*.*1* expression complements the Arabidopsis *nramp3nramp4* double mutant growth defects under iron starvation. (A) Representative pictures of *nramp3nramp4 pUb10:PotriNRAMP3*.*1 and pUb10:PotriNRAMP3*.*2* T3 Arabidopsis lines together with wild-type (Col-0) as positive control and *nramp3nramp4* as negative control grown vertically for 8 days in ABIS supplemented with 50 μM FeHBED (control, bottom panel) or without iron (- Fe, top panel). (B) Quantification of main root lengths of wild-type (Col-0), *nramp3nramp4* and 3 independent *nramp3nramp4 pUb10:PotriNRAMP3*.*1 and pUb10:PotriNRAMP3*.*2 T3* lines. Values represent mean of 10 - 12 roots and bars represent SD. Different letters reflect significant differences according to a Kruskal-Wallis test followed by Dunn’s test for multiple comparison (*p < 0*.*01*). No significant differences among genotypes were detected in the presence of Fe, left panel in (B).

### PotriNRAMP3.1 and PotriNRAMP3.2 have distinct subcellular localizations in plant cells

Change in intracellular localization is one of the mechanisms leading to neofunctionalization (Ren et al. 2014). To examine PotriNRAMP3.1 and PotriNRAMP3.2 subcellular localizations, *A. thaliana* and poplar transgenic lines expressing C-terminal GFP fusion proteins of these two transporters were generated. Previous studies showed that tagging with GFP at the C-terminal end does not affect NRAMP targeting and function in plants (Lanquar et al. 2005; Cailliatte et al. 2010; Alejandro et al. 2017). The *A. thaliana nramp3nramp4* double mutant (Col-0) was stably transformed with *p35S:PotriNRAMP3*.*1-GFP* and *p35S:PotriNRAMP3*.*2-GFP*. Roots of these plants were then observed by confocal microscopy (fig. 6A). Interestingly, distinct subcellular localizations were observed for PotriNRAMP3.1-GFP and PotriNRAMP3.2-GFP. While PotriNRAMP3.2-GFP was targeted to the vacuolar membrane, as its homologues in *A. thaliana* and *Noccaea caerulescens* (Thomine et al. 2003; Lanquar et al. 2005; Oomen et al. 2009), PotriNRAMP3.1 was localized in intracellular punctuate structures (fig. 6A). The localization of PotriNRAMP3.2 on the vacuolar membrane is consistent with its ability to complement the *A. thaliana nramp3nramp4* double mutant. The partial complementation observed with PotriNRAMP3.1-GFP could be due to the mis-targeting of a small fraction of this protein, too low to generate detectable fluorescence, to the vacuolar membrane. Similar PotriNRAMP3.1 and PotriNRAMP3.2 localizations were observed in poplar root cells (fig. 6A) as well as in mesophyll protoplasts and leaf epidermal cells (fig. S10).

**Fig. 6.**
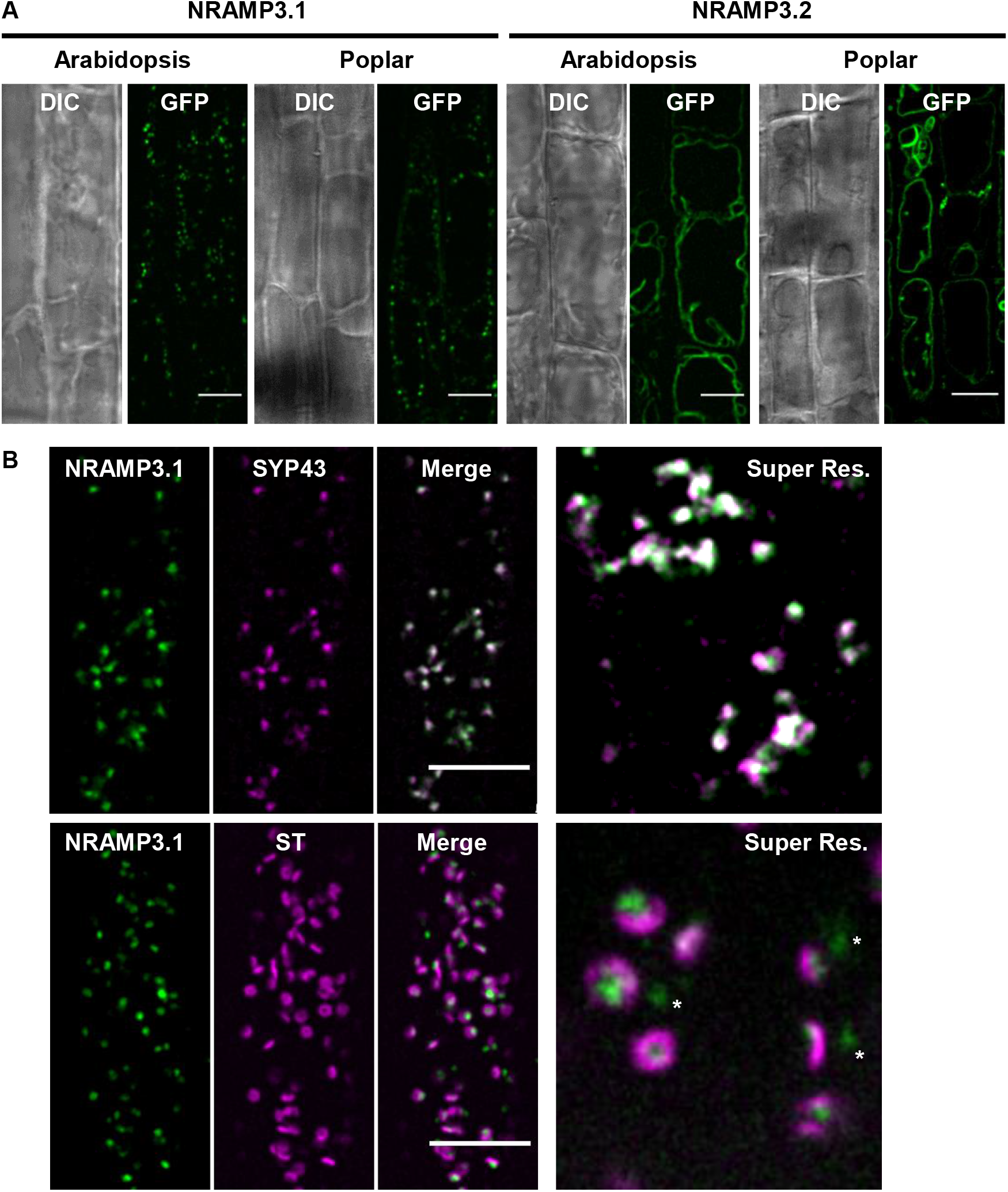
PotriNRAMP3.1 localizes to the Trans-Golgi Network (TGN) while PotriNRAMP3.2 localizes to the vacuolar membrane. (A) GFP translational fusions of PotriNRAMP3.1 and PotriNRAMP3.2 were imaged in vacuolar planes by spinning disk confocal microscopy in root epidermal cells (early elongation zone) of transgenic Arabidopsis T3 seedlings and poplar. NRAMP3.1-GFP labels granular cytoplasmic structures at the cell periphery, in both Arabidopsis and Poplar. In contrast, NRAMP3.2-GFP is present on the vacuolar membrane. Transmitted-light (differential interference contrast, DIC) and fluorescence (GFP) acquisitions are shown. (B) Colocalization of PotriNRAMP3.1-GFP with the TGN marker mRFP-SYP43 (*top panel*) and juxtaposition of PotriNRAMP3.1-GFP with the trans-Golgi apparatus marker mRFP-ST (*bottom panel*) in root epidermal cells (early elongation zone) of Arabidopsis F1 seedlings. Translational fusions were imaged in the cortical planes by spinning disk confocal. Note how PotriNRAMP3.1-GFP fluorescence either faces the center of the toroidal structure of the trans-Golgi or is present as Golgi-independent structures (white asterisks). On the merged images the overlap of GFP (green) and mRFP (magenta) channels appears white. Scale bar : 10 µm. Super resolution acquisitions (Super Res.) are 10 µm wide.

To determine more precisely the subcellular localization of PotriNRAMP3.1, we tested the colocalization of PotriNRAMP3.1-GFP with RFP markers for different cell compartments (Ebine et al. 2011; Uemura et al. 2012; Inada et al. 2016). To this aim, *A. thaliana* lines expressing PotriNRAMP3.1-GFP were crossed with stable lines expressing markers for the trans Golgi apparatus *i*.*e*., mRFP-ST (Sialyl Transferase), the TGN *i*.*e*., mRFP-Syp43 and two endosomal markers *i*.*e*., ARA6-mRFP and ARA7-mRFP. Spinning disk confocal microscopy performed on the F1 seedlings showed an extensive overlap between PotriNRAMP3.1-GFP and mRFP-SYP43 fluorescence (fig. 6B). Interestingly, although PotriNRAMP3.1-GFP fluorescence did not overlap with that of mRFP-ST, it was most often in close vicinity (fig. 6B). In contrast, little or no colocalization was observed with endosomal markers (fig. S11). These colocalization experiments indicate that PotriNRAMP3.1 resides on the TGN, and that it is present in both Golgi-associated and Golgi-independent TGN compartments (Viotti et al. 2010; Uemura et al. 2019). Together, these results show that the *PotriNRAMP3*.*2* copy has retained the subcellular localization and function of the *NRAMP3* genes characterized in other species, whereas *PotriNRAMP3*.*1* has likely acquired a novel function due to mutations that modified its subcellular localization to the TGN. However, *PotriNRAMP3*.*1* expression in *A. thaliana* did not lead to any phenotypic alteration that could provide hints at this novel function.

### PotriNRAMP3.1, but not PotriNRAMP3.2, affects manganese homeostasis in poplar

To investigate the functions of PotriNRAMP3.1 and PotriNRAMP3.2 in poplar, the genes coding these transporters were over-expressed as GFP fusions under the control of the *p35S*. For each construct, four independent transgenic lines over-expressing (OE) the *PotriNRAMP3s* at levels 10 to 25 times higher than non-transgenic (NT) control trees were analyzed (fig. S12). Poplar lines with high levels of *PotriNRAMP3*.*1* expression displayed reduced height as well as interneval chlorosis on mature leaves compared to NT control trees (fig. 7A, C, F; fig. S12). In contrast, *PotriNRAMP3*.*2* OE trees were indistinguishable from NT poplars (fig. 7B, C, H). To better understand the origin of the chlorosis, the maximum quantum yield of PS II was imaged using an Imaging PAM (Walz, Germany) in leaves of control poplars as well as *PotriNRAMP3*.*1* and *PotriNRAMP3*.*2* OE lines (fig. 7E, G, I). The chlorotic areas in *PotriNRAMP3*.*1* OE lines coincided with strongly decreased PS II maximum quantum yield (0.472±0.035). In contrast, PS II efficiency was close to the optimal value of 0.82 in leaves from control (0.756±0.002) and *PotriNRAMP3*.*2* OE trees (0.751±0.008). As internerval chlorosis is a symptom of Fe deficiency and decrease in PS II efficiency may be a symptom of Mn deficiency (Connorton et al. 2017; Alejandro et al. 2020), we quantified metals in young, mature and senescent leaves from the different poplar genotypes. These analyses revealed that Mn concentrations in young and mature leaves were significantly lower in PotriNRAMP3.1 OE lines compared to NT control or PotriNRAMP3.2 OE lines (fig. 8A, B, C). In contrast, no significant difference in Fe or Zn concentrations was detected among the different genotypes (fig. S13). Interestingly, opposite to what was observed in leaves, Mn concentrations in stems of PotriNRAMP3.1 OE lines were higher than in NT control or PotriNRAMP3.2 OE lines (fig. 8D). Mn concentrations were also significantly higher in stems of PotriNRAMP3.2 OE lines compared to NT control. The defect in Mn distribution observed in PotriNRAMP3.1 OE lines suggests that the phenotypes observed in these lines are due to a defect in Mn transfer from stems to leaves leading to limited Mn supply to leaves.

**Fig. 7.**
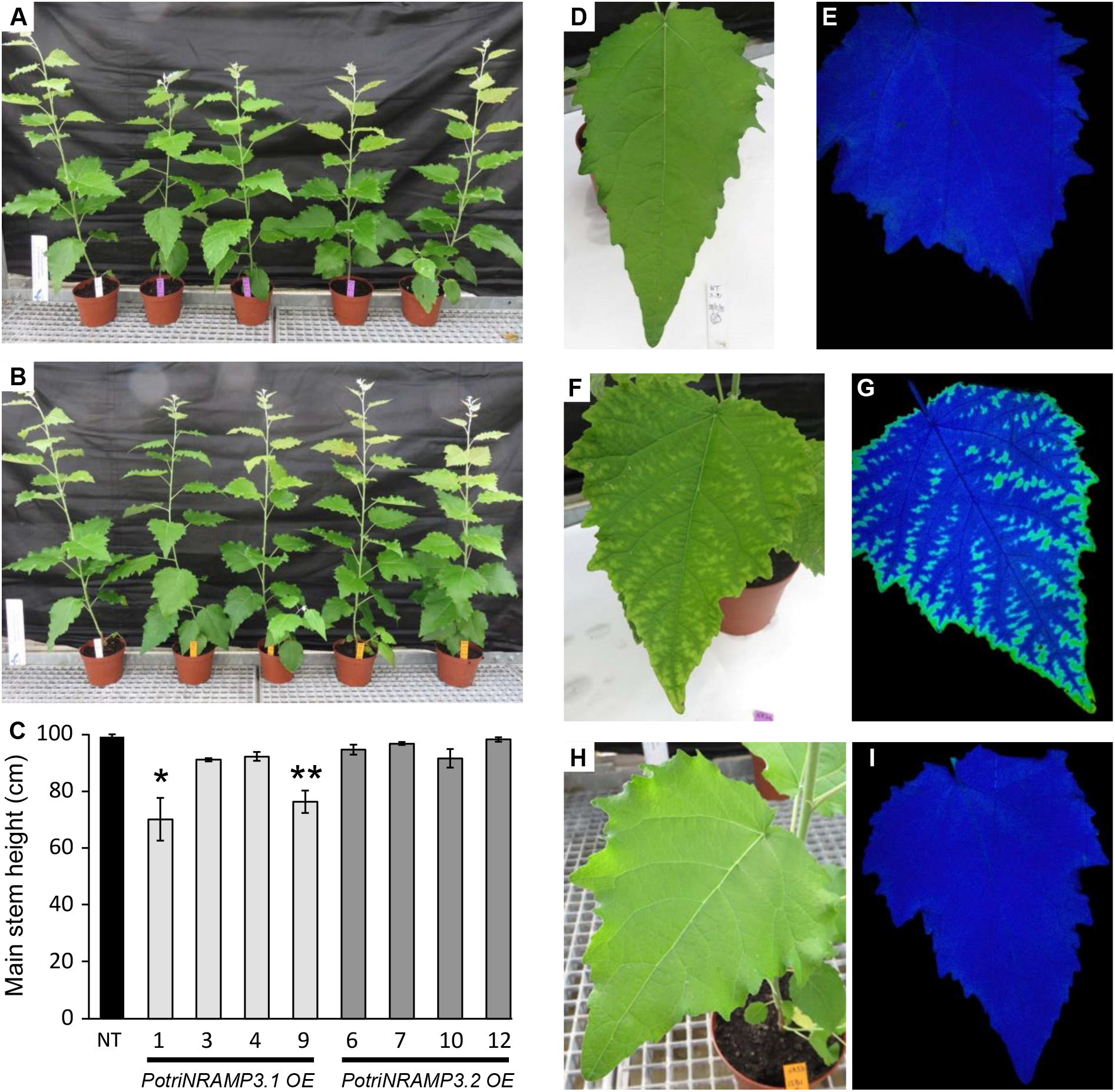
The ectopic over expression of *PotriNRAMP3*.*1* but not that of *PotriNRAMP3*.*2* leads to phenotypic alterations in poplar. Overview of 4 independent transgenic poplar lines over-expressing *PotriNRAMP3*.*1-GFP* (A, purple tags) or *PotriNRAMP3*.*2-GFP* (B, orange tags) along with NT control (A and B, white tag), 2 months after transfer from *in vitro* to soil. (C) Mean heights of poplar from the different genotypes. Error bars represent SE (n = 4-7 trees per genotype). Asterisks denote significant difference with respect to NT control according to a Mann-Whitney test (*: *p < 0*.*05*, **: *p < 0*.*01*). (D-I) Leaf phenotypes of representative trees. (D,E) NT control, (F,G) PotriNRAMP3.1-GFP line 9 (H,I), PotriNRAMP3.2-GFP line 12. (D, F, H) pictures; (E, G, I) PS II maximum quantum yield measured with imaging Pulse-Amplitude-Modulation. The blue color indicates values of Fv/Fm around 0.75 close to the optimal value of 0.8, turquoise indicates a lower value around 0.45. Relative *PotriNRAMP3*.*1* and *PotriNRAMP3*.*2* mRNA levels of over-expressing lines are shown in supplementary figure S5.

**Fig. 8.**
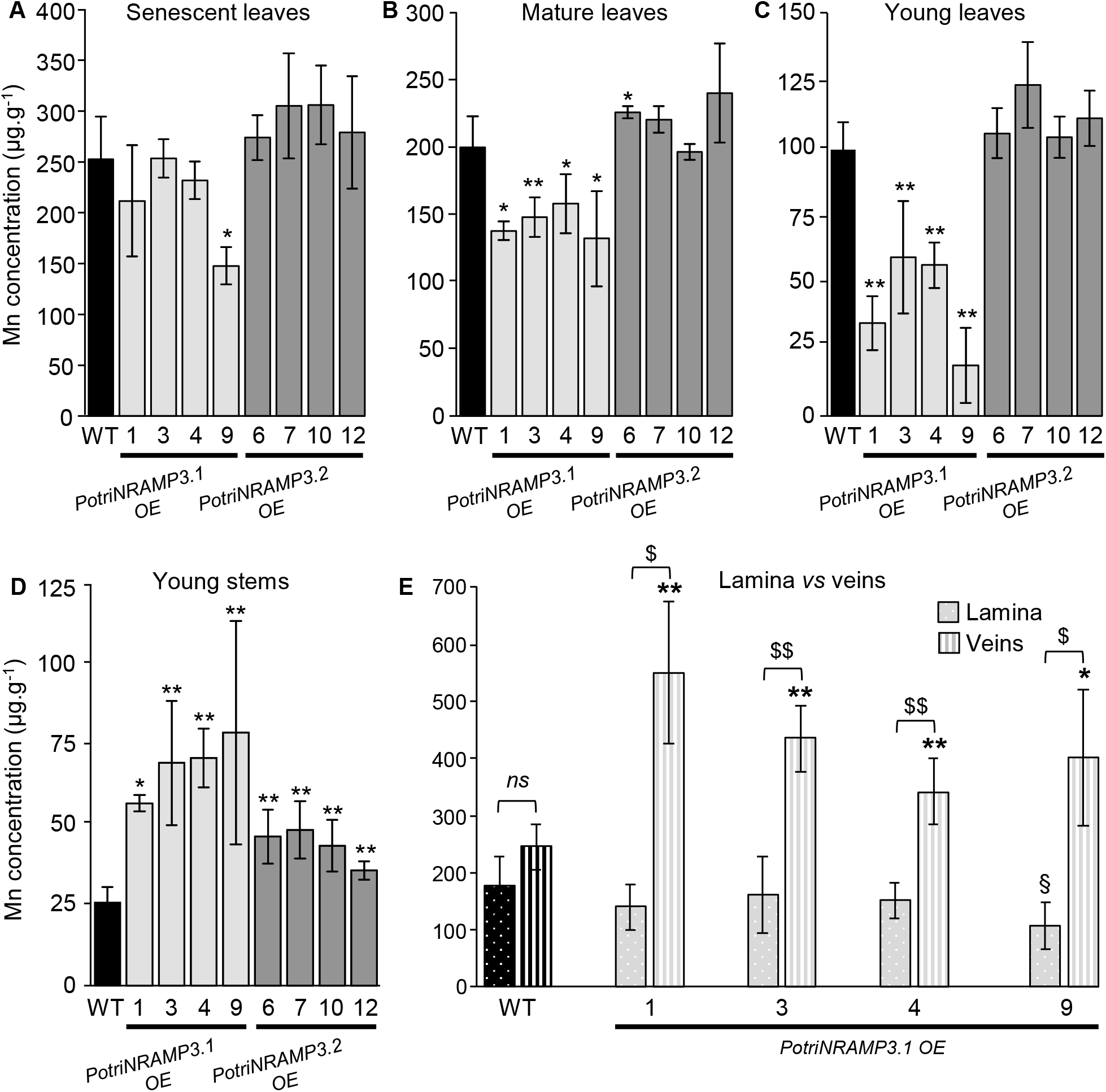
Ectopic over-expression of *PotriNRAMP3*.*1-GFP* but not that of *PotriNRAMP3*.*2-GFP* perturbs Mn distribution in poplar leaves. Mn concentrations in senescent (A), mature (B), young leaves (C) and young stems (D) of poplars 2 months after transfer from *in vitro* to soil were determined using Atomic Emission Spectroscopy. Mean Mn concentrations of PotriNRAMP3.1-GFP or PotriNRAMP3.2-GFP OE lines were compared with NT control. Error bars represent SD (n = 4-7 plants from each independent transgenic line). (E) Mean Mn concentrations in veins and lamina of mature leaves (B) from NT control and 4 *PotriNRAMP3*.*1-GFP* OE lines. Error bars represent SD (n = 4-8 leaves from 2-8 plants for each independent transgenic line). Symbols denote significant differences with the NT control (* or ^§^) or between lamina and veins (^$^) according to a Mann-Whitney test (*/^§^/^$^: *p < 0*.*05*, **/^§§^/^$$^: *p < 0*.*01*).

To further test this hypothesis, we analyzed Mn distribution in mature leaves from PotriNRAMP3.1 OE lines compared to NT control. We dissected leaves into vein and lamina, and measured Mn separately (fig. 8E). In PotriNRAMP3.1 OE lines, the Mn concentration was lower in lamina compared with veins, whereas the concentrations in these two parts of the leaf were similar in NT control. Moreover, the Mn concentration in the lamina tended to be lower in PotriNRAMP3.1 OE lines than in the NT control line, while the opposite was observed in veins. The decrease in Mn concentration in the lamina of PotriNRAMP3.1 OE lines compared to NT control was only significant for line 9, which displayed the most severe internerval chlorosis. These results confirm that over-expression of *PotriNRAMP3*.*1* in poplar perturbs Mn distribution between and within organs. Analysis of PotriNRAMP3.1 expression using RT-qPCR showed that this gene is expressed at similar levels in the lamina and the veins (fig. S14). Together, the results presented indicate that PotriNRAMP3.1 expression can modulate Mn transport between organs and tissues.

To confirm that the leaf chlorosis symptoms observed in PotriNRAMP3.1 OE lines were due to Mn depletion in the lamina, we supplemented the trees with Mn. We grew tree cuttings for 4 weeks and then started watering half of them with 0.5 mM of MnSO_4_ for an additional 5 weeks. We observed that chlorosis did not appear in newly formed leaves of Mn treated trees (fig. S15A). In these leaves, Mn concentrations were higher in lamina and veins, but the treatment did not restore the defect in Mn distribution between these tissues (fig. S15B). Chlorosis was not reverted in leaves formed prior to the treatment indicating that the treatment prevented chlorosis in newly formed leaves rather than corrected it in older leaves.

## Discussion

In this study, we have characterized two poplar NRAMP3 metal transporters using a combination of phylogenetic, cell biology and molecular genetic approaches. We found that poplar genomes harbor two tandem copies of *NRAMP3* gene under selection, whereas only one copy is present in the closest genus, *Salix*. Moreover, we demonstrated that the two paralogues encode functional metal transporters but that their functions *in planta* have diverged. Whereas PotriNRAMP3.2 has the same function in metal retrieval from the vacuole as AtNRAMP3 and AtNRAMP4, PotriNRAMP3.1 displays a distinct subcellular localization to the TGN as well as a distinct function. Our results suggest that PotriNRAMP3.1 could be involved in Mn distribution in poplar aerial organs. Elemental analyses show that poplar lines ectopically expressing *PotriNRAMP3*.*1* are impaired in Mn transfer from the stem to the leaves and, within the leaves, from the veins to the lamina, resulting in chlorosis and a decreased PS II efficiency. Together our results show that a gene duplication of NRAMP3 specific to the poplar genus gave rise to the neofunctionalization of one of the copies, while the other retained the conserved function described in other species and highlight an unsuspected role of the secretory pathway in cell-to-cell transport of Mn.

### Distinct mechanisms for the formation of *NRAMP* gene pairs in *A. thaliana* and poplar

Duplication events are a driving force in evolution, facilitating adaptation to changing environments. Although gene duplication may be followed by accumulation of deleterious mutations and gene elimination, it may also lead to diversification of gene function and sub-or neo-functionalization (Yang et al. 2006). The ancestral angiosperm genome contained only 14000 genes or less (Proost et al. 2011). However, whole genome triplications that occurred about 120 Ma significantly increased the size of the Eudicotyledon genomes.

In *A. thaliana AtNRAMP3* and *AtNRAMP4* encode functionally redundant metal transporters (Lanquar et al. 2005). This pair of genes located on two different chromosomes is present in *A. thaliana, Arabidopsis lyrata* and *Noccea caerulescens* (Oomen et al. 2009), but only one gene is found in *Carica papaya* and *Ricinus communis* genomes. Thus, they probably originate from a duplication that took place after the *Carica papaya* divergence that happened 72 Ma. The analysis of the duplicated regions of the *A. thaliana* genome showed that *AtNRAMP*3 and *AtNRAMP*4 loci are located on the duplicated block 0204146800380, suggesting that the pair originates from one of the two whole genome duplications that occurred between 70 and 23 Ma in the *A. thaliana* lineage (Lanquar et al. 2005; Ming et al. 2008; Proost et al. 2011).

The poplar lineage has also undergone one whole genome duplication 60-65 Ma *i*.*e*., before the *Populus* and *Salix* divergence that took place 52 Ma (Tuskan et al. 2006; Hou et al. 2016). Comparing gene order in *S. purpurea* and in *P. trichocarpa* showed genomic collinearity upstream and downstream *NRAMP3 loci*, except that only one copy of *NRAMP3* is found in *S. purpurea* (fig. 1; fig. S2). In contrast, two copies of *NRAMP3* were found in all sequenced poplar genotypes (fig. 2). It is unlikely that the whole genome duplication accounts for the emergence of *NRAMP3*.*1* and *NRAMP3*.*2* genes specific to *Populus* species since this event happened before the *Populus* and *Salix* divergence. Moreover, gene tandem arrangements generally imply local duplication processes rather than whole genome duplications. The genomic sequence surrounding *Populus NRAMP3*.*1* and *NRAMP3*.*2* shows homologies with Class I long terminal repeats (LTR) retrotransposon elements (Gypsy) mainly located between the two genes. Retrotransposons can mediate gene duplications. However, such duplications usually create a typical intron-free copy, which can be integrated throughout the genome and not specifically close to the initial copy (Freeling 2009). The conservation of intron/exon structure and the tandem arrangement of *Populus NRAMP3*.*1* and *NRAMP3*.*2* suggest another mechanism. Repeated sequences of retrotransposons are known to stimulate intrachromosomal recombination events or unequal crossing over, leading to gene duplication (White et al. 1994; Flagel and Wendel 2009). The genome of poplar which contains significantly more gene tandems than that of *A. thaliana* (Proost et al. 2011), contains also three time more transposons (Ming et al. 2008). Thus, it is most likely that this mechanism accounts for the tandem duplication of *Populus NRAMP3*.*1* and *NRAMP3*.*2*. Therefore, distinct mechanisms of gene duplication led to *NRAMP* gene pair formation in poplar and *A. thaliana*.

### *Populus NRAMP3* copies are subjected to both positive and purifying selection

Non-synonymous (dN) vs synonymous codons (dS) analyses (dN/dS) highlight that *Populus NRAMP3*.*1* and *NRAMP3*.*2* sequences are mostly under purifying selection (fig. 3 and S3; tables S2 and S3) acting on many residues in the core conserved transmembrane domains of the protein. This is in agreement with the finding that both PotriNRAMP3.1 and PotriNRAMP3.2 have retained metal transport ability. However, the evidence suggests both positive and relaxed purifying selection especially on residues localized at the N and C terminal ends of the protein (fig. 3; fig. S3, tables S2 and S3). Mutations in the N terminal region, where motives involved in the targeting of AtNRAMP3 and AtNRAMP4 have previously been identified, likely enabled the protein to acquire distinct subcellular localizations (Müdsam et al. 2018). The vacuolar membrane localization of PotriNRAMP3.2 (fig. 6A), its transport capacities (fig. 4; fig. S6 and S7) and its ability to complement *A. thaliana nramp3nramp4* double mutant phenotypes (fig. 5) indicate that PotriNRAMP3.2 is the functional homologue of AtNRAMP3 and AtNRAMP4. The ability of PotriNRAMP3.1 to transport Fe and Mn in yeast (fig. 4; fig. S6 and S7), its distinct intracellular localization (fig. 6), its inability to complement the *nramp3nramp4* double mutant phenotypes (fig. 5) and the signature of purifying selection (fig. 3, fig. S3, tables S2 and S3) argue in favor of neo-functionalization rather than non-functionalization. The finding that *Populus NRAMP3*.*1* and *Populus NRAMP3*.*2* promoter sequences lack significant sequence identities further suggests that the regulation of the two copies has diverged (fig. S1D). This is in agreement with our previous report showing that *Populus NRAMP3*.*1* and *NRAMP3*.*2* belong to different networks of co-expressed genes (Pottier, Garcia de la Torre, et al. 2015). We propose a scenario in which after *Populus NRAMP3* gene duplication, relaxation of purifying selection allowed mutations altering PotriNRAMP3.1 subcellular localization. This mutated version of *Populus NRAMP3* was subsequently maintained by purifying selection, probably because it conferred improved fitness. Specific inactivation of *Populus NRAMP3*.*1* would allow to further test this scenario.

### PotriNRAMP3.1 modulates tissue distribution of Mn

To investigate the function of PotriNRAMP3.1 and PotriNRAMP3.2 *in planta*, we generated transgenic poplars over-expressing *PotriNRAMP3*.*1-GFP* and *PotriNRAMP3*.*2-GFP*. PotriNRAMP3.1 and PotriNRAMP3.2 subcellular localizations in poplar roots or leaves were similar to those observed in *A. thaliana*. (fig. 6; fig. S10). Analysis of metal concentrations revealed a specific decrease in Mn concentrations in leaves from lines over-expressing *PotriNRAMP3*.*1-GFP*. Moreover, lines with the lowest Mn leaf concentrations displayed internerval chlorosis (fig. 7). In agreement with the role of Mn in photosynthesis, PS II efficiency was decreased in the chlorotic parts of *PotriNRAMP3*.*1* OE leaves (fig. 7). Further analysis showed that the decrease in Mn leaf concentration was associated to an increase in Mn concentration in stems (fig. 8). Furthermore, Mn distribution within leaves was also affected in *PotriNRAMP3*.*1* OE lines: Mn accumulated at higher levels in the veins while it was depleted in the lamina. Together, these results indicate that PotriNRAMP3.1 can modulate transcellular transport of Mn. This phenotype is unexpected as PotriNRAMP3.1-GFP exhibits a clear intracellular localization to the TGN. Even though the partial complementation of *nramp3nramp4* suggests that a small fraction of PotriNRAMP3.1-GFP might be targeted to the vacuolar membrane (fig. S9), the absence of similar phenotypes in PotriNRAMP3.2-GFP OE lines support the hypothesis that the fraction of PotriNRAMP3.1 associated to the TGN is responsible for the phenotypes observed in PotriNRAMP3.1 OE lines. To account for this phenotype, we propose a working model (fig. 9) based on two hypotheses (1) Mn cell-to-cell transport requires Mn secretion (2) expression of *PotriNRAMP3*.*1* limits Mn secretion by allowing the retrieval of Mn from the TGN, which would otherwise be secreted. PtNRAMP3.1 overexpression seems to restrict specifically Mn transfer from the veins to the lamina. However, PotriNRAMP3.1 is expressed at similar levels in the veins and in the lamina (Fig S13). Mn secretion to the cell wall is expected to be important for cell to cell transfer when cells are not or poorly connected by plasmodesmata. We propose that the effect of PotriNRAMP3.1 expression is more pronounced in the lamina because (1) veins are directly supplied with Mn upon unloading from the xylem sap, and (2) the density of plasmodesmata between lamina cells is lower than between the cells of the veins (Russin and Evert 1985).

**Fig. 9.**
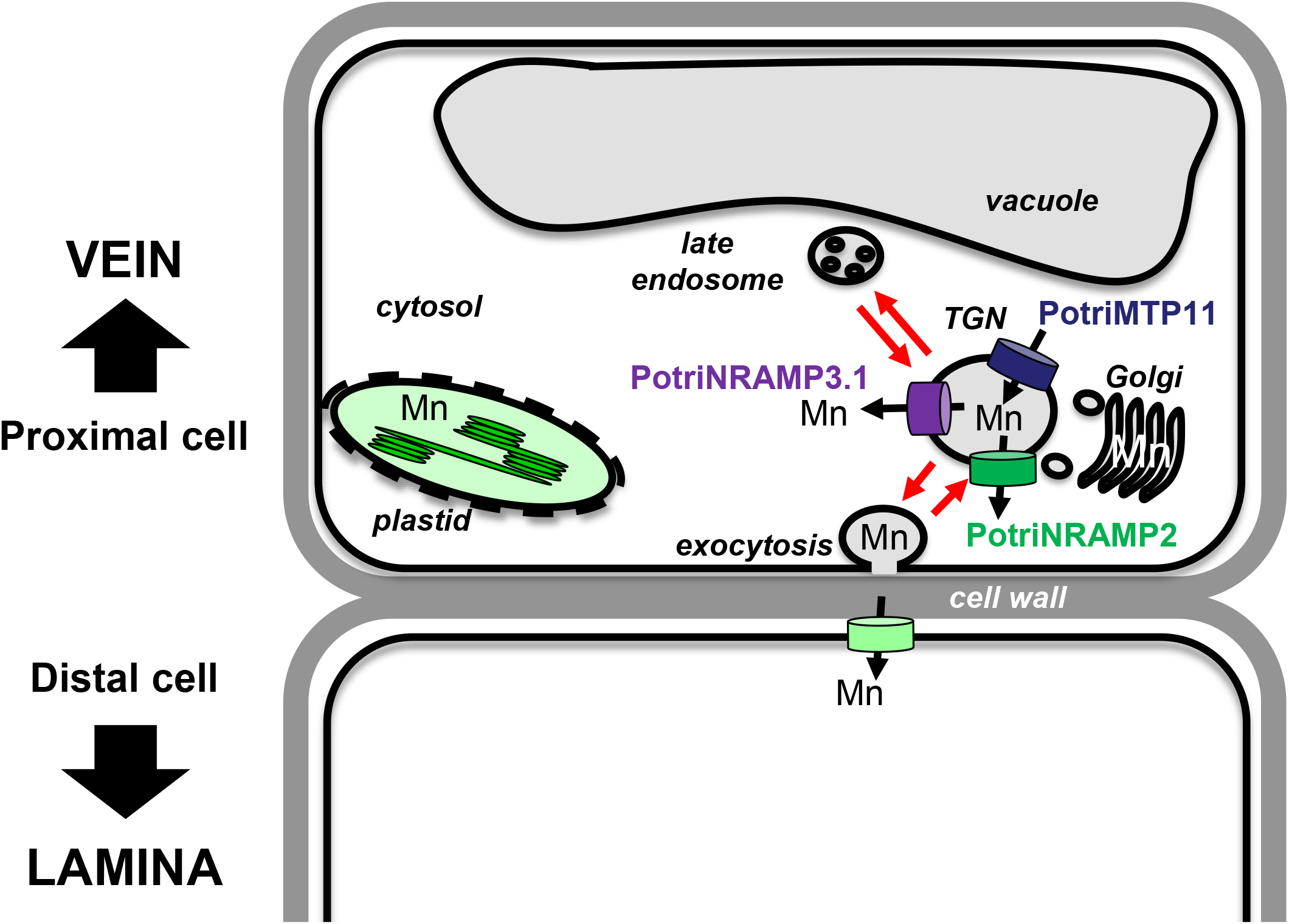
Working model to account of PotriNRAMP3.1 role in Mn transport from cell to cell. The model is based on the hypothesis that Mn moves through the transcellular pathway, being secreted in the apoplast via exocytosis by the cells proximal to the veins and taken up by the cells that are distal to the veins. According to this hypothesis, the transporters loading Mn (PotriMTP11) or unloading Mn (PotriNRAMP3.1 as well as PotriNRAMP2 assuming function conservation with Arabidopsis AtNRAMP2) from the secretory pathway would determine the amount of Mn made available by proximal cells for uptake by distal cells. In this context, efficient removal of Mn from the secretory pathway in the proximal cells by PotriNRAMP3.1 over-expression would limit Mn availability for the distal cell.

The function of PotriNRAMP3.1 in retrieval of Mn from the secretory system would be equivalent to the function of AtNRAMP2 (Alejandro et al. 2017; Gao et al. 2018). AtNRAMP2 localizes to the TGN and was proposed to retrieve Mn from this compartment to make it available for uptake into chloroplasts. Previous work also showed that loss of AtMTP11, which also localizes in the secretory system, leads to an increase in plant Mn concentration (Peiter et al. 2007). As AtMTP11 is involved in Mn loading into the secretory system, the observed phenotype also agrees with the hypothesis that Mn concentrations in plant tissues are, at least in part, controlled by Mn secretion. As SMF2 has also been proposed to retrieve Mn from the secretory system in yeast, the model of figure 9 would also account for the strong decrease in Mn content in the *smf2* mutant (Luk and Culotta 2001). Homologues of AtMTP11 and AtNRAMP2 are also present in poplar and could act in concert with PotriNRAMP3.1 to control the rate of Mn secretion vs intracellular distribution in poplar cells. Interestingly, mRNA levels of *PotriNRAMP2* and *PotriNRAMP3*.*1* are correlated in leaves (Pottier, García de la Torre, et al. 2015).

Taken together, our results provide a clear case for neofunctionalization in a tandem of NRAMP genes specific to poplar. It also provides new insights into the cell-to-cell transport of divalent cations by showing the significant contribution of the secretory pathway in the cellular export of Mn as well as new evidence of the essential role of transporters located at the secretory pathway in the regulation of the cellular storage of Mn. In the future, it would be interesting to find out what is the advantage conferred by the newly functionalized PotriNRAMP3.1 that led to the conservation of this gene in all examined poplar species. It will also be interesting to understand the interplay between PotriNRAMP3.1, PotriNRAMP2 and PotriMTP11 in the control of Mn concentration in the secretory pathway.

## Materials and methods

### Sequences analysis

Sequences were retrieved as indicated in the table S1. To obtain the homologous genomic sequence of *PotriNRAMP3*.*1* (Potri007G050600) and *PotriNRAMP3*.*2* (Potri007G050600) from non-assembled poplar and willow genomes, raw reads were aligned to *P. trichocarpa* genome V4.1. *PotriNRAMP3*.*1* and *PotriNRAMP3*.*2* were blasted on the obtained genome consensus using QIAGEN CLC Genomics Workbench 12.0. The best hits were then blasted back on *P. trichocarpa* genome V4.1 to confirm the sequence homology relationship. *P. trichocarpa* and *S. purpurea* genomic DNA homologies were investigated by Dot-plot analyses using Gepard softwares V1.30 and V1.40 (Krumsiek et al. 2007).

### Phylogenetic tree construction

The tree shown in figure 2 was generated from amino acid sequences listed in supplementary data S3 and outgroup sequences listed in the figure legend. For figure S1, the accession numbers of the sequences are provided in the figure legend. Full-length sequences were imported into the Molecular Evolutionary Genetics Analysis (MEGA) package version 7, and aligned by CLUSTALW (Kumar et al. 2016). Only conserved positions were used for figure 2, while all positions with at least 90% site coverage was used for figure S1. Phylogenetic analyses were conducted using the Maximum Likelihood method. Thanks to the “Find best protein model (ML)” tool available in MEGA 7, the lower BIC (Bayesian Information Criterion) model was selected for each tree. Therefore, JTT matrix-based model and the Le_Gascuel (LG) model were used for the trees displayed in figure 2 and figure S1, respectively (Jones et al. 1992; Le and Gascuel 2008). A discrete Gamma distribution was used to model evolutionary rate differences among sites (5 categories, +*G*, parameter = 0.55 and parameter = 1.15 for figure 2 and figure S1, respectively). Initial tree(s) for the heuristic search were obtained automatically by applying Neighbor-Joining and BioNJ algorithms to a matrix of pairwise distances estimated using a JTT model, and then selecting the topology with superior log likelihood value. The bootstrap consensus tree inferred from 1000 replicates was taken to represent the evolutionary history of the analyzed genes. Branches corresponding to partitions reproduced in less than 50% bootstrap replicates were collapsed. Trees are drawn to scale, with branch lengths measured in the number of substitutions per site. To determine the codon specific selective pressure by FEL method (see “selective pressure analysis”), a nucleotide substitution-based tree has also been generated. The best-fitting nucleotide model was selected by iterative procedure as described (Kosakovsky Pond and Frost 2005a), and initial estimate of the phylogeny was reconstructed by neighbor-joining (Saitou and Nei 1987) using the Tamura-Nei distance (Tamura and Nei 1993; Kosakovsky Pond and Frost 2005b).

### Selective pressure analysis

After codon based alignment, global dN/dS was first calculated for each pair of the 26 poplar NRAMP CDS using HYPHY in MEGA7. Then, the distribution of dN/dS along the protein sequence was computed through the Neij Gojobori algorithm using a 20 residue window with a shift of 10 residues using JCoDA 1.4 (Nei and Gojobori 1986; Steinway et al. 2010). Finally, positive and purifying selection at individual sites were inferred using the FEL method available at https://www.datamonkey.org (Kosakovsky Pond and Frost 2005b; Weaver et al. 2018). FEL generated a phylogenetic tree with the 26 Populus NRAMP CDS. A subset of branches encompassing either the *NRAMP3*.*1s* or the *NRAMP3*.*2*s were analyzed separately to estimate dS and dN at a site (α and β, respectively). A maximum-likelihood approach was then undertaken to calculate the dN/dS for each codon site (L; p < 0.05).

### Construction of expression vectors

*PotriNRAMP3*.*1* and *PotriNRAMP3*.*2* CDSs were amplified by PCR from cDNA synthesized from leaf RNA of *P. trichocarpa* cv Nisqually-1 using the Phusion high-fidelity DNA polymerase (Thermo-Scientific) and primers listed in table S4. The gel-purified PCR products were recombined into pDONR207 for *PotriNRAMP3*.*1* and into pDONR201 for *PotriNRAMP3*.*2* following the BP Clonase (Invitrogen) manufacturer’s instruction. LR reactions were performed using the pDR195gtw vector (Rentsch et al. 1995; Oomen et al. 2009) for the generation of yeast expression vectors, and using pB7FWG2 (Karimi et al. 2002), pMDC83 (Curtis and Grossniklaus 2003) and pUB-DEST binary vectors (Grefen et al. 2010) for the generation of plant expression vectors.

### Yeast growth assays

Yeasts were transformed as indicated in supplementary materials and methods. Transformed *smf1* and *smf2* yeast mutants were grown overnight in liquid Synthetic Dextrose -ura (SD -ura, pH 6). The cultures were diluted to ODs of 1 to 10^−3^ and spotted on SD -ura plates (pH 6). Transformed *smf1* and *smf2* strains were spotted on SD -ura, supplemented with 5 mM (*smf1*) or 10 mM (*smf2*) ethylene glycol*bis* (beta-aminoethyl ether)-*N,N,N*′,*N*′-tetraacetic acid (EGTA) and 100 µM MnSO_4_ (+ Mn) or with 5 mM (*smf1*) or 10 mM (*smf2*) EGTA without MnSO_4_ (-Mn).

### Confocal imaging

Roots of *in vitro* grown 6-day-old *A. thaliana* seedlings or 2-3 week-old poplar explants were mounted in liquid culture medium, and confocal images of epidermal cells in the elongation zone were obtained by high-speed (100 ms) sequential acquisition of GFP (λex = 490 nm, λem = 500–550 nm) and mRFP (λex = 590 nm, λem = 600–650 nm, employing a Nipkow spinning disk confocal system equipped with a Prime 95Bcamera (Photometrics) and a Nikon 100X 1.4 aperture oil immersion objective. Super resolution images were generated with a Live-SR module for optically demodulated structured illumination (GATACA Systems). Image processing (cropping, contrast adjustment and background subtraction) was performed with ImageJ 1.45s program (Schneider et al. 2012).

### Plant material and plant transformation

The generation of the *nramp3nramp4* double mutants of *A. thaliana* Col-0 was described previously (Bastow et al. 2018). pUB-DEST and pB7FWG2 constructs were introduced in *nramp3nramp4* mutants through *Agrobacterium tumefaciens* (strain AGL0) mediated transformation using the flower dip method (Clough and Bent 1998). Independent homozygous *A. thaliana* Col-0 *nramp3nramp4* transformants with a single insertion locus were obtained by plant selection based on Basta resistance. The poplar INRA 717-1-B4 clone (*P. tremula* x *P. alba*) was transformed as described in supplementary materials and methods using media listed in table S5 (Leplé et al. 1992).

### A. thaliana growth conditions

*A. thaliana* seedlings were grown on ABIS medium plates containing 2.5 mM H_3_PO_4_, 5 mM KNO_3_, 2 mM MgSO_4_, 1 mM Ca(NO_3_)_2_, MS microelements, 1% sucrose, 1% Phytagel, 1 mM MES adjusted with KOH to pH 6.1 and FeHBED (Strem Chemicals, Newburyport, MA, USA) as indicated in figure legends. FeHBED was prepared as described by Lanquar *et al*. (2005). For low Fe sensitivity growth assays, plants were grown for 8 days on plates where Fe was omitted. Plates were placed vertically in environmental growth chambers (Sanyo MLR-350, Morigushi, Japan) at 21°C with a 16 h photoperiod under 120 µmol photon·m^-2^·s^-1^.

### Photosystem II maximum quantum yield

Phostosystem II maximum quantum yield was determined using an Imaging PAM (Walz, Germany). Efficiency of the photosynthetic electron transport (Fv/Fm) was assayed by calculating the ratio of variable fluorescence (Fv) to maximal fluorescence (Fm) after a saturating light pulse (Maxwell and Johnson 2000). Plants were dark adapted for 15 min prior to measurements. The fluorescence was measured under low measuring light (F_0_) and after a flash of saturating light (Fm). Fv/Fm was calculated as (Fm-F_0_)/Fm. Five areas of interest (AOI) of each leaf were selected for quantification in three leaves from different individuals of each genotype.

### Root length measurements

Plants grown vertically on plates were photographed at the indicated times and root length was determined using ImageJ and a digitizer tablet (Schneider et al. 2012; Intuos 4M WACOM, Krefeld, Germany).

### Elemental analysis

For metal analyses in yeast, liquid SD -ura medium containing transformed *smf2* strain growing overnight were diluted to OD 0.3 in liquid SD -ura supplemented with 30 µM FeCl_3_ and 10 µM MnSO_4_. After 30 h of incubation at 30°C under agitation, yeast cells were recovered by centrifugation (3340*g*, 5 min, 4°C) and washed twice in 50 ml ice cold YNB supplemented with EDTA 20 mM and MES 50 mM pH6, pelleted and then washed in ice cold ultrapure water. For metal analyses in plants, tissues were harvested and washed. The dry weight (DW) of the samples (yeasts or plants) was measured after drying at 60°C for 3 days. Dried samples were mineralized and analyzed for metal content as previously described (Pottier et al. 2019).

### Statistical analysis

Data were analyzed with Kruskal-Wallis and Mann-Whitney non-parametric tests for multiple comparisons and pair comparisons, respectively. For multiple comparisons, a Dunn’s post hoc test was performed when significant differences were detected. Both tests were performed using GraphPad Prism 7.

## Acknowledgments

This work was supported by grants from the DIM Astréa (Région Ile-de-France) to MP, from the VIED organisation to VALT, ANR PHYTOPOP (ANR-06-ECOT-0015) and MOBIFER (ANR-17-CE20-0008-02) to ST. CPB was funded by AgriObtentions. The ST team is supported by the CNRS and benefits and from the support of Saclay Plant Sciences-SPS (ANR-17-EUR-0007). This work has benefited from the facilities and the expertise of Imagerie-Gif microscopy platform, which is supported by France-BioImaging (ANR-INBS-04, ‘Investments for the future’). The authors thank Prof. Wolf B. Frommer for providing support.

## Data availability statement

The data underlying this article are available in the article and in its online supplementary material.

