## Supplementary Figures for "Duplication of *NRAMP3* gene in poplars generated two homologous transporters with distinct functions"

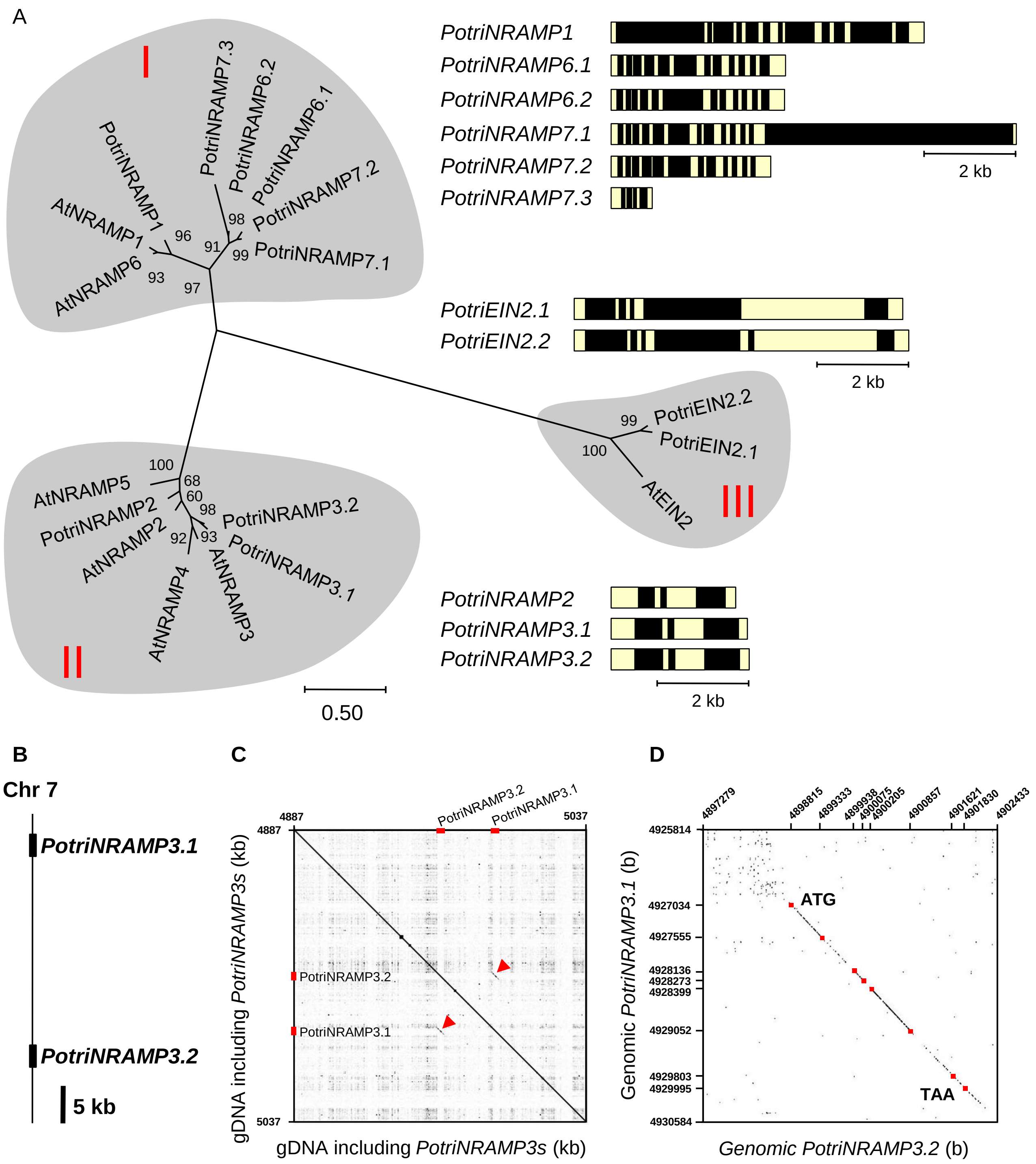


**Fig. S1**. Phylogeny, genomic structure and duplication among *PotriNRAMP* genes. (A) Phylogeny and genomic structure of the NRAMP family genes in *P. trichocarpa*. The intron (black) -exon (yellow) structure of the *PotriNRAMPs* is represented beside the phylogenetic tree. Phylogenetic analyses were conducted as described in Materials and methods with the protein sequences corresponding to the following accession numbers and gene models of *A. thaliana* and *P. trichocarpa*: At1G80830.1 (AtNRAMP1), At1G47240.1 (AtNRAMP2), At2G23150.1 (AtNRAMP3), At5G67330.1 (AtNRAMP4), At4G18790.1 (AtNRAMP5), At1G15960.1 (AtNRAMP6), AT5G03280 (AtEIN2), Potri.001G044900.1 (PotriNRAMP1), Potri.002G121000.1 (PotriNRAMP2), Potri007G050700.1 (PotriNRAMP3.1), Potri007G050600.1 (PotriNRAMP3.2), Potri.002G080400.1 (PotriNRAMP6.1), Potri.002G080500.1 (PotriNRAMP6.2), Potri.005G181000.1 (PotriNRAMP7.1), Potri.005G181100.1 (PotriNRAMP7.2), Potri.005G180900.2 (PotriNRAMP7.3), Potri.016G090800.4 (PotriEIN2.1), Potri.006G127100.7 (PotriEIN2.1). (B) Schematic representation of PotriNRAMP3 duplication on Chromosome 7. (C) Dot-plots performed with Gepard software V1.30 (Krumsiek *et al.,* 2007), between 150 kb of genomic DNA centred on PotriNRAMP3.1- PotriN*RAMP3.2* sequence and the same sequence. Sequence homologies are indicated by black dots. Red arrows highlight the specific duplication of *PotriNRAMP3* sequence. (D) Dot-plots between *PotriNRAMP3.1* and *PotriNRAMP3.2* genomic DNAs. Homology is restricted to the coding sequences. Red squares delimit intron and exon.

**A**

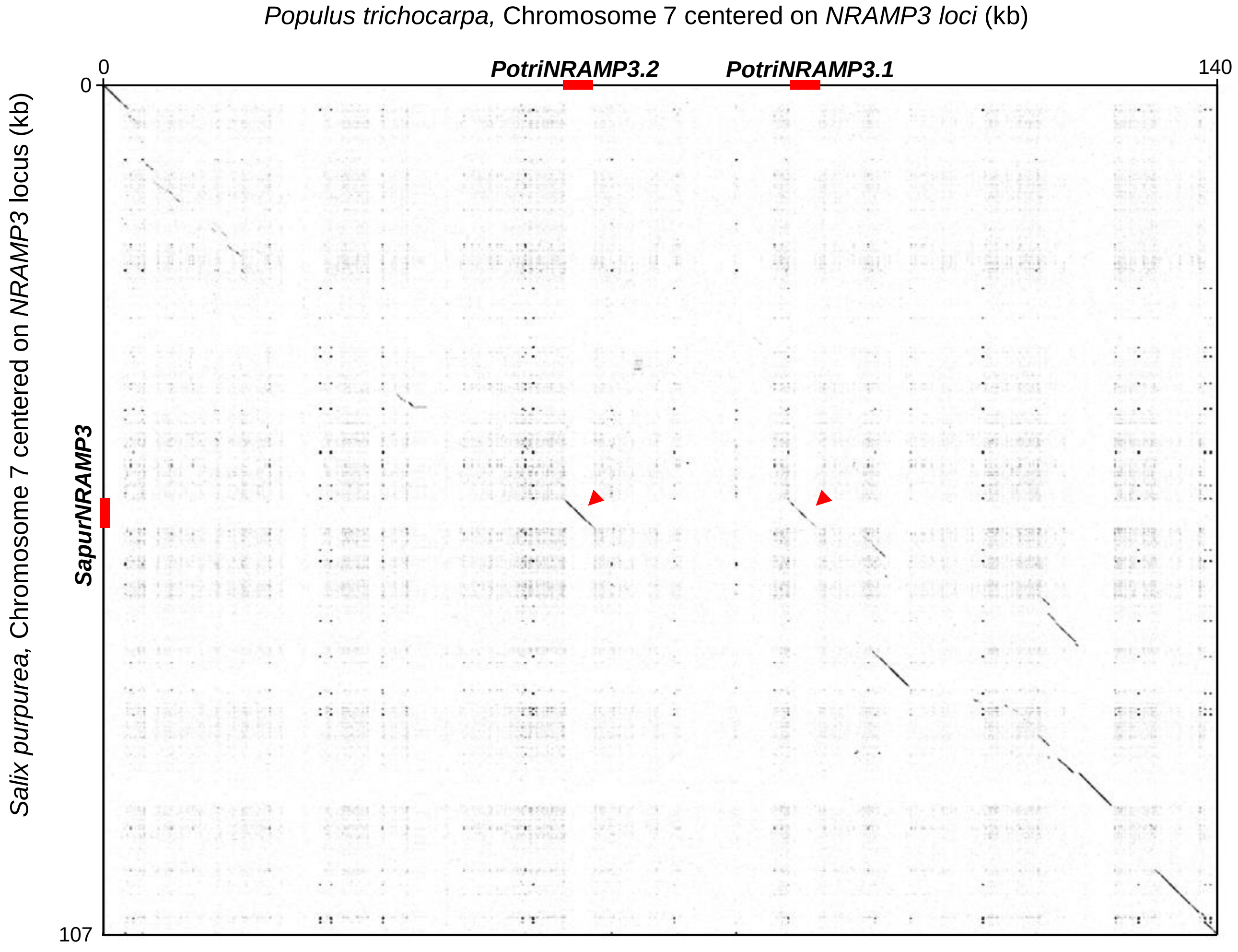


**Fig. S2.** Collinearity between *Salix purpurea* and *Populus trichocarpa* genomes in the area surrounding *NRAMP3* loci. Dot-plot centered on *NRAMP3* loci, performed on Gepard software V1.40 (Krumsiek *et al.,* 2007), using 107 kb and 140 kb sequences of *S. purpurea* and *P. trichocarpa* genomes, respectively. Sequence homology between *S. purpurea* and *P. trichocarpa* are indicated by black dots. Red arrows indicate homology between *SapurNRAMP3* and *PotriNRAMP3.2* as well as between *SapurNRAMP3* and *PotriNRAMP3.1.*

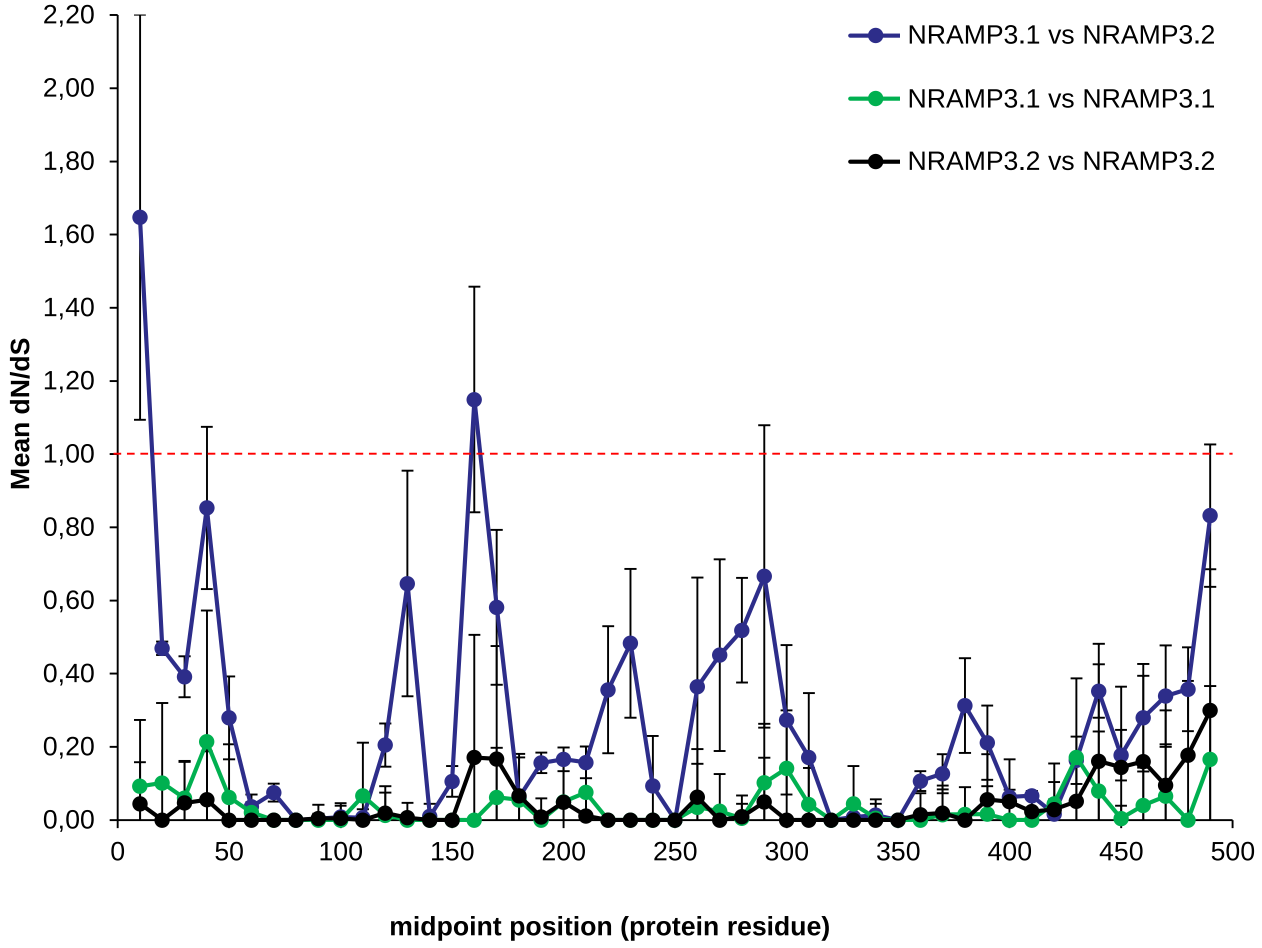


**Fig. S3.** Non-synonymous (dN) vs synonymous (dS) codon analysis along PotriNRAMP3 coding sequences. Codon alignment were performed with cDNA encoding 13 NRAMP3.1 and 13 NRAMP3.2 proteins from 13 different *Populus* species. dN/dS was calculated for each pair of cDNA (325 comparisons) through the Neij Gobodojori algorithm along the protein sequence using a 20 residue window with shift of 10 residues. Mean dN/dS for each window position was calculated independently for NRAMP3.1 vs NRAMP3.1 comparisons (78), NRAMP3.2 vs NRAMP3.2 comparisons (78) and NRAMP3.1 vs NRAMP3.2 comparisons between species (169). The mean dN/dS ± SD is shown for each window midpoint position. Analyses were conducted using JCoDA 1.4 (Steinway *et al.,* 2010).

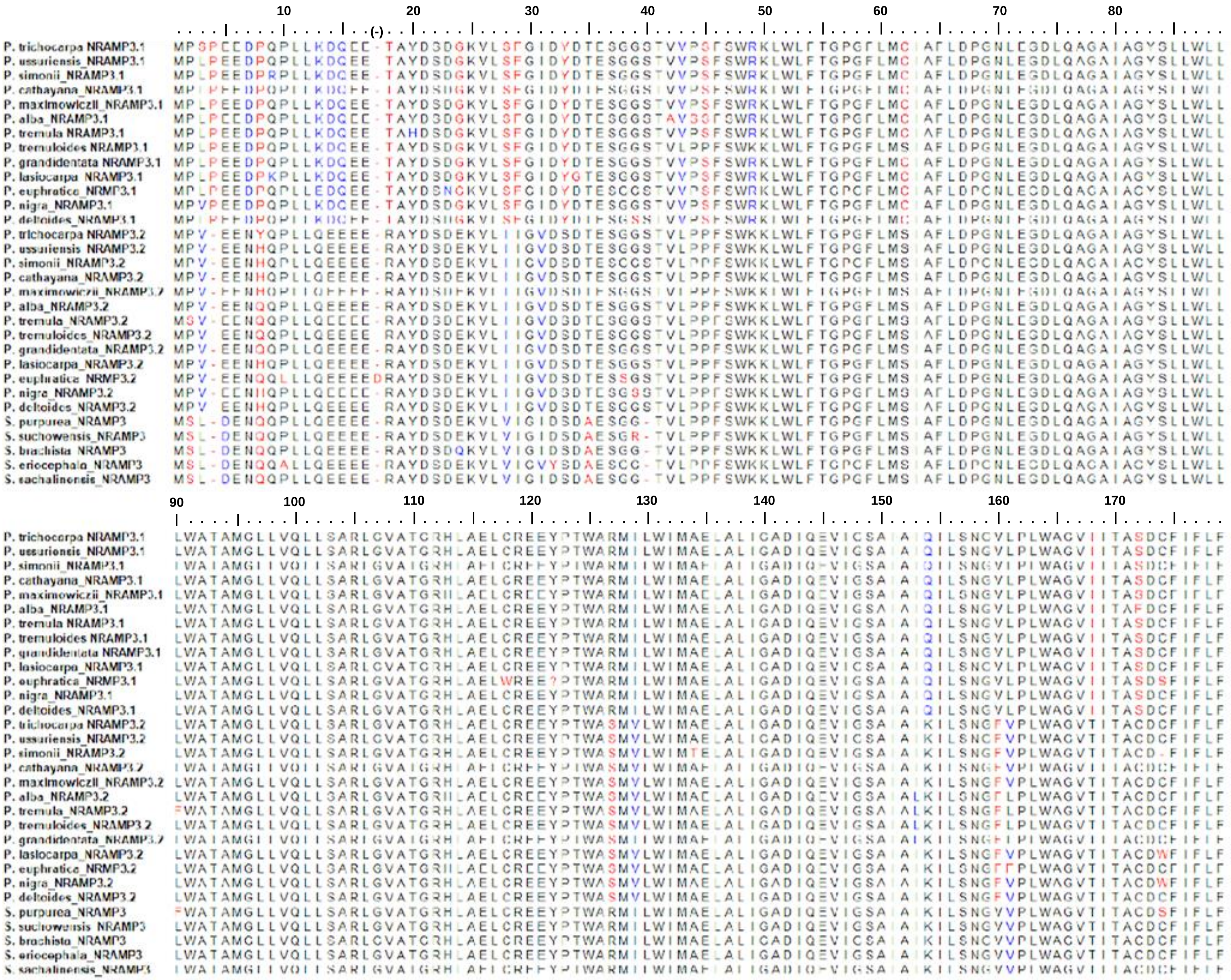

**Fig. S4.** Alignment of NRAMP3 protein sequences from *Populus* and *Salix sp*. Identical, similar and different residues between sequences are indicated in black, blue and red respectively.

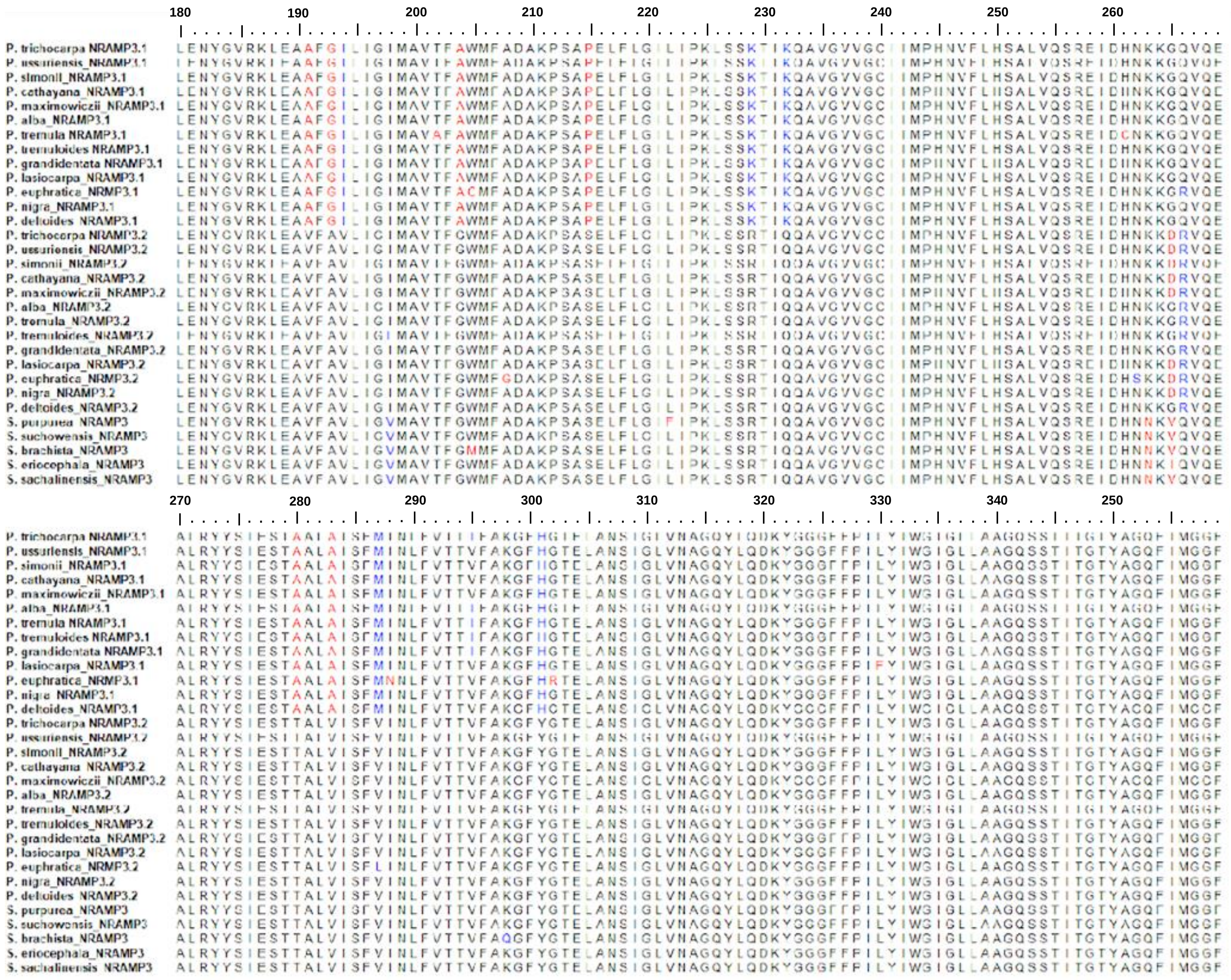

**Fig. S4 (continuation).** Alignment of NRAMP3 protein sequences from *Populus* and *Salix sp*. Identical, similar and different residues between sequences are indicated in black, blue and red respectively.

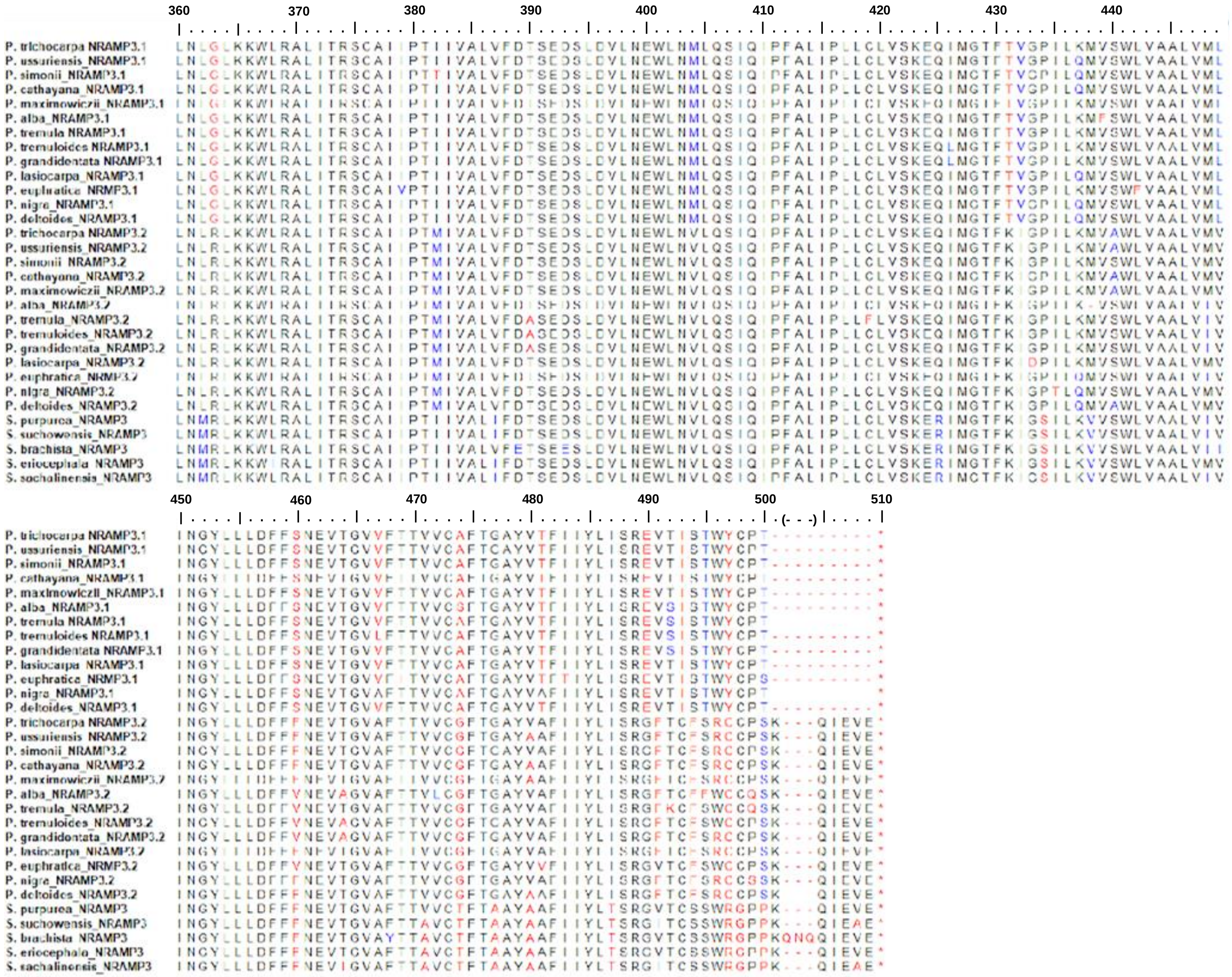

**Fig. S4 (continuation).** Alignment of NRAMP3 protein sequences from *Populus* and *Salix sp*. Identical, similar and different residues between sequences are indicated in black, blue and red respectively.

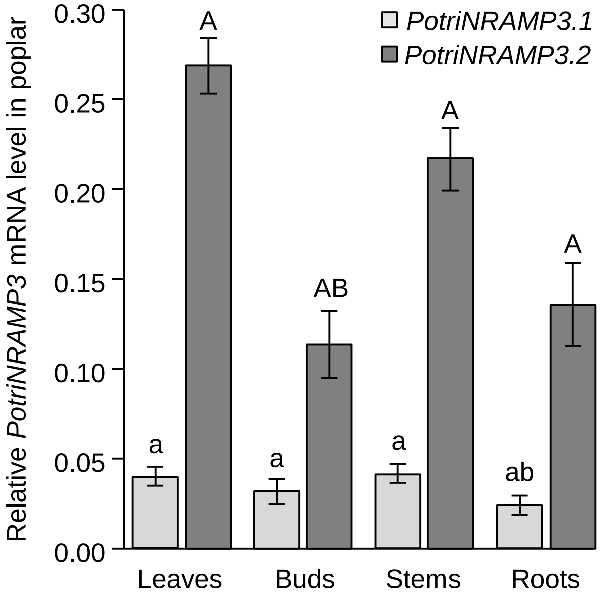


**Fig. S5.** *PotriNRAMP3.1* and *PotriNRAMP3.2* are both expressed in all organs. *PotriNRAMP3.1* and *PotriNRAMP3.2* transcript levels were determined by RT-qPCR using specific primers for each isoform and normalization by three reference genes, *i.e., UBQ*, *EF1,* and *PP2A*. The graph shows means ± SD of four replicates on different individuals. Different letters denote significant differences among organs for *PotriNRAMP3.1* (lowercase) or *PotriNRAMP3.2* (capitals) according to a Kruskal-Wallis test followed by Dunn's test for multiple comparisons ($p < 0.01$).

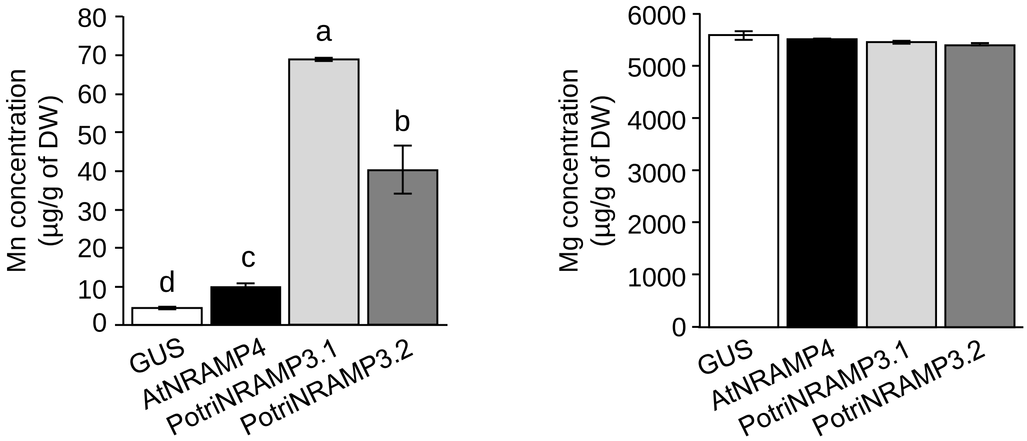


**Fig. S6.** Expression of *PotriNRAMP3.1* or *PotriNRAMP3.2* increase Mn accumulation in yeast cells. Liquid SD -ura medium containing *smf2* cells expressing *GUS*, *AtNRAMP4*, *PotriNRAMP3.1* or *PotriNRAMP3.2* were grown overnight and diluted to OD 0.3 in liquid SD -ura supplemented with 30 µM $FeCl_3$ and 10 µM $MnSO_4$. After 30 hrs of incubation at 30°C, yeast cells were washed twice, and metal concentrations were determined by atomic absorption spectrometry. Error bars represent the SD of three replicates. Different letters denote significant differences between transformed strains according to a Kruskal-Wallis test followed by Dunn's test for multiple comparisons ($p < 0.05$).

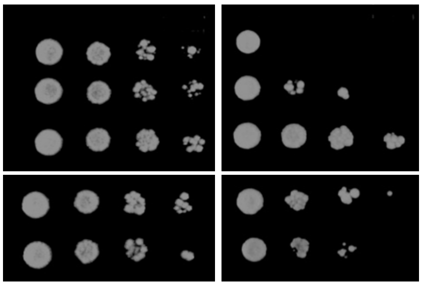


**Fig. S7.** PotriNRAMP3.1 and PotriNRAMP3.2 transport Fe. Functional complementation of the *fet3fet4* yeast iron uptake mutant by *PotriNRAMP3.1* and *PotriNRAMP3.2*. *fet3fet4* yeast cells, which are impaired in low (FET4) and high (FET3) affinity Fe uptake systems, were transformed with pDR195gtw vector containing the cDNA of *GUS, AtNRAMP1, AtNRAMP4*, *PotriNRAMP3.1* and *PotriNRAMP3.2*. AtNRAMP1 and AtNRAMP4, which were previously shown to transport Fe and rescue the *fet3fet4* yeast double mutant, were used as positive controls. GUS, which encodes the enzyme β-glucuronidase, was used as a negative control. Transformed *fet3fet4* yeast strains were grown overnight in liquid synthetic dextrose -ura supplemented with 0.2 mM $FeCl_3$. Cultures were diluted to ODs of 1 to $10^{-3}$ and spotted on synthetic dextrose -ura plates supplemented with 100 µM of the Fe chelator BathoPhenanthroline-Di-Sulfonic acid (BPDS) and 0.2 mM $FeCl_3$ (+ Fe) or with 100 µM BPDS and 40 µM $FeCl_3$ (- Fe). The plates were incubated at 30°C for 4 days before photography.

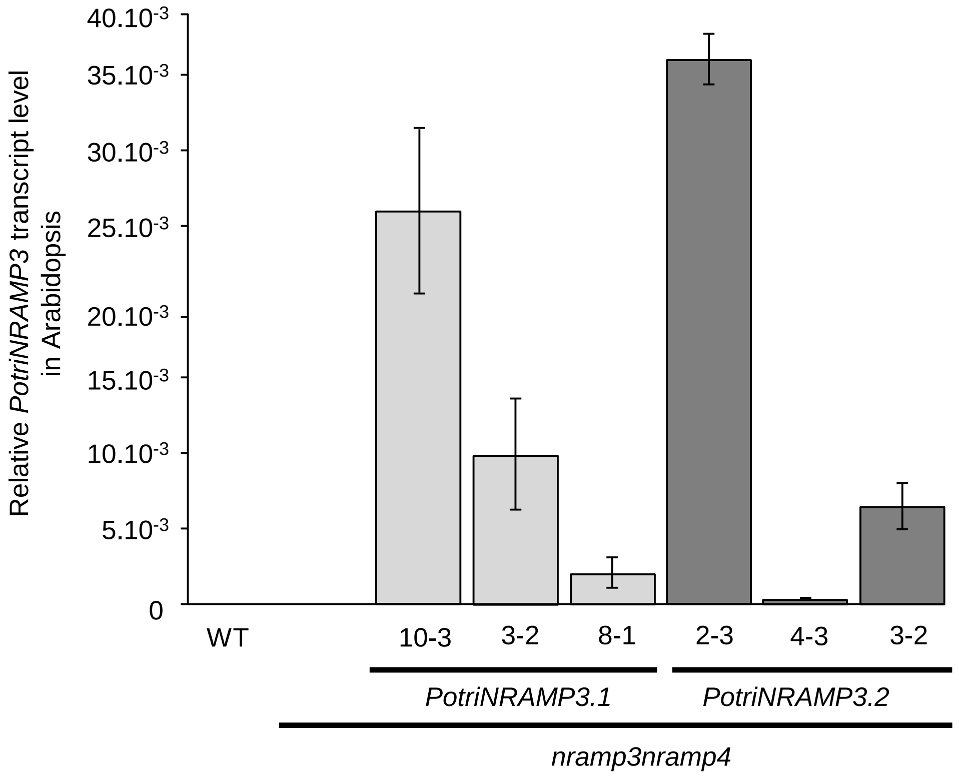


**Fig. S8.** Transcript levels of *PotriNRAMP3.1* and *PotritNRAMP3.2* in Arabidopsis *nramp3nramp4* transgenic lines. *PotriNRAMP3.1* and *PotriNRAMP3.2* transcript levels were determined by RT-qPCR using specific primers for each isoform and normalization with *AtACTIN* as a reference gene. The graph shows means ± SD of three replicates on different individuals.

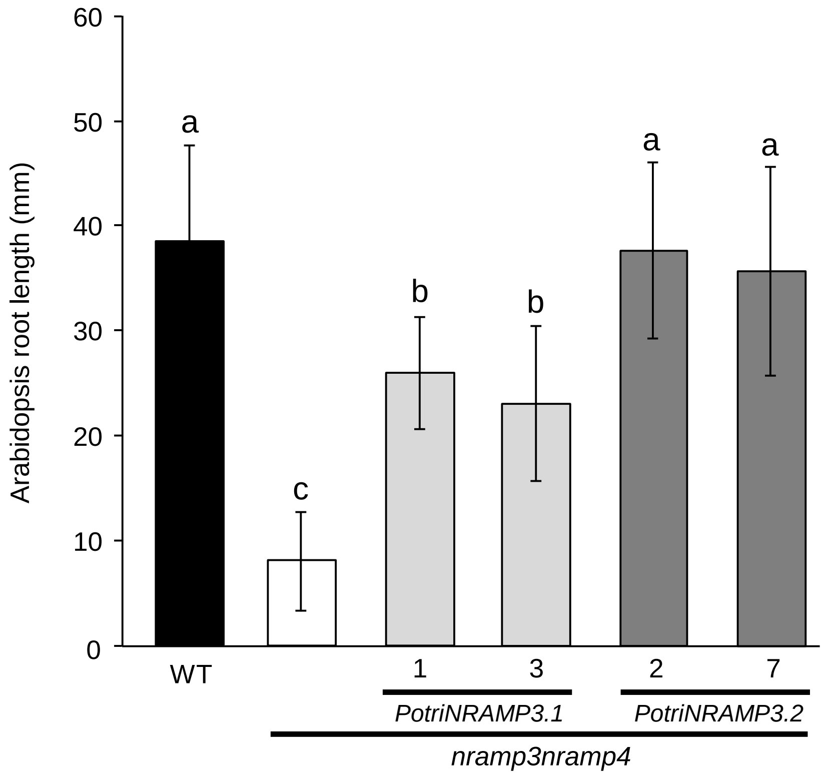


**Fig. S9.** Root length of Col-0 wild-type, *nramp3nramp4* mutant and representative transformed with *p35S::PotriNRAMP3.1-GFP* or *p35S::PotriNRAMP3.2-GFP*. Roots were measured 12 days after germination on ABIS medium without Fe. The graph shows means ± SD (n = 10-25 roots for each genotype). Different letters indicate significant difference between genotypes based on ANOVA analysis ($p < 0.05$).

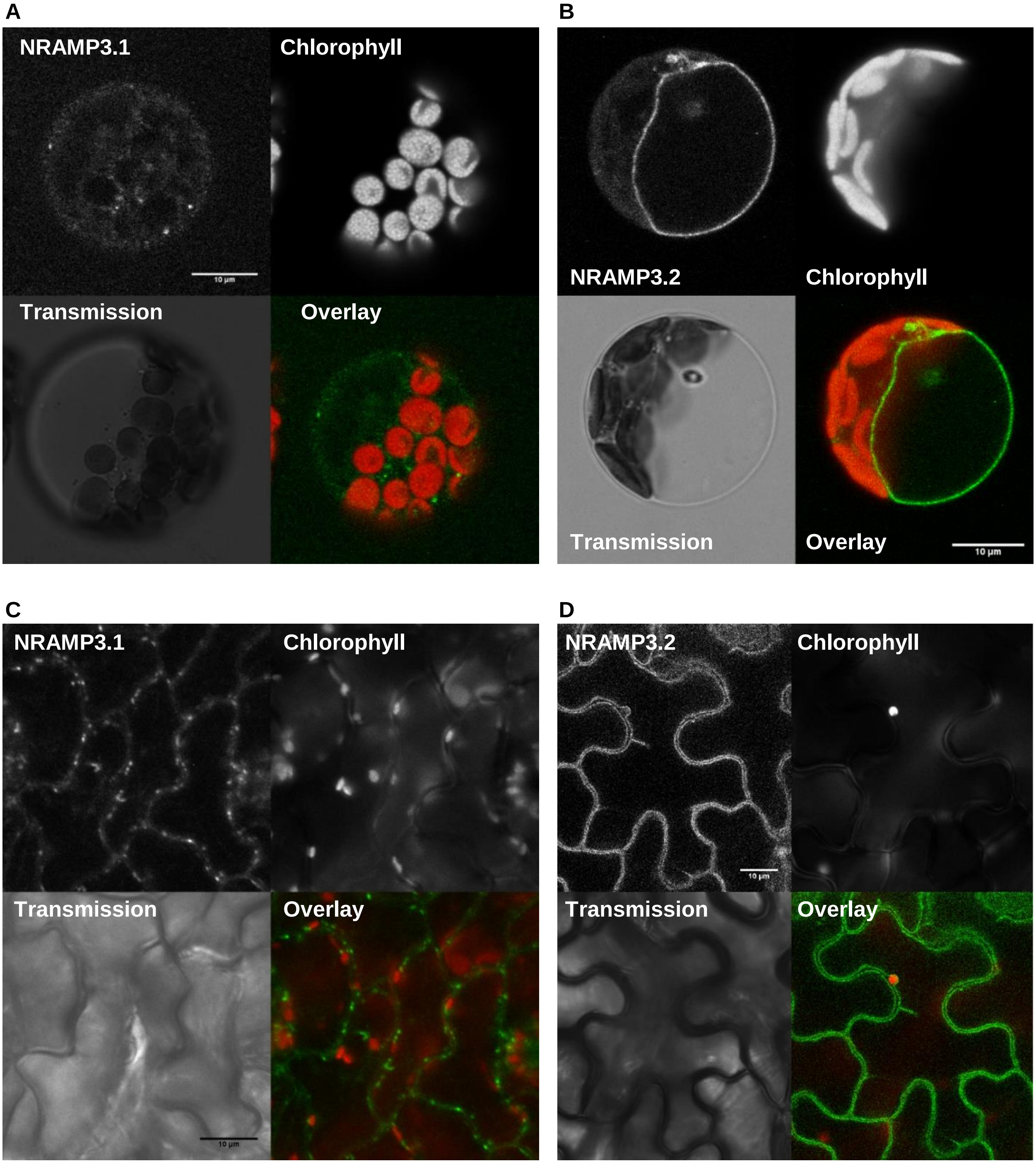


**Fig. S10.** PotriNRAMP3.1 and PotriNRAMP3.2 subcellular localizations in leaf cells. LSCM images of leaf protoplasts (A, B) or epidermis (C, D) from poplar lines expressing PotriNRAMP3.1-GFP (A, C) or PotriNRAMP3.2 B, D). The localizations of the two isoforms are similar to those observed in root cells. Scale bars : 10 µm.

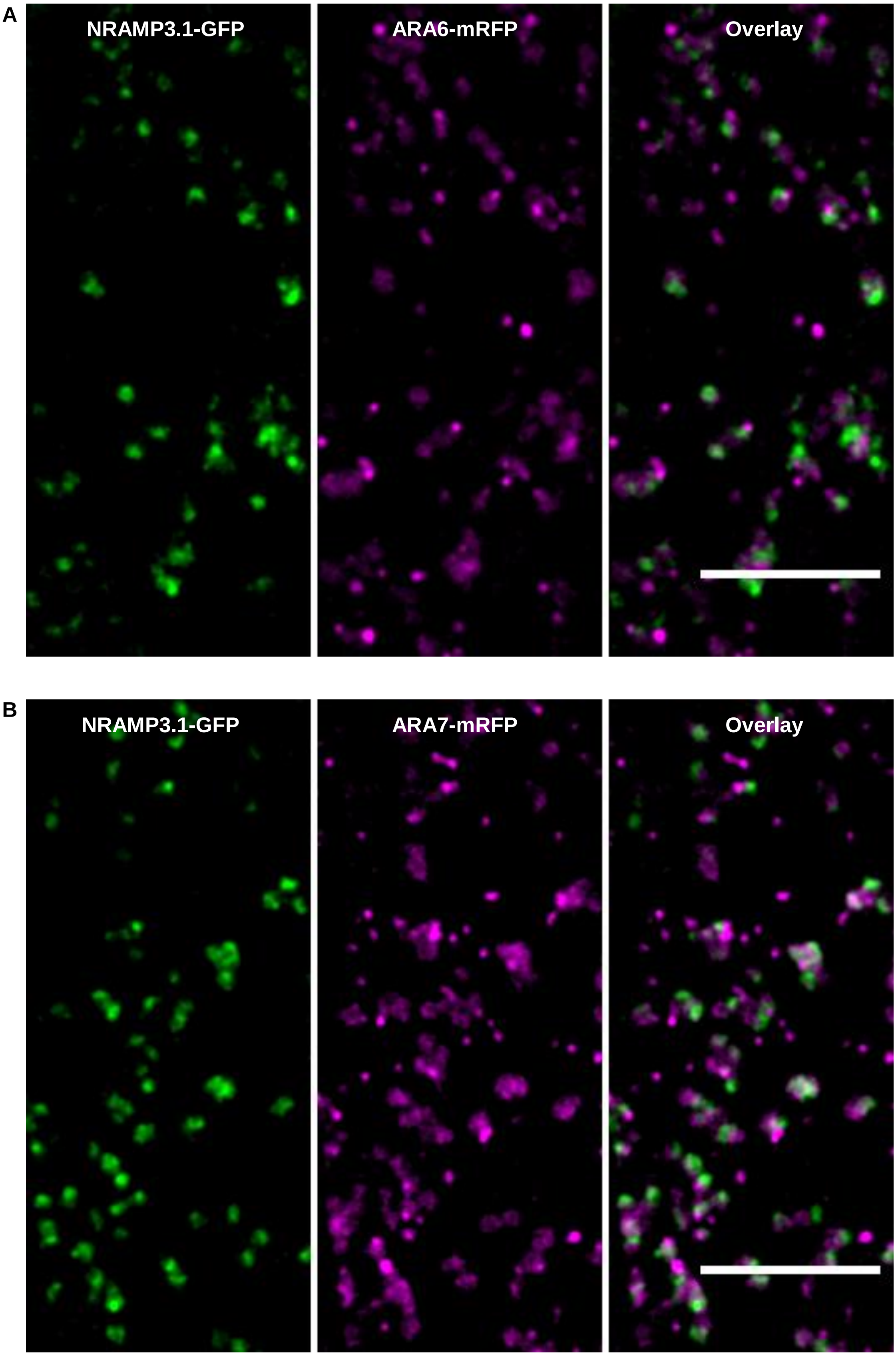


**Fig. S11.** PotriNRAMP3.1-GFP does not colocalize with late endosomal / MVB markers. PotriNRAMP3.1-GFP fluorescence is often in proximity of ARA6 (A) and ARA7 (B) but has little overlap with these markers of late endosomes / MVBs. On the merged images the overlap of GFP (green) and mRFP (magenta) channels appears white. Spinning disk confocal images were acquired in the cortical planes of root epidermal cells in the early elongation zone of Arabidopsis F1 seedlings. Scale bar : 10 µm.

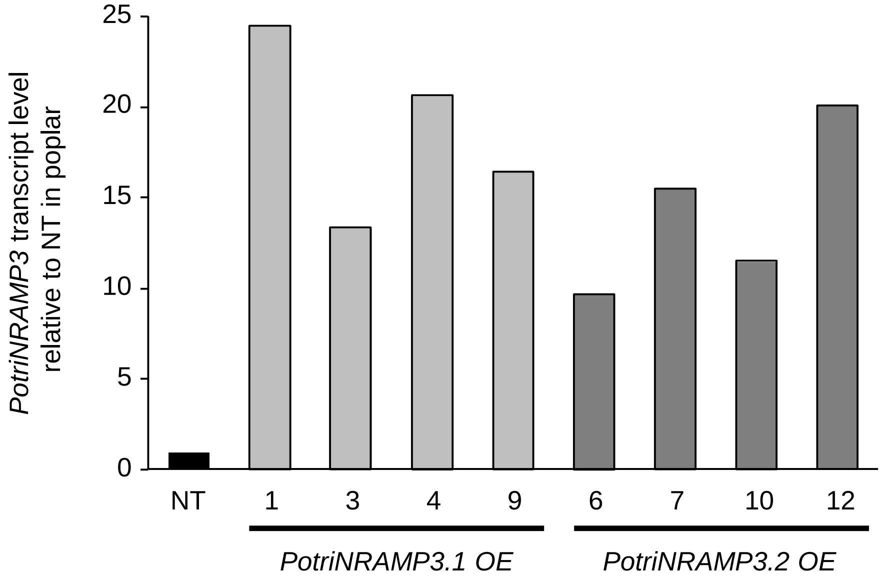


**Fig. S12.** Transcript level of *PotriNRAMP3.1* and *PotriNRAMP3.2* mRNA in leaves of transgenic poplar lines relative to NT. Relative transcript levels were determined by RT-qPCR using specific primers for each isoform and normalization by three reference genes, *i.e., PtUBQ*, *PtEF1*, and *PtPP2A*.

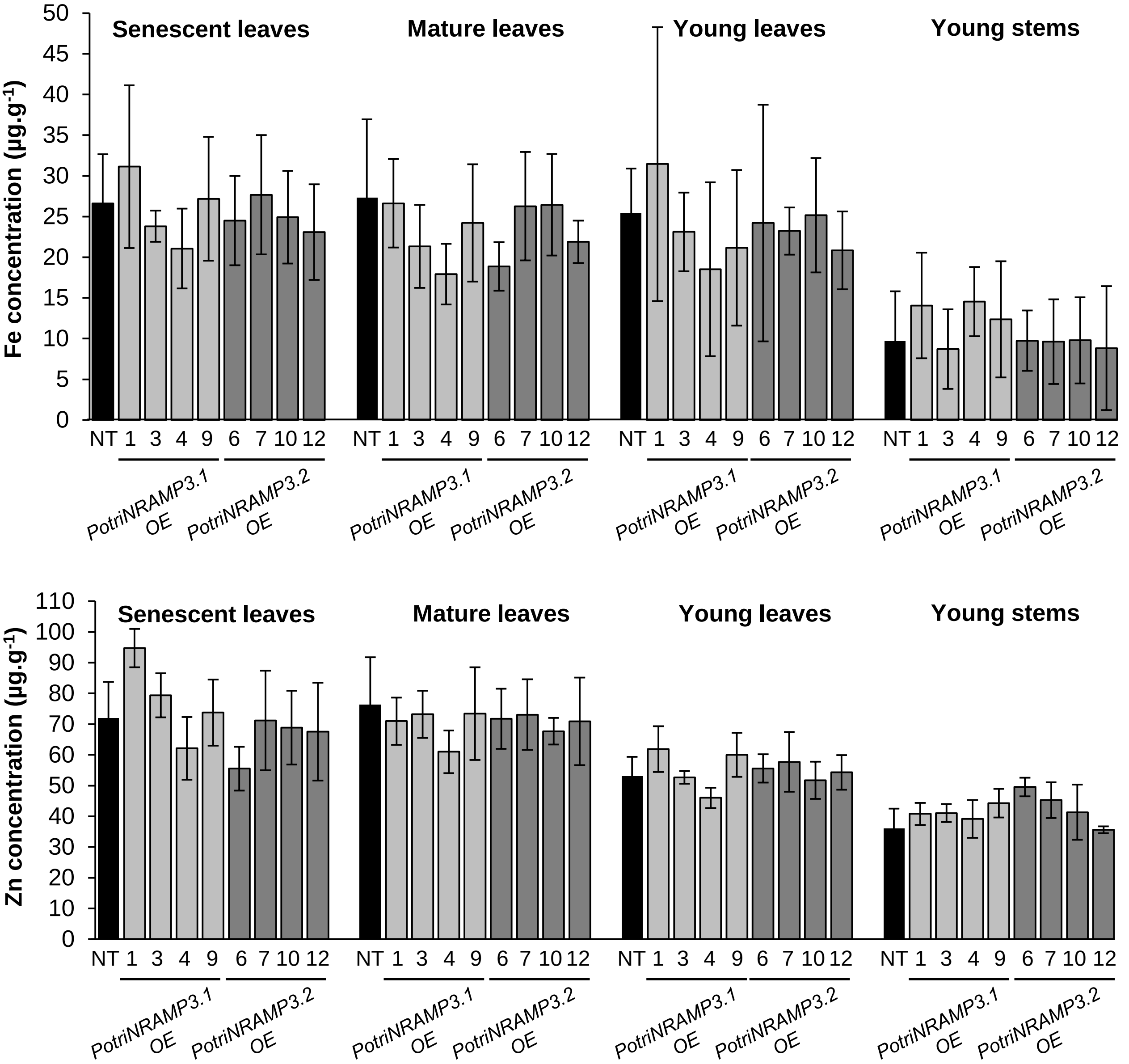


**Fig. S13.** Metal concentrations in poplar transgenic lines. Ectopic over expression of *PotriNRAMP3.1-GFP* or *PotriNRAMP3.2-GFP* does not perturb Fe or Zn distribution in poplar leaves or stems. Fe and Zn concentrations in senescent, mature, young leaves and young stems of poplars 2 months after transfer from *in vitro* to soil was determined using Atomic Emission Spectroscopy. Mean Fe or Zn concentrations of PotriNRAMP3.1-GFP or PotriNRAMP3.2-GFP OE lines were compared with NT control. Error bars represent SD (n = 4-7 plants from each transgenic line). Kruskal Wallis multiple comparison tests did not reveal any statistically significant difference among genotypes for any organ.

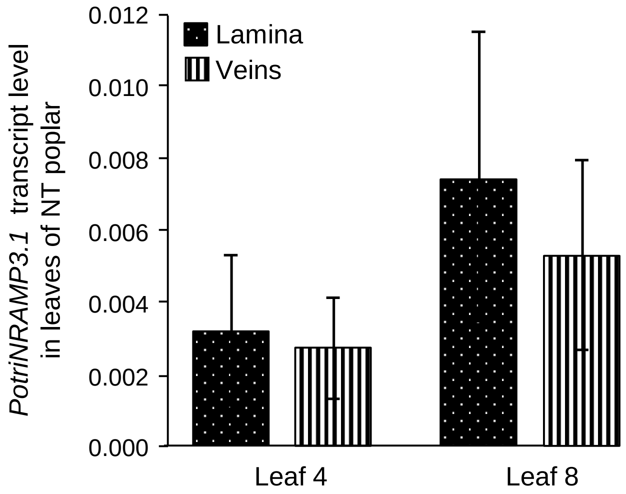


**Fig. S14.** Transcript level of *PotriNRAMP3.1* in lamina and veins of NT poplar (n = 3 leaves from independent individuals. Relative transcript levels were determined by RT-qPCR using specific primers for each isoform and normalization by three reference genes, *i.e., PtUBQ*, *PtEF1*, and *PtPP2A*. Leaves 4 and 8 correspond to the $4^{th}$ and $8^{th}$ leaf starting from the apex, respectively.

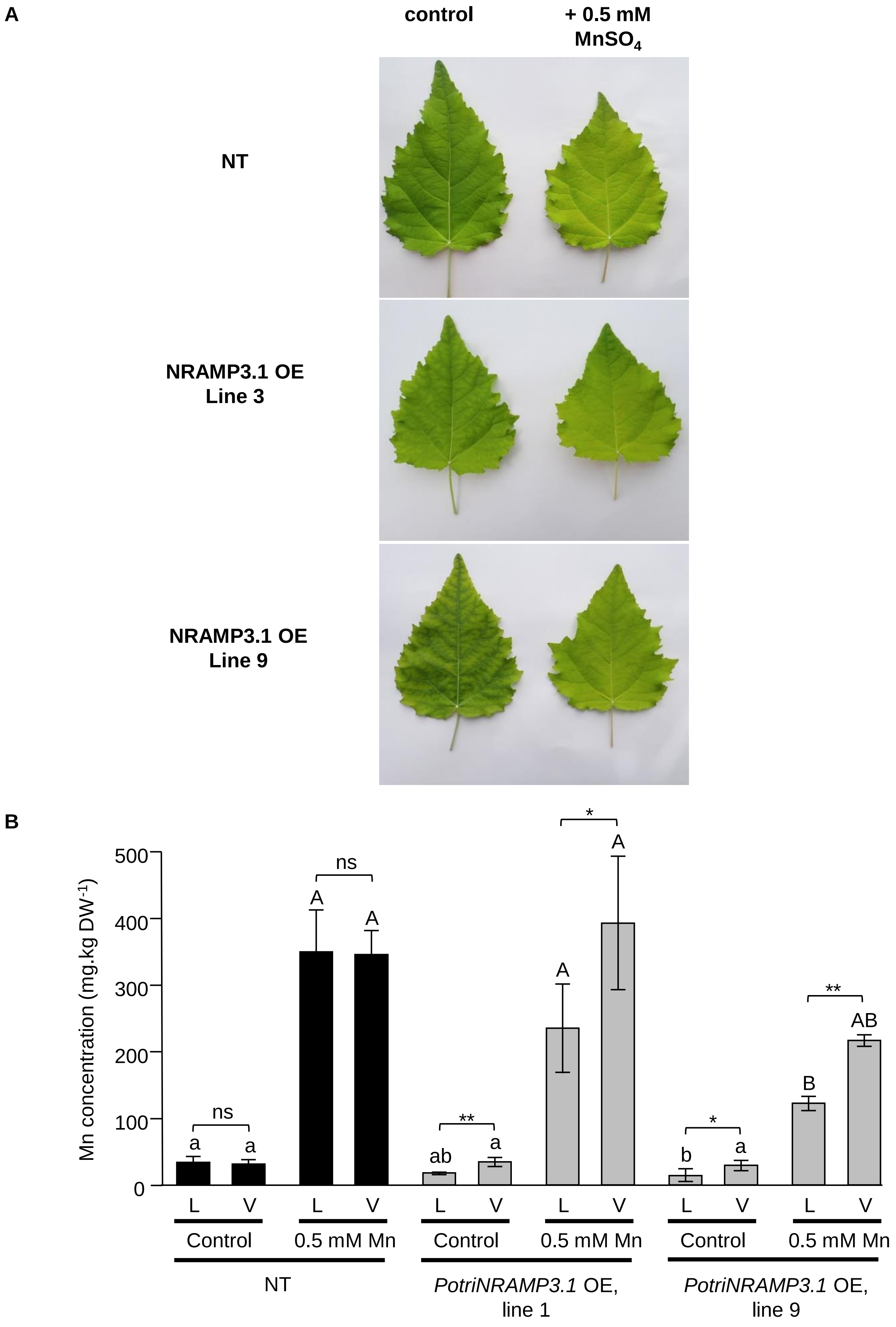


**Fig. S15.** High Mn supply mitigates the interneval chlorosis in NRAMP3.1 OE lines (A) photographs of mature leaves of the wild type and two lines overexpressing *PotriNRAMP3.1* supplied or not with 0.5 mM $MnSO_4$ through watering. (B) Mn concentration in the lamina (L) and veins (V) of the same leaves. (A) and (B) use leaves number 8-13 starting from the apex. Asterisks denote significant differences between lamina and veins according to a Mann-Whitney test (* $p < 0.05$, ** $p < 0.01$). Different letters denote significant differences among concentrations in control soil (lowercase) and soil supplemented with Mn (capitals) according to a Kruskal-Wallis test followed by Dunn's test for multiple comparison ($p < 0.05$).
