## Supplementary Materials for "Duplication of *NRAMP3* gene in poplars generated two homologous transporters with distinct functions"

*Author for correspondence

Sébastien Thomine

### Supplementary Tables

**Supplementary table S1.** List of the species and corresponding resources exploited in this work. Sequences were retrieved from the Phytozome V13 (https://phytozome-next.jgi.doe.gov/) (Goodstein et al. 2012), PopGenIE V3 (https://PopGenIE.org/) (Sjödin et al. 2009), NCBI (https://www.ncbi.nlm.nih.gov/) (Sayers et al. 2021), and 1KP Project (https://db.cngb.org/onekp/) (One Thousand Plant Transcriptomes Initiative 2019) databases.

| Species | Section (poplar) | Resource | Database | Reference |
|---|---|---|---|---|
| *Arabidopsis thaliana* | | Genome TAIR 10 | TAIR | **(Lamesch et al. 2012)** |
| *Populus trichocarpa* | Tacahamaca | Genome V4.1 | Phytozome V13 | (Tuskan et al. 2006) |
| *Populus tremula* | Populus | Genome V1.0 | PopGenIE V3 | (Lin et al. 2018) |
| *Populus tremuloides* | Populus | Genome V1.1 | PopGenIE V3 | |
| *Populus grandidentata* | Populus | Genome <V0.99 | PopGenIE V3 | UPSC Genomes Archive |
| *Populus alba* | Populus | SRA: SRR9007070 | NCBI | (Zhang et al. 2019) |
| *Populus cathayana* | Tacahamaca | SRA: SRR9007071 | NCBI | |
| *Populus simonii* | Tacahamaca | SRA: SRR9007072 | NCBI | |
| *Populus lasiocarpa* | Leucoides | SRA: SRR9007073 | NCBI | |
| *Populus maximowiczii* | Tacahamaca | SRA: SRR9007074 | NCBI | |
| *Populus euphratica* | Turanga | SRA: SRR9007075 | NCBI | |
| *Populus ussuriensis* | Tacahamaca | SRA: SRR9007077 | NCBI | |
| *Populus nigra* | Aigeiros | SRA: SRR9007078 | NCBI | |
| *Populus deltoides* | Aigeiros | SRA: SRR9007079 | NCBI | |
| *Populus mexicana* | Abaso | SRA:<br>-SRR11580850<br>-SRR11580851 | NCBI | |
| *Salix purpurea* | | Genome V5.0 | Phytozome 13 | (Zhou et al. 2018) |
| *Salix suchowensis* | | Genome | NCBI | (Dai et al. 2014) |
| *Salix brachista* | | Genome | NCBI | (Chen et al. 2019) |
| *Salix dasyclados* | | Transcriptome | 1KP Project | (One Thousand Plant Transcriptomes Initiative 2019) |
| *Salix fargesii* | | Transcriptome | 1KP Project | |
| *Salix eriocephala* | | Transcriptome | 1KP Project | |
| *Salix sachalinensis* | | Transcriptome | 1KP Project | |
| *Salix viminalis* | | Transcriptome | 1KP Project | |

**Supplementary table S2.** Sites under positive and purifying selection in *Populus NRAMP3.1* according to FEL method. Sites subjected to positive or purifying selections were estimated by the FEL method (https://www.datamonkey.org/)(Kosakovsky Pond and Frost 2005). *NRAMP3.1* coding sequences of 13 *Populus* species retrieved as indicated in Materials and Methods were used to generate a phylogenetic tree. A subset of branches encompassing the *NRAMP3.1s* were analyzed separately to estimate the synonymous substitution rate (dS) at a site (α) and the non-synonymous substitution rate (dN) at a site (β), and a maximum-likelihood approach was undertaken to calculate dN/dS rates for each codon site (ϖ). Amino-acids correspond to that of the consensus *Populus* NRAMP3.1 sequence. Dots indicate less than 90% of conservation within all investigated *Populus* NRAMP3.1 sequences. Sites subjected to positive selection are highlighted in dark ($p < 0.05$) and light green ($p < 0.1$). Sites subject to purifying selection are highlighted in dark ($p < 0.05$) and light red ($p < 0.1$). α=β : the rate estimated under the neutral model; LRT: likelihood ratio test.

| Site | amino-acid | α | β | ϖ | α=β | LRT | p-value |
|---|---|---|---|---|---|---|---|
| 1 | M | 0 | 0 | No value | 0 | 0 | 1 |
| 2 | P | 3.207 | 0 | 0 | 0.768 | 2.754 | 0.097 |
| 3 | . | 0 | 9.05 | Infinity | 3.725 | 4.756 | 0.029 |
| 4 | P | 0 | 0 | No value | 0 | 0 | 1 |
| 5 | E | 0 | 0 | No value | 0 | 0 | 1 |
| 6 | E | 0 | 0 | No value | 0 | 0 | 1 |
| 7 | D | 0 | 1.063 | Infinity | 0.736 | 0.728 | 0.394 |
| 8 | P | 5.82 | 1.186 | 0.204 | 2.494 | 1.579 | 0.209 |
| 9 | . | 0 | 2.809 | Infinity | 1.934 | 1.437 | 0.231 |
| 10 | P | 3.162 | 0 | 0 | 0.78 | 2.691 | 0.101 |
| 11 | L | 0 | 0 | No value | 0 | 0 | 1 |
| 12 | L | 2.026 | 0 | 0 | 1.044 | 1.319 | 0.251 |
| 13 | K | 0 | 2.603 | Infinity | 1.858 | 1.334 | 0.248 |
| 14 | D | 0 | 1.112 | Infinity | 0.84 | 0.563 | 0.453 |
| 15 | Q | 0 | 1.448 | Infinity | 0.991 | 0.754 | 0.385 |
| 16 | E | 0 | 0 | No value | 0 | 0 | 1 |
| 17 | E | 0 | 0 | No value | 0 | 0 | 1 |
| 18 | T | 0 | 1.3 | Infinity | 0.789 | 0.981 | 0.322 |
| 19 | A | 0 | 0 | No value | 0 | 0 | 1 |
| 20 | Y | 0 | 1.675 | Infinity | 1.262 | 0.556 | 0.456 |
| 21 | D | 0 | 0 | No value | 0 | 0 | 1 |
| 22 | S | 0 | 0 | No value | 0 | 0 | 1 |
| 23 | D | 3.393 | 1.134 | 0.334 | 1.741 | 0.509 | 0.476 |
| 24 | G | 0 | 1.378 | Infinity | 0.844 | 0.966 | 0.326 |
| 25 | K | 0 | 0 | No value | 0 | 0 | 1 |
| 26 | V | 4.189 | 0 | 0 | 1.879 | 3.269 | 0.071 |
| 27 | L | 2.005 | 0 | 0 | 0.801 | 1.819 | 0.177 |
| 28 | S | 2.724 | 4.033 | 1.481 | 3.495 | 0.095 | 0.758 |
| 29 | F | 0 | 1.965 | Infinity | 1.33 | 0.75 | 0.386 |
| 30 | G | 1.966 | 0 | 0 | 0.788 | 1.793 | 0.181 |
| 31 | I | 3.598 | 1.437 | 0.399 | 2.046 | 0.37 | 0.543 |
| 32 | D | 0 | 0 | No value | 0 | 0 | 1 |
| 33 | Y | 0 | 1.787 | Infinity | 1.199 | 0.771 | 0.38 |
| 34 | D | 0 | 1.094 | Infinity | 0.747 | 0.75 | 0.386 |
| 35 | T | 9.333 | 0 | 0 | 3.15 | 8.605 | 0.003 |
| 36 | E | 0 | 0 | No value | 0 | 0 | 1 |
| 37 | S | 2.649 | 0 | 0 | 0.699 | 2.605 | 0.107 |

| | | | | | | | |
|---|---|---|---|---|---|---|---|
| 38 | G | 0 | 0 | No value | 0 | 0 | 1 |
| 39 | G | 0 | 1.397 | Infinity | 0.689 | 1.361 | 0.243 |
| 40 | S | 0 | 0 | No value | 0 | 0 | 1 |
| 41 | T | 0 | 0 | No value | 0 | 0 | 1 |
| 42 | V | 0 | 1.693 | Infinity | 0.865 | 1.288 | 0.256 |
| 43 | V | 0 | 2.052 | Infinity | 0.981 | 1.442 | 0.23 |
| 44 | P | 1.903 | 1.077 | 0.566 | 1.374 | 0.157 | 0.692 |
| 45 | S | 2.32 | 1.47 | 0.634 | 1.826 | 0.094 | 0.759 |
| 46 | F | 0 | 0 | No value | 0 | 0 | 1 |
| 47 | S | 0 | 0 | No value | 0 | 0 | 1 |
| 48 | W | 0 | 0 | No value | 0 | 0 | 1 |
| 49 | R | 0 | 1.342 | Infinity | 0.897 | 0.767 | 0.381 |
| 50 | K | 4.546 | 0 | 0 | 0.898 | 3.182 | 0.074 |
| 51 | L | 0 | 0 | No value | 0 | 0 | 1 |
| 52 | W | 0 | 0 | No value | 0 | 0 | 1 |
| 53 | L | 2.183 | 0 | 0 | 1.044 | 1.408 | 0.235 |
| 54 | F | 5.723 | 0 | 0 | 1.224 | 3.249 | 0.071 |
| 55 | T | 0 | 0 | No value | 0 | 0 | 1 |
| 56 | G | 0 | 0 | No value | 0 | 0 | 1 |
| 57 | P | 0 | 0 | No value | 0 | 0 | 1 |
| 58 | G | 1.949 | 0 | 0 | 0.784 | 1.781 | 0.182 |
| 59 | F | 0 | 0 | No value | 0 | 0 | 1 |
| 60 | L | 0 | 0 | No value | 0 | 0 | 1 |
| 61 | M | 0 | 0 | No value | 0 | 0 | 1 |
| 62 | C | 1.704 | 1.575 | 0.924 | 1.636 | 0.003 | 0.96 |
| 63 | I | 0 | 0 | No value | 0 | 0 | 1 |
| 64 | A | 3.044 | 0 | 0 | 0.861 | 2.566 | 0.109 |
| 65 | F | 0 | 0 | No value | 0 | 0 | 1 |
| 66 | L | 0 | 0 | No value | 0 | 0 | 1 |
| 67 | D | 3.407 | 0 | 0 | 0.837 | 2.77 | 0.096 |
| 68 | P | 0 | 0 | No value | 0 | 0 | 1 |
| 69 | G | 6.296 | 0 | 0 | 2.577 | 5.407 | 0.02 |
| 70 | N | 0 | 0 | No value | 0 | 0 | 1 |
| 71 | L | 0 | 0 | No value | 0 | 0 | 1 |
| 72 | E | 0 | 0 | No value | 0 | 0 | 1 |
| 73 | G | 0 | 0 | No value | 0 | 0 | 1 |
| 74 | D | 0 | 0 | No value | 0 | 0 | 1 |
| 75 | L | 0 | 0 | No value | 0 | 0 | 1 |
| 76 | Q | 0 | 0 | No value | 0 | 0 | 1 |
| 77 | A | 0 | 0 | No value | 0 | 0 | 1 |
| 78 | G | 0 | 0 | No value | 0 | 0 | 1 |
| 79 | A | 1.979 | 0 | 0 | 0.727 | 1.943 | 0.163 |
| 80 | I | 2.45 | 0 | 0 | 0.853 | 2.105 | 0.147 |
| 81 | A | 1.942 | 0 | 0 | 0.725 | 1.931 | 0.165 |
| 82 | G | 6.48 | 0 | 0 | 2.625 | 5.495 | 0.019 |
| 83 | Y | 3.553 | 0 | 0 | 1.106 | 2.295 | 0.13 |
| 84 | S | 0 | 0 | No value | 0 | 0 | 1 |
| 85 | L | 0 | 0 | No value | 0 | 0 | 1 |
| 86 | L | 0 | 0 | No value | 0 | 0 | 1 |

| | | | | | | | |
|---|---|---|---|---|---|---|---|
| 87 | W | 0 | 0 | No value | 0 | 0 | 1 |
| 88 | L | 0 | 0 | No value | 0 | 0 | 1 |
| 89 | L | 1.929 | 0 | 0 | 0.789 | 1.774 | 0.183 |
| 90 | L | 0 | 0 | No value | 0 | 0 | 1 |
| 91 | W | 0 | 0 | No value | 0 | 0 | 1 |
| 92 | A | 0 | 0 | No value | 0 | 0 | 1 |
| 93 | T | 0 | 0 | No value | 0 | 0 | 1 |
| 94 | A | 0 | 0 | No value | 0 | 0 | 1 |
| 95 | M | 0 | 0 | No value | 0 | 0 | 1 |
| 96 | G | 6.514 | 0 | 0 | 1.995 | 4.826 | 0.028 |
| 97 | L | 0 | 0 | No value | 0 | 0 | 1 |
| 98 | L | 0 | 0 | No value | 0 | 0 | 1 |
| 99 | V | 0 | 0 | No value | 0 | 0 | 1 |
| 100 | Q | 0 | 0 | No value | 0 | 0 | 1 |
| 101 | L | 0 | 0 | No value | 0 | 0 | 1 |
| 102 | L | 1.996 | 0 | 0 | 0.933 | 1.52 | 0.218 |
| 103 | S | 0 | 0 | No value | 0 | 0 | 1 |
| 104 | A | 3.973 | 0 | 0 | 1.503 | 3.862 | 0.049 |
| 105 | R | 0 | 0 | No value | 0 | 0 | 1 |
| 106 | L | 0 | 0 | No value | 0 | 0 | 1 |
| 107 | G | 1.927 | 0 | 0 | 0.817 | 1.663 | 0.197 |
| 108 | V | 0 | 0 | No value | 0 | 0 | 1 |
| 109 | A | 0 | 0 | No value | 0 | 0 | 1 |
| 110 | T | 0 | 0 | No value | 0 | 0 | 1 |
| 111 | G | 3.292 | 0 | 0 | 1.156 | 2.579 | 0.108 |
| 112 | R | 2.398 | 0 | 0 | 0.75 | 2.241 | 0.134 |
| 113 | H | 0 | 0 | No value | 0 | 0 | 1 |
| 114 | L | 4.346 | 0 | 0 | 2.169 | 2.687 | 0.101 |
| 115 | A | 0 | 0 | No value | 0 | 0 | 1 |
| 116 | E | 0 | 0 | No value | 0 | 0 | 1 |
| 117 | L | 1.165 | 0 | 0 | 0.726 | 0.948 | 0.33 |
| 118 | C | 0 | 1.503 | Infinity | 1.192 | 0.462 | 0.496 |
| 119 | R | 0 | 0 | No value | 0 | 0 | 1 |
| 120 | E | 0 | 0 | No value | 0 | 0 | 1 |
| 121 | E | 0 | 0 | No value | 0 | 0 | 1 |
| 122 | Y | 0 | 0 | No value | 0 | 0 | 1 |
| 123 | P | 0 | 0 | No value | 0 | 0 | 1 |
| 124 | T | 4.515 | 0 | 0 | 1.426 | 4.283 | 0.038 |
| 125 | W | 0 | 0 | No value | 0 | 0 | 1 |
| 126 | A | 3.245 | 0 | 0 | 0.85 | 2.575 | 0.109 |
| 127 | R | 0 | 4.487 | Infinity | 1.927 | 3.294 | 0.07 |
| 128 | M | 0 | 0 | No value | 0 | 0 | 1 |
| 129 | I | 0 | 1.419 | Infinity | 0.968 | 0.732 | 0.392 |
| 130 | L | 0 | 0 | No value | 0 | 0 | 1 |
| 131 | W | 0 | 0 | No value | 0 | 0 | 1 |
| 132 | I | 0 | 0 | No value | 0 | 0 | 1 |
| 133 | M | 0 | 0 | No value | 0 | 0 | 1 |
| 134 | A | 0 | 0 | No value | 0 | 0 | 1 |
| 135 | E | 0 | 0 | No value | 0 | 0 | 1 |

| | | | | | | | |
|---|---|---|---|---|---|---|---|
| 136 | L | 0 | 0 | No value | 0 | 0 | 1 |
| 137 | A | 0 | 0 | No value | 0 | 0 | 1 |
| 138 | L | 0 | 0 | No value | 0 | 0 | 1 |
| 139 | I | 0 | 0 | No value | 0 | 0 | 1 |
| 140 | G | 0 | 0 | No value | 0 | 0 | 1 |
| 141 | A | 0 | 0 | No value | 0 | 0 | 1 |
| 142 | D | 0 | 0 | No value | 0 | 0 | 1 |
| 143 | I | 0 | 0 | No value | 0 | 0 | 1 |
| 144 | Q | 0 | 0 | No value | 0 | 0 | 1 |
| 145 | E | 3.161 | 0 | 0 | 0.885 | 2.509 | 0.113 |
| 146 | V | 0 | 0 | No value | 0 | 0 | 1 |
| 147 | I | 0 | 0 | No value | 0 | 0 | 1 |
| 148 | G | 1.908 | 0 | 0 | 0.792 | 1.741 | 0.187 |
| 149 | S | 0 | 0 | No value | 0 | 0 | 1 |
| 150 | A | 0 | 0 | No value | 0 | 0 | 1 |
| 151 | I | 14.276 | 0 | 0 | 2.316 | 7.011 | 0.008 |
| 152 | A | 3.118 | 0 | 0 | 0.844 | 2.533 | 0.111 |
| 153 | I | 0 | 0 | No value | 0 | 0 | 1 |
| 154 | Q | 0 | 1.412 | Infinity | 0.943 | 0.793 | 0.373 |
| 155 | I | 2.45 | 0 | 0 | 0.853 | 2.105 | 0.147 |
| 156 | L | 0 | 0 | No value | 0 | 0 | 1 |
| 157 | S | 0 | 0 | No value | 0 | 0 | 1 |
| 158 | N | 0 | 0 | No value | 0 | 0 | 1 |
| 159 | G | 0 | 0 | No value | 0 | 0 | 1 |
| 160 | V | 0 | 2.022 | Infinity | 1.288 | 0.882 | 0.348 |
| 161 | L | 0 | 0 | No value | 0 | 0 | 1 |
| 162 | P | 0 | 0 | No value | 0 | 0 | 1 |
| 163 | L | 2.038 | 0 | 0 | 1.01 | 1.345 | 0.246 |
| 164 | W | 0 | 0 | No value | 0 | 0 | 1 |
| 165 | A | 0 | 0 | No value | 0 | 0 | 1 |
| 166 | G | 0 | 0 | No value | 0 | 0 | 1 |
| 167 | V | 0 | 0 | No value | 0 | 0 | 1 |
| 168 | I | 0 | 1.297 | Infinity | 0.916 | 0.684 | 0.408 |
| 169 | I | 0 | 0 | No value | 0 | 0 | 1 |
| 170 | T | 0 | 0 | No value | 0 | 0 | 1 |
| 171 | A | 3.028 | 0 | 0 | 0.836 | 2.487 | 0.115 |
| 172 | S | 4.936 | 3.611 | 0.732 | 4.175 | 0.092 | 0.762 |
| 173 | D | 0 | 0 | No value | 0 | 0 | 1 |
| 174 | C | 0 | 1.453 | Infinity | 0.911 | 0.898 | 0.343 |
| 175 | F | 3.553 | 0 | 0 | 1.079 | 2.353 | 0.125 |
| 176 | I | 1.775 | 0 | 0 | 0.729 | 1.749 | 0.186 |
| 177 | F | 0 | 0 | No value | 0 | 0 | 1 |
| 178 | L | 0 | 0 | No value | 0 | 0 | 1 |
| 179 | F | 0 | 0 | No value | 0 | 0 | 1 |
| 180 | L | 2.072 | 0 | 0 | 0.857 | 1.779 | 0.182 |
| 181 | E | 0 | 0 | No value | 0 | 0 | 1 |
| 182 | N | 0 | 0 | No value | 0 | 0 | 1 |
| 183 | Y | 0 | 0 | No value | 0 | 0 | 1 |

| | | | | | | | |
|---|---|---|---|---|---|---|---|
| 184 | G | 3.106 | 0 | 0 | 0.923 | 2.349 | 0.125 |
| 185 | V | 0 | 0 | No value | 0 | 0 | 1 |
| 186 | R | 2.349 | 0 | 0 | 0.767 | 2.205 | 0.138 |
| 187 | K | 0 | 0 | No value | 0 | 0 | 1 |
| 188 | L | 2.025 | 0 | 0 | 1.01 | 1.345 | 0.246 |
| 189 | E | 0 | 0 | No value | 0 | 0 | 1 |
| 190 | A | 0 | 0 | No value | 0 | 0 | 1 |
| 191 | A | 2.973 | 1.332 | 0.448 | 1.863 | 0.309 | 0.578 |
| 192 | F | 0 | 0 | No value | 0 | 0 | 1 |
| 193 | G | 0 | 1.616 | Infinity | 0.85 | 1.245 | 0.265 |
| 194 | I | 9.042 | 1.417 | 0.157 | 3.898 | 2.654 | 0.103 |
| 195 | L | 1.929 | 0 | 0 | 0.789 | 1.774 | 0.183 |
| 196 | I | 0 | 0 | No value | 0 | 0 | 1 |
| 197 | G | 0 | 0 | No value | 0 | 0 | 1 |
| 198 | I | 0 | 0 | No value | 0 | 0 | 1 |
| 199 | M | 0 | 0 | No value | 0 | 0 | 1 |
| 200 | A | 0 | 0 | No value | 0 | 0 | 1 |
| 201 | V | 2.955 | 0 | 0 | 1.058 | 2.107 | 0.147 |
| 202 | T | 0 | 1.118 | Infinity | 0.679 | 0.957 | 0.328 |
| 203 | F | 0 | 0 | No value | 0 | 0 | 1 |
| 204 | A | 1.912 | 1.515 | 0.792 | 1.689 | 0.026 | 0.872 |
| 205 | W | 0 | 1.829 | Infinity | 1.806 | 0.025 | 0.874 |
| 206 | M | 0 | 0 | No value | 0 | 0 | 1 |
| 207 | F | 0 | 0 | No value | 0 | 0 | 1 |
| 208 | A | 2.996 | 0 | 0 | 0.868 | 2.516 | 0.113 |
| 209 | D | 0 | 0 | No value | 0 | 0 | 1 |
| 210 | A | 0 | 0 | No value | 0 | 0 | 1 |
| 211 | K | 0 | 0 | No value | 0 | 0 | 1 |
| 212 | P | 0 | 0 | No value | 0 | 0 | 1 |
| 213 | S | 0 | 0 | No value | 0 | 0 | 1 |
| 214 | A | 1.426 | 0 | 0 | 0.642 | 1.594 | 0.207 |
| 215 | P | 3.9 | 1.138 | 0.292 | 2.153 | 0.986 | 0.321 |
| 216 | E | 0 | 0 | No value | 0 | 0 | 1 |
| 217 | L | 3.133 | 0 | 0 | 0.924 | 2.35 | 0.125 |
| 218 | F | 0 | 0 | No value | 0 | 0 | 1 |
| 219 | L | 7.839 | 0 | 0 | 4.251 | 5.245 | 0.022 |
| 220 | G | 1.943 | 0 | 0 | 0.788 | 1.783 | 0.182 |
| 221 | I | 1.744 | 0 | 0 | 0.727 | 1.738 | 0.187 |
| 222 | L | 0 | 0 | No value | 0 | 0 | 1 |
| 223 | I | 0 | 0 | No value | 0 | 0 | 1 |
| 224 | P | 1.881 | 0 | 0 | 0.657 | 2.046 | 0.153 |
| 225 | K | 0 | 0 | No value | 0 | 0 | 1 |
| 226 | L | 0 | 0 | No value | 0 | 0 | 1 |
| 227 | S | 0 | 0 | No value | 0 | 0 | 1 |
| 228 | S | 0 | 0 | No value | 0 | 0 | 1 |
| 229 | K | 0 | 1.168 | Infinity | 0.809 | 0.712 | 0.399 |
| 230 | T | 0 | 0 | No value | 0 | 0 | 1 |
| 231 | I | 0 | 0 | No value | 0 | 0 | 1 |
| 232 | K | 0 | 1.242 | Infinity | 0.888 | 0.659 | 0.417 |

| | | | | | | | |
|---|---|---|---|---|---|---|---|
| 233 | Q | 3.606 | 0 | 0 | 0.947 | 2.53 | 0.112 |
| 234 | A | 3.324 | 0 | 0 | 0.858 | 2.62 | 0.106 |
| 235 | V | 9.822 | 0 | 0 | 2.802 | 5.58 | 0.018 |
| 236 | G | 0 | 0 | No value | 0 | 0 | 1 |
| 237 | V | 0 | 0 | No value | 0 | 0 | 1 |
| 238 | V | 0 | 0 | No value | 0 | 0 | 1 |
| 239 | G | 0 | 0 | No value | 0 | 0 | 1 |
| 240 | C | 0 | 0 | No value | 0 | 0 | 1 |
| 241 | I | 0 | 0 | No value | 0 | 0 | 1 |
| 242 | I | 0 | 0 | No value | 0 | 0 | 1 |
| 243 | M | 0 | 0 | No value | 0 | 0 | 1 |
| 244 | P | 0 | 0 | No value | 0 | 0 | 1 |
| 245 | H | 0 | 0 | No value | 0 | 0 | 1 |
| 246 | N | 0 | 0 | No value | 0 | 0 | 1 |
| 247 | V | 0 | 0 | No value | 0 | 0 | 1 |
| 248 | F | 0 | 0 | No value | 0 | 0 | 1 |
| 249 | L | 0 | 0 | No value | 0 | 0 | 1 |
| 250 | H | 0 | 0 | No value | 0 | 0 | 1 |
| 251 | S | 0 | 0 | No value | 0 | 0 | 1 |
| 252 | A | 0 | 0 | No value | 0 | 0 | 1 |
| 253 | L | 0 | 0 | No value | 0 | 0 | 1 |
| 254 | V | 1.917 | 0 | 0 | 0.861 | 1.569 | 0.21 |
| 255 | Q | 0 | 0 | No value | 0 | 0 | 1 |
| 256 | S | 0 | 0 | No value | 0 | 0 | 1 |
| 257 | R | 0 | 0 | No value | 0 | 0 | 1 |
| 258 | E | 0 | 0 | No value | 0 | 0 | 1 |
| 259 | I | 12.387 | 0 | 0 | 4.028 | 8.678 | 0.003 |
| 260 | D | 0 | 0 | No value | 0 | 0 | 1 |
| 261 | H | 0 | 2.029 | Infinity | 1.417 | 1.38 | 0.24 |
| 262 | N | 0 | 0 | No value | 0 | 0 | 1 |
| 263 | K | 0 | 0 | No value | 0 | 0 | 1 |
| 264 | K | 0 | 0 | No value | 0 | 0 | 1 |
| 265 | G | 0 | 0 | No value | 0 | 0 | 1 |
| 266 | Q | 1.963 | 2.898 | 1.476 | 2.507 | 0.097 | 0.755 |
| 267 | V | 0 | 0 | No value | 0 | 0 | 1 |
| 268 | Q | 0 | 0 | No value | 0 | 0 | 1 |
| 269 | E | 0 | 0 | No value | 0 | 0 | 1 |
| 270 | A | 0 | 0 | No value | 0 | 0 | 1 |
| 271 | L | 1.451 | 0 | 0 | 0.692 | 1.465 | 0.226 |
| 272 | R | 0 | 0 | No value | 0 | 0 | 1 |
| 273 | Y | 0 | 0 | No value | 0 | 0 | 1 |
| 274 | Y | 0 | 0 | No value | 0 | 0 | 1 |
| 275 | S | 0 | 0 | No value | 0 | 0 | 1 |
| 276 | I | 0 | 0 | No value | 0 | 0 | 1 |
| 277 | E | 0 | 0 | No value | 0 | 0 | 1 |
| 278 | S | 0 | 0 | No value | 0 | 0 | 1 |
| 279 | T | 2.004 | 0 | 0 | 0.698 | 2.104 | 0.147 |
| 280 | A | 0 | 1.182 | Infinity | 0.83 | 0.701 | 0.402 |

| | | | | | | | |
|---|---|---|---|---|---|---|---|
| 281 | A | 0 | 0 | No value | 0 | 0 | 1 |
| 282 | L | 0 | 0 | No value | 0 | 0 | 1 |
| 283 | A | 0 | 1.312 | Infinity | 0.753 | 1.092 | 0.296 |
| 284 | I | 0 | 0 | No value | 0 | 0 | 1 |
| 285 | S | 6.339 | 0 | 0 | 2.916 | 4.512 | 0.034 |
| 286 | F | 0 | 0 | No value | 0 | 0 | 1 |
| 287 | M | 2.025 | 1.506 | 0.744 | 1.605 | 0.015 | 0.904 |
| 288 | I | 0 | 1.345 | Infinity | 0.737 | 1.156 | 0.282 |
| 289 | N | 0 | 0 | No value | 0 | 0 | 1 |
| 290 | L | 0 | 0 | No value | 0 | 0 | 1 |
| 291 | F | 0 | 0 | No value | 0 | 0 | 1 |
| 292 | V | 0 | 0 | No value | 0 | 0 | 1 |
| 293 | T | 4.524 | 0 | 0 | 1.449 | 4.343 | 0.037 |
| 294 | T | 6.503 | 0 | 0 | 2.223 | 6.293 | 0.012 |
| 295 | . | 3.263 | 8.293 | 2.541 | 6.069 | 0.508 | 0.476 |
| 296 | F | 0 | 0 | No value | 0 | 0 | 1 |
| 297 | A | 0 | 0 | No value | 0 | 0 | 1 |
| 298 | K | 0 | 0 | No value | 0 | 0 | 1 |
| 299 | G | 0 | 0 | No value | 0 | 0 | 1 |
| 300 | F | 6.037 | 0 | 0 | 1.863 | 4.385 | 0.036 |
| 301 | H | 8.263 | 1.107 | 0.134 | 2.543 | 2.435 | 0.119 |
| 302 | G | 0 | 1.377 | Infinity | 0.781 | 1.105 | 0.293 |
| 303 | T | 0 | 0 | No value | 0 | 0 | 1 |
| 304 | E | 0 | 0 | No value | 0 | 0 | 1 |
| 305 | L | 1.178 | 0 | 0 | 0.726 | 0.948 | 0.33 |
| 306 | A | 0 | 0 | No value | 0 | 0 | 1 |
| 307 | N | 0 | 0 | No value | 0 | 0 | 1 |
| 308 | S | 0 | 0 | No value | 0 | 0 | 1 |
| 309 | I | 0 | 0 | No value | 0 | 0 | 1 |
| 310 | G | 0 | 0 | No value | 0 | 0 | 1 |
| 311 | L | 0 | 0 | No value | 0 | 0 | 1 |
| 312 | V | 0 | 0 | No value | 0 | 0 | 1 |
| 313 | N | 0 | 0 | No value | 0 | 0 | 1 |
| 314 | A | 0 | 0 | No value | 0 | 0 | 1 |
| 315 | G | 0 | 0 | No value | 0 | 0 | 1 |
| 316 | Q | 0 | 0 | No value | 0 | 0 | 1 |
| 317 | Y | 0 | 0 | No value | 0 | 0 | 1 |
| 318 | L | 0 | 0 | No value | 0 | 0 | 1 |
| 319 | Q | 3.691 | 0 | 0 | 0.928 | 2.597 | 0.107 |
| 320 | D | 7.486 | 0 | 0 | 1.843 | 5.493 | 0.019 |
| 321 | K | 0 | 0 | No value | 0 | 0 | 1 |
| 322 | Y | 0 | 0 | No value | 0 | 0 | 1 |
| 323 | G | 1.951 | 0 | 0 | 0.784 | 1.776 | 0.183 |
| 324 | G | 0 | 0 | No value | 0 | 0 | 1 |
| 325 | G | 1.91 | 0 | 0 | 0.816 | 1.664 | 0.197 |
| 326 | F | 0 | 0 | No value | 0 | 0 | 1 |
| 327 | F | 0 | 0 | No value | 0 | 0 | 1 |
| 328 | P | 0 | 0 | No value | 0 | 0 | 1 |
| 329 | I | 0 | 0 | No value | 0 | 0 | 1 |

| | | | | | | | |
|---|---|---|---|---|---|---|---|
| 330 | L | 0 | 2.266 | Infinity | 1.071 | 1.454 | 0.228 |
| 331 | Y | 0 | 0 | No value | 0 | 0 | 1 |
| 332 | I | 0 | 0 | No value | 0 | 0 | 1 |
| 333 | W | 0 | 0 | No value | 0 | 0 | 1 |
| 334 | G | 3.127 | 0 | 0 | 0.923 | 2.349 | 0.125 |
| 335 | I | 4.186 | 0 | 0 | 0.983 | 2.763 | 0.096 |
| 336 | G | 1.908 | 0 | 0 | 0.792 | 1.741 | 0.187 |
| 337 | L | 0 | 0 | No value | 0 | 0 | 1 |
| 338 | L | 0 | 0 | No value | 0 | 0 | 1 |
| 339 | A | 0 | 0 | No value | 0 | 0 | 1 |
| 340 | A | 0 | 0 | No value | 0 | 0 | 1 |
| 341 | G | 0 | 0 | No value | 0 | 0 | 1 |
| 342 | Q | 0 | 0 | No value | 0 | 0 | 1 |
| 343 | S | 7.48 | 0 | 0 | 1.669 | 5.858 | 0.016 |
| 344 | S | 0 | 0 | No value | 0 | 0 | 1 |
| 345 | T | 0 | 0 | No value | 0 | 0 | 1 |
| 346 | I | 0 | 0 | No value | 0 | 0 | 1 |
| 347 | T | 0 | 0 | No value | 0 | 0 | 1 |
| 348 | G | 1.454 | 0 | 0 | 0.693 | 1.475 | 0.224 |
| 349 | T | 3.129 | 0 | 0 | 0.787 | 2.663 | 0.103 |
| 350 | Y | 0 | 0 | No value | 0 | 0 | 1 |
| 351 | A | 0 | 0 | No value | 0 | 0 | 1 |
| 352 | G | 3.997 | 0 | 0 | 1.643 | 3.514 | 0.061 |
| 353 | Q | 0 | 0 | No value | 0 | 0 | 1 |
| 354 | F | 0 | 0 | No value | 0 | 0 | 1 |
| 355 | I | 0 | 0 | No value | 0 | 0 | 1 |
| 356 | M | 0 | 0 | No value | 0 | 0 | 1 |
| 357 | G | 0 | 0 | No value | 0 | 0 | 1 |
| 358 | G | 0 | 0 | No value | 0 | 0 | 1 |
| 359 | F | 0 | 0 | No value | 0 | 0 | 1 |
| 360 | L | 0 | 0 | No value | 0 | 0 | 1 |
| 361 | N | 0 | 0 | No value | 0 | 0 | 1 |
| 362 | L | 0 | 0 | No value | 0 | 0 | 1 |
| 363 | G | 5.852 | 1.344 | 0.23 | 2.839 | 1.469 | 0.225 |
| 364 | L | 0 | 0 | No value | 0 | 0 | 1 |
| 365 | K | 0 | 0 | No value | 0 | 0 | 1 |
| 366 | K | 0 | 0 | No value | 0 | 0 | 1 |
| 367 | W | 0 | 0 | No value | 0 | 0 | 1 |
| 368 | L | 4.046 | 0 | 0 | 2.387 | 3.039 | 0.081 |
| 369 | R | 0 | 0 | No value | 0 | 0 | 1 |
| 370 | A | 0 | 0 | No value | 0 | 0 | 1 |
| 371 | L | 2.048 | 0 | 0 | 1.011 | 1.347 | 0.246 |
| 372 | I | 2.45 | 0 | 0 | 0.853 | 2.105 | 0.147 |
| 373 | T | 0 | 0 | No value | 0 | 0 | 1 |
| 374 | R | 0 | 0 | No value | 0 | 0 | 1 |
| 375 | S | 0 | 0 | No value | 0 | 0 | 1 |
| 376 | C | 0 | 0 | No value | 0 | 0 | 1 |
| 377 | A | 0 | 0 | No value | 0 | 0 | 1 |
| 378 | I | 1.758 | 0 | 0 | 0.729 | 1.747 | 0.186 |

| | | | | | | | |
|---|---|---|---|---|---|---|---|
| 379 | I | 0 | 1.345 | Infinity | 0.728 | 1.166 | 0.28 |
| 380 | P | 0 | 0 | No value | 0 | 0 | 1 |
| 381 | T | 0 | 0 | No value | 0 | 0 | 1 |
| 382 | I | 0 | 2.192 | Infinity | 1.978 | 0.421 | 0.517 |
| 383 | I | 0 | 0 | No value | 0 | 0 | 1 |
| 384 | V | 0 | 0 | No value | 0 | 0 | 1 |
| 385 | A | 1.925 | 0 | 0 | 0.719 | 1.912 | 0.167 |
| 386 | L | 0 | 0 | No value | 0 | 0 | 1 |
| 387 | V | 3.186 | 0 | 0 | 1.053 | 2.147 | 0.143 |
| 388 | F | 0 | 0 | No value | 0 | 0 | 1 |
| 389 | D | 0 | 0 | No value | 0 | 0 | 1 |
| 390 | T | 1.994 | 0 | 0 | 0.691 | 2.061 | 0.151 |
| 391 | S | 7.301 | 0 | 0 | 2.68 | 5.831 | 0.016 |
| 392 | E | 0 | 0 | No value | 0 | 0 | 1 |
| 393 | D | 0 | 0 | No value | 0 | 0 | 1 |
| 394 | S | 1.925 | 0 | 0 | 0.909 | 1.447 | 0.229 |
| 395 | L | 0 | 0 | No value | 0 | 0 | 1 |
| 396 | D | 0 | 0 | No value | 0 | 0 | 1 |
| 397 | V | 0 | 0 | No value | 0 | 0 | 1 |
| 398 | L | 0 | 0 | No value | 0 | 0 | 1 |
| 399 | N | 0 | 0 | No value | 0 | 0 | 1 |
| 400 | E | 0 | 0 | No value | 0 | 0 | 1 |
| 401 | W | 0 | 0 | No value | 0 | 0 | 1 |
| 402 | L | 0 | 0 | No value | 0 | 0 | 1 |
| 403 | N | 0 | 0 | No value | 0 | 0 | 1 |
| 404 | M | 0 | 1.049 | Infinity | 0.821 | 0.481 | 0.488 |
| 405 | L | 3.19 | 0 | 0 | 0.976 | 2.432 | 0.119 |
| 406 | Q | 3.346 | 0 | 0 | 0.925 | 2.415 | 0.12 |
| 407 | S | 2.976 | 0 | 0 | 0.977 | 2.263 | 0.132 |
| 408 | I | 4.96 | 0 | 0 | 1.081 | 3.138 | 0.076 |
| 409 | Q | 0 | 0 | No value | 0 | 0 | 1 |
| 410 | I | 0 | 0 | No value | 0 | 0 | 1 |
| 411 | P | 0 | 0 | No value | 0 | 0 | 1 |
| 412 | F | 7.043 | 0 | 0 | 1.235 | 3.153 | 0.076 |
| 413 | A | 0 | 0 | No value | 0 | 0 | 1 |
| 414 | L | 0 | 0 | No value | 0 | 0 | 1 |
| 415 | I | 1.756 | 0 | 0 | 0.727 | 1.745 | 0.187 |
| 416 | P | 0 | 0 | No value | 0 | 0 | 1 |
| 417 | L | 0 | 0 | No value | 0 | 0 | 1 |
| 418 | L | 2.005 | 0 | 0 | 0.801 | 1.819 | 0.177 |
| 419 | C | 0 | 0 | No value | 0 | 0 | 1 |
| 420 | L | 4.313 | 0 | 0 | 2.157 | 2.704 | 0.1 |
| 421 | V | 1.812 | 0 | 0 | 0.848 | 1.523 | 0.217 |
| 422 | S | 9.857 | 0 | 0 | 3.391 | 7.201 | 0.007 |
| 423 | K | 0 | 0 | No value | 0 | 0 | 1 |
| 424 | E | 0 | 0 | No value | 0 | 0 | 1 |
| 425 | Q | 0 | 0 | No value | 0 | 0 | 1 |
| 426 | . | 0 | 3.661 | Infinity | 1.629 | 2.777 | 0.096 |
| 427 | M | 0 | 0 | No value | 0 | 0 | 1 |

| | | | | | | | |
|---|---|---|---|---|---|---|---|
| 428 | G | 0 | 0 | No value | 0 | 0 | 1 |
| 429 | T | 0 | 0 | No value | 0 | 0 | 1 |
| 430 | F | 2.609 | 0 | 0 | 0.884 | 2.117 | 0.146 |
| 431 | T | 0 | 1.191 | Infinity | 0.776 | 0.845 | 0.358 |
| 432 | V | 4.701 | 1.598 | 0.34 | 2.906 | 0.757 | 0.384 |
| 433 | G | 1.633 | 0 | 0 | 0.733 | 1.596 | 0.206 |
| 434 | P | 0 | 0 | No value | 0 | 0 | 1 |
| 435 | I | 0 | 0 | No value | 0 | 0 | 1 |
| 436 | L | 0 | 0 | No value | 0 | 0 | 1 |
| 437 | . | 0 | 9.322 | Infinity | 4.902 | 4.004 | 0.045 |
| 438 | M | 0 | 0 | No value | 0 | 0 | 1 |
| 439 | V | 3.006 | 1.749 | 0.582 | 2.267 | 0.131 | 0.717 |
| 440 | S | 0 | 0 | No value | 0 | 0 | 1 |
| 441 | W | 0 | 0 | No value | 0 | 0 | 1 |
| 442 | L | 0 | 1.415 | Infinity | 0.937 | 0.794 | 0.373 |
| 443 | V | 1.906 | 0 | 0 | 0.867 | 1.569 | 0.21 |
| 444 | A | 0 | 0 | No value | 0 | 0 | 1 |
| 445 | A | 0 | 0 | No value | 0 | 0 | 1 |
| 446 | L | 1.492 | 0 | 0 | 0.869 | 1.068 | 0.301 |
| 447 | V | 0 | 0 | No value | 0 | 0 | 1 |
| 448 | M | 0 | 0 | No value | 0 | 0 | 1 |
| 449 | L | 1.52 | 2.286 | 1.504 | 1.832 | 0.076 | 0.782 |
| 450 | I | 0 | 0 | No value | 0 | 0 | 1 |
| 451 | N | 0 | 0 | No value | 0 | 0 | 1 |
| 452 | G | 0 | 0 | No value | 0 | 0 | 1 |
| 453 | Y | 0 | 0 | No value | 0 | 0 | 1 |
| 454 | L | 0 | 0 | No value | 0 | 0 | 1 |
| 455 | L | 5.033 | 0 | 0 | 2.219 | 2.971 | 0.085 |
| 456 | L | 3.702 | 0 | 0 | 1.206 | 2.991 | 0.084 |
| 457 | D | 0 | 0 | No value | 0 | 0 | 1 |
| 458 | F | 0 | 0 | No value | 0 | 0 | 1 |
| 459 | F | 0 | 0 | No value | 0 | 0 | 1 |
| 460 | S | 0 | 1.382 | Infinity | 0.872 | 0.904 | 0.342 |
| 461 | N | 0 | 0 | No value | 0 | 0 | 1 |
| 462 | E | 0 | 0 | No value | 0 | 0 | 1 |
| 463 | V | 3.97 | 0 | 0 | 1.814 | 3.142 | 0.076 |
| 464 | T | 4.204 | 0 | 0 | 1.453 | 4.217 | 0.04 |
| 465 | G | 0 | 0 | No value | 0 | 0 | 1 |
| 466 | V | 0 | 0 | No value | 0 | 0 | 1 |
| 467 | . | 2.088 | 6.752 | 3.234 | 4.289 | 1.055 | 0.304 |
| 468 | F | 0 | 0 | No value | 0 | 0 | 1 |
| 469 | T | 0 | 1.125 | Infinity | 0.614 | 1.165 | 0.28 |
| 470 | T | 3.168 | 0 | 0 | 0.789 | 2.68 | 0.102 |
| 471 | V | 1.908 | 0 | 0 | 0.867 | 1.571 | 0.21 |
| 472 | V | 0 | 0 | No value | 0 | 0 | 1 |
| 473 | C | 0 | 0 | No value | 0 | 0 | 1 |
| 474 | A | 0 | 3.212 | Infinity | 2.047 | 1.732 | 0.188 |
| 475 | F | 0 | 0 | No value | 0 | 0 | 1 |
| 476 | T | 0 | 0 | No value | 0 | 0 | 1 |

| | | | | | | | |
|---|---|---|---|---|---|---|---|
| 477 | G | 2.909 | 0 | 0 | 0.97 | 2.226 | 0.136 |
| 478 | A | 1.924 | 0 | 0 | 0.72 | 1.913 | 0.167 |
| 479 | Y | 7.755 | 0 | 0 | 1.312 | 3.329 | 0.068 |
| 480 | V | 3.204 | 0 | 0 | 1.057 | 2.154 | 0.142 |
| 481 | T | 1.95 | 2.479 | 1.271 | 2.271 | 0.035 | 0.852 |
| 482 | F | 0 | 0 | No value | 0 | 0 | 1 |
| 483 | I | 0 | 1.125 | Infinity | 0.863 | 0.491 | 0.484 |
| 484 | I | 0 | 0 | No value | 0 | 0 | 1 |
| 485 | Y | 0 | 0 | No value | 0 | 0 | 1 |
| 486 | L | 0 | 0 | No value | 0 | 0 | 1 |
| 487 | I | 4.444 | 0 | 0 | 0.994 | 2.84 | 0.092 |
| 488 | S | 0 | 0 | No value | 0 | 0 | 1 |
| 489 | R | 0 | 0 | No value | 0 | 0 | 1 |
| 490 | E | 0 | 2.675 | Infinity | 1.695 | 0.312 | 0.577 |
| 491 | V | 11.922 | 0 | 0 | 3.623 | 4.557 | 0.033 |
| 492 | . | 3.85 | 2.96 | 0.769 | 3.412 | 0.054 | 0.816 |
| 493 | I | 0 | 4.109 | Infinity | 2.808 | 1.54 | 0.215 |
| 494 | S | 0 | 1.402 | Infinity | 0.77 | 1.167 | 0.28 |
| 495 | T | 2.03 | 1.348 | 0.664 | 1.627 | 0.079 | 0.779 |
| 496 | W | 0 | 0 | No value | 0 | 0 | 1 |
| 497 | Y | 2.601 | 1.625 | 0.625 | 1.989 | 0.104 | 0.747 |
| 498 | C | 0 | 0 | No value | 0 | 0 | 1 |
| 499 | P | 3.96 | 0 | 0 | 1.364 | 4.185 | 0.041 |
| 500 | T | 6.648 | 3.08 | 0.463 | 4.616 | 0.633 | 0.426 |

**Supplementary table S3.** Sites under positive and purifying selection in *Populus NRAMP3.2* according to FEL method. Sites subjected to positive and purifying selections were estimated by the FEL method (https://www.datamonkey.org/) (Kosakovsky Pond and Frost 2005). *NRAMP3.2* coding sequences of 13 *Populus* species retrieved as indicated in Materials and Methods were used to generate a phylogenetic tree. A subset of branches encompassing the *NRAMP3.2s* were analyzed separately to estimate the synonymous substitution rate (dS) at a site (α) and the non-synonymous substitution rate (dN) at a site (β), and a maximum-likelihood approach was undertaken to calculate dN/dS rates for each codon site (ϖ). Amino-acids correspond to that of the consensus *Populus* NRAMP3.1 sequence. Dots indicate less than 90% of conservation within all investigated Populus NRAMP3.1 sequences. Sites subjected to positive selection are highlighted in dark ($p < 0.05$) and light green ($p \leq 0.1$). Sites subject to purifying selection are highlighted in dark ($p < 0.05$) and light red ($p < 0.1$). α=β : the rate estimated under the neutral model; LRT: likelihood ratio test.

| Site | amino-acid | α | β | ϖ | α=β | LRT | p-value |
|---|---|---|---|---|---|---|---|
| 1 | M | 0 | 0 | No value | 0 | 0 | 1 |
| 2 | P | 3.207 | 2.489 | 0.776 | 2.784 | 0.031 | 0.86 |
| 3 | V | 0 | 0 | No value | 0 | 0 | 1 |
| 4 | - | 0 | 0 | No value | 0 | 0 | 1 |
| 5 | E | 0 | 0 | No value | 0 | 0 | 1 |
| 6 | E | 0 | 0 | No value | 0 | 0 | 1 |
| 7 | N | 0 | 0 | No value | 0 | 0 | 1 |
| 8 | . | 5.82 | 5.527 | 0.95 | 5.656 | 0.002 | 0.964 |
| 9 | Q | 0 | 0 | No value | 0 | 0 | 1 |
| 10 | P | 3.162 | 2.493 | 0.788 | 2.774 | 0.025 | 0.875 |
| 11 | L | 0 | 0 | No value | 0 | 0 | 1 |
| 12 | L | 2.026 | 0 | 0 | 1.417 | 0.701 | 0.402 |
| 13 | Q | 0 | 0 | No value | 0 | 0 | 1 |
| 14 | E | 0 | 0 | No value | 0 | 0 | 1 |
| 15 | E | 0 | 0 | No value | 0 | 0 | 1 |
| 16 | E | 0 | 0 | No value | 0 | 0 | 1 |
| 17 | E | 0 | 0 | No value | 0 | 0 | 1 |
| 18 | R | 0 | 0 | No value | 0 | 0 | 1 |
| 19 | A | 0 | 0 | No value | 0 | 0 | 1 |
| 20 | Y | 0 | 0 | No value | 0 | 0 | 1 |
| 21 | D | 0 | 0 | No value | 0 | 0 | 1 |
| 22 | S | 0 | 0 | No value | 0 | 0 | 1 |
| 23 | D | 3.393 | 0 | 0 | 1.482 | 1.566 | 0.211 |
| 24 | E | 0 | 0 | No value | 0 | 0 | 1 |
| 25 | K | 0 | 0 | No value | 0 | 0 | 1 |
| 26 | V | 4.189 | 0 | 0 | 2.641 | 1.786 | 0.181 |
| 27 | L | 2.005 | 0 | 0 | 1.21 | 0.984 | 0.321 |
| 28 | I | 2.724 | 0 | 0 | 1.271 | 1.46 | 0.227 |
| 29 | I | 0 | 0 | No value | 0 | 0 | 1 |
| 30 | G | 1.966 | 0 | 0 | 1.184 | 0.972 | 0.324 |
| 31 | V | 3.598 | 0 | 0 | 1.756 | 1.324 | 0.25 |
| 32 | D | 0 | 0 | No value | 0 | 0 | 1 |
| 33 | S | 0 | 0 | No value | 0 | 0 | 1 |
| 34 | D | 0 | 0 | No value | 0 | 0 | 1 |
| 35 | T | 9.333 | 0 | 0 | 4.662 | 5.081 | 0.024 |
| 36 | E | 0 | 0 | No value | 0 | 0 | 1 |

| | | | | | | | |
|---|---|---|---|---|---|---|---|
| 37 | S | 2.649 | 0 | 0 | 1.186 | 1.54 | 0.215 |
| 38 | G | 0 | 3.103 | Infinity | 1.507 | 1.453 | 0.228 |
| 39 | G | 0 | 3.139 | Infinity | 0.976 | 2.329 | 0.127 |
| 40 | S | 0 | 0 | No value | 0 | 0 | 1 |
| 41 | T | 0 | 0 | No value | 0 | 0 | 1 |
| 42 | V | 0 | 0 | No value | 0 | 0 | 1 |
| 43 | L | 0 | 0 | No value | 0 | 0 | 1 |
| 44 | P | 1.903 | 0 | 0 | 1.049 | 1.151 | 0.283 |
| 45 | P | 2.32 | 0 | 0 | 1.171 | 1.333 | 0.248 |
| 46 | F | 0 | 0 | No value | 0 | 0 | 1 |
| 47 | S | 0 | 0 | No value | 0 | 0 | 1 |
| 48 | W | 0 | 0 | No value | 0 | 0 | 1 |
| 49 | K | 0 | 0 | No value | 0 | 0 | 1 |
| 50 | K | 4.546 | 0 | 0 | 1.638 | 1.963 | 0.161 |
| 51 | L | 0 | 0 | No value | 0 | 0 | 1 |
| 52 | W | 0 | 0 | No value | 0 | 0 | 1 |
| 53 | L | 2.183 | 0 | 0 | 1.467 | 0.728 | 0.394 |
| 54 | F | 5.723 | 0 | 0 | 2.298 | 1.919 | 0.166 |
| 55 | T | 0 | 0 | No value | 0 | 0 | 1 |
| 56 | G | 0 | 0 | No value | 0 | 0 | 1 |
| 57 | P | 0 | 0 | No value | 0 | 0 | 1 |
| 58 | G | 1.949 | 0 | 0 | 1.178 | 0.961 | 0.327 |
| 59 | F | 0 | 0 | No value | 0 | 0 | 1 |
| 60 | L | 0 | 0 | No value | 0 | 0 | 1 |
| 61 | M | 0 | 0 | No value | 0 | 0 | 1 |
| 62 | S | 1.704 | 0 | 0 | 1.088 | 0.856 | 0.355 |
| 63 | I | 0 | 0 | No value | 0 | 0 | 1 |
| 64 | A | 3.044 | 0 | 0 | 1.444 | 1.501 | 0.221 |
| 65 | F | 0 | 0 | No value | 0 | 0 | 1 |
| 66 | L | 0 | 0 | No value | 0 | 0 | 1 |
| 67 | D | 3.407 | 0 | 0 | 1.505 | 1.58 | 0.209 |
| 68 | P | 0 | 0 | No value | 0 | 0 | 1 |
| 69 | G | 6.296 | 0 | 0 | 3.728 | 3.013 | 0.083 |
| 70 | N | 0 | 0 | No value | 0 | 0 | 1 |
| 71 | L | 0 | 0 | No value | 0 | 0 | 1 |
| 72 | E | 0 | 0 | No value | 0 | 0 | 1 |
| 73 | G | 0 | 0 | No value | 0 | 0 | 1 |
| 74 | D | 0 | 0 | No value | 0 | 0 | 1 |
| 75 | L | 0 | 0 | No value | 0 | 0 | 1 |
| 76 | Q | 0 | 0 | No value | 0 | 0 | 1 |
| 77 | A | 0 | 0 | No value | 0 | 0 | 1 |
| 78 | G | 0 | 0 | No value | 0 | 0 | 1 |
| 79 | A | 1.979 | 0 | 0 | 1.121 | 1.071 | 0.301 |
| 80 | I | 2.45 | 0 | 0 | 1.318 | 1.215 | 0.27 |
| 81 | A | 1.942 | 0 | 0 | 1.114 | 1.063 | 0.303 |
| 82 | G | 6.48 | 0 | 0 | 3.774 | 3.104 | 0.078 |
| 83 | Y | 3.553 | 0 | 0 | 1.793 | 1.314 | 0.252 |
| 84 | S | 0 | 0 | No value | 0 | 0 | 1 |

| | | | | | | | |
|---|---|---|---|---|---|---|---|
| 85 | L | 0 | 0 | No value | 0 | 0 | 1 |
| 86 | L | 0 | 0 | No value | 0 | 0 | 1 |
| 87 | W | 0 | 0 | No value | 0 | 0 | 1 |
| 88 | L | 0 | 0 | No value | 0 | 0 | 1 |
| 89 | L | 1.929 | 0 | 0 | 1.181 | 0.954 | 0.329 |
| 90 | L | 0 | 4.902 | Infinity | 1.492 | 2.417 | 0.12 |
| 91 | W | 0 | 0 | No value | 0 | 0 | 1 |
| 92 | A | 0 | 0 | No value | 0 | 0 | 1 |
| 93 | T | 0 | 0 | No value | 0 | 0 | 1 |
| 94 | A | 0 | 0 | No value | 0 | 0 | 1 |
| 95 | M | 0 | 0 | No value | 0 | 0 | 1 |
| 96 | G | 6.514 | 0 | 0 | 3.172 | 2.834 | 0.092 |
| 97 | L | 0 | 0 | No value | 0 | 0 | 1 |
| 98 | L | 0 | 0 | No value | 0 | 0 | 1 |
| 99 | V | 0 | 0 | No value | 0 | 0 | 1 |
| 100 | Q | 0 | 0 | No value | 0 | 0 | 1 |
| 101 | L | 0 | 0 | No value | 0 | 0 | 1 |
| 102 | L | 1.996 | 0 | 0 | 1.211 | 0.982 | 0.322 |
| 103 | S | 0 | 0 | No value | 0 | 0 | 1 |
| 104 | A | 3.973 | 0 | 0 | 2.267 | 2.155 | 0.142 |
| 105 | R | 0 | 0 | No value | 0 | 0 | 1 |
| 106 | L | 0 | 0 | No value | 0 | 0 | 1 |
| 107 | G | 1.927 | 0 | 0 | 1.2 | 0.889 | 0.346 |
| 108 | V | 0 | 0 | No value | 0 | 0 | 1 |
| 109 | A | 0 | 0 | No value | 0 | 0 | 1 |
| 110 | T | 0 | 0 | No value | 0 | 0 | 1 |
| 111 | G | 3.292 | 0 | 0 | 1.787 | 1.522 | 0.217 |
| 112 | R | 2.398 | 0 | 0 | 1.216 | 1.271 | 0.26 |
| 113 | H | 0 | 0 | No value | 0 | 0 | 1 |
| 114 | L | 4.346 | 0 | 0 | 2.942 | 1.433 | 0.231 |
| 115 | A | 0 | 0 | No value | 0 | 0 | 1 |
| 116 | E | 0 | 0 | No value | 0 | 0 | 1 |
| 117 | L | 1.165 | 0 | 0 | 0.925 | 0.461 | 0.497 |
| 118 | C | 0 | 0 | No value | 0 | 0 | 1 |
| 119 | R | 0 | 0 | No value | 0 | 0 | 1 |
| 120 | E | 0 | 0 | No value | 0 | 0 | 1 |
| 121 | E | 0 | 0 | No value | 0 | 0 | 1 |
| 122 | Y | 0 | 0 | No value | 0 | 0 | 1 |
| 123 | P | 0 | 0 | No value | 0 | 0 | 1 |
| 124 | T | 4.515 | 0 | 0 | 2.239 | 2.464 | 0.116 |
| 125 | W | 0 | 0 | No value | 0 | 0 | 1 |
| 126 | A | 3.245 | 0 | 0 | 1.433 | 1.521 | 0.217 |
| 127 | S | 0 | 0 | No value | 0 | 0 | 1 |
| 128 | M | 0 | 0 | No value | 0 | 0 | 1 |
| 129 | V | 0 | 3.806 | Infinity | 1.759 | 1.546 | 0.214 |
| 130 | L | 0 | 0 | No value | 0 | 0 | 1 |
| 131 | W | 0 | 0 | No value | 0 | 0 | 1 |
| 132 | I | 0 | 0 | No value | 0 | 0 | 1 |

| | | | | | | | |
|---|---|---|---|---|---|---|---|
| 133 | M | 0 | 0 | No value | 0 | 0 | 1 |
| 134 | A | 0 | 2.754 | Infinity | 1.415 | 1.356 | 0.244 |
| 135 | E | 0 | 0 | No value | 0 | 0 | 1 |
| 136 | L | 0 | 0 | No value | 0 | 0 | 1 |
| 137 | A | 0 | 0 | No value | 0 | 0 | 1 |
| 138 | L | 0 | 0 | No value | 0 | 0 | 1 |
| 139 | I | 0 | 0 | No value | 0 | 0 | 1 |
| 140 | G | 0 | 0 | No value | 0 | 0 | 1 |
| 141 | A | 0 | 0 | No value | 0 | 0 | 1 |
| 142 | D | 0 | 0 | No value | 0 | 0 | 1 |
| 143 | I | 0 | 0 | No value | 0 | 0 | 1 |
| 144 | Q | 0 | 0 | No value | 0 | 0 | 1 |
| 145 | E | 3.161 | 0 | 0 | 1.486 | 1.454 | 0.228 |
| 146 | V | 0 | 0 | No value | 0 | 0 | 1 |
| 147 | I | 0 | 0 | No value | 0 | 0 | 1 |
| 148 | G | 1.908 | 0 | 0 | 1.207 | 0.889 | 0.346 |
| 149 | S | 0 | 0 | No value | 0 | 0 | 1 |
| 150 | A | 0 | 0 | No value | 0 | 0 | 1 |
| 151 | I | 14.276 | 0 | 0 | 4.079 | 4.641 | 0.031 |
| 152 | A | 3.118 | 0 | 0 | 1.415 | 1.49 | 0.222 |
| 153 | . | 0 | 3.19 | Infinity | 1.692 | 1.281 | 0.258 |
| 154 | K | 0 | 0 | No value | 0 | 0 | 1 |
| 155 | I | 2.45 | 0 | 0 | 1.318 | 1.215 | 0.27 |
| 156 | L | 0 | 0 | No value | 0 | 0 | 1 |
| 157 | S | 0 | 0 | No value | 0 | 0 | 1 |
| 158 | N | 0 | 0 | No value | 0 | 0 | 1 |
| 159 | G | 0 | 0 | No value | 0 | 0 | 1 |
| 160 | F | 0 | 0 | No value | 0 | 0 | 1 |
| 161 | . | 0 | 8.927 | Infinity | 2.985 | 4.488 | 0.034 |
| 162 | P | 0 | 0 | No value | 0 | 0 | 1 |
| 163 | L | 2.038 | 0 | 0 | 1.401 | 0.692 | 0.405 |
| 164 | W | 0 | 0 | No value | 0 | 0 | 1 |
| 165 | A | 0 | 0 | No value | 0 | 0 | 1 |
| 166 | G | 0 | 0 | No value | 0 | 0 | 1 |
| 167 | V | 0 | 0 | No value | 0 | 0 | 1 |
| 168 | T | 0 | 0 | No value | 0 | 0 | 1 |
| 169 | I | 0 | 0 | No value | 0 | 0 | 1 |
| 170 | T | 0 | 0 | No value | 0 | 0 | 1 |
| 171 | A | 3.028 | 0 | 0 | 1.394 | 1.455 | 0.228 |
| 172 | C | 4.936 | 0 | 0 | 2.781 | 2.162 | 0.141 |
| 173 | D | 0 | 0 | No value | 0 | 0 | 1 |
| 174 | . | 0 | 6.937 | Infinity | 3.048 | 3.355 | 0.067 |
| 175 | F | 3.553 | 0 | 0 | 1.642 | 1.487 | 0.223 |
| 176 | I | 1.775 | 0 | 0 | 1.086 | 0.941 | 0.332 |
| 177 | F | 0 | 0 | No value | 0 | 0 | 1 |
| 178 | L | 0 | 0 | No value | 0 | 0 | 1 |
| 179 | F | 0 | 0 | No value | 0 | 0 | 1 |
| 180 | L | 2.072 | 0 | 0 | 1.419 | 0.757 | 0.384 |

| | | | | | | | |
|---|---|---|---|---|---|---|---|
| 181 | E | 0 | 0 | No value | 0 | 0 | 1 |
| 182 | N | 0 | 0 | No value | 0 | 0 | 1 |
| 183 | Y | 0 | 0 | No value | 0 | 0 | 1 |
| 184 | G | 3.106 | 0 | 0 | 1.509 | 1.358 | 0.244 |
| 185 | V | 0 | 0 | No value | 0 | 0 | 1 |
| 186 | R | 2.349 | 0 | 0 | 1.242 | 1.229 | 0.268 |
| 187 | K | 0 | 0 | No value | 0 | 0 | 1 |
| 188 | L | 2.025 | 0 | 0 | 1.399 | 0.695 | 0.404 |
| 189 | E | 0 | 0 | No value | 0 | 0 | 1 |
| 190 | A | 0 | 0 | No value | 0 | 0 | 1 |
| 191 | V | 2.973 | 0 | 0 | 1.65 | 1.186 | 0.276 |
| 192 | F | 0 | 0 | No value | 0 | 0 | 1 |
| 193 | A | 0 | 0 | No value | 0 | 0 | 1 |
| 194 | V | 9.042 | 0 | 0 | 4.683 | 3.359 | 0.067 |
| 195 | L | 1.929 | 0 | 0 | 1.181 | 0.954 | 0.329 |
| 196 | I | 0 | 0 | No value | 0 | 0 | 1 |
| 197 | G | 0 | 0 | No value | 0 | 0 | 1 |
| 198 | I | 0 | 2.513 | Infinity | 1.578 | 0.935 | 0.334 |
| 199 | M | 0 | 0 | No value | 0 | 0 | 1 |
| 200 | A | 0 | 0 | No value | 0 | 0 | 1 |
| 201 | V | 2.955 | 0 | 0 | 1.642 | 1.185 | 0.276 |
| 202 | T | 0 | 0 | No value | 0 | 0 | 1 |
| 203 | F | 0 | 0 | No value | 0 | 0 | 1 |
| 204 | G | 1.912 | 0 | 0 | 1.209 | 0.89 | 0.345 |
| 205 | W | 0 | 0 | No value | 0 | 0 | 1 |
| 206 | M | 0 | 0 | No value | 0 | 0 | 1 |
| 207 | F | 0 | 0 | No value | 0 | 0 | 1 |
| 208 | A | 2.996 | 2.985 | 0.996 | 2.975 | 0 | 1 |
| 209 | D | 0 | 0 | No value | 0 | 0 | 1 |
| 210 | A | 0 | 0 | No value | 0 | 0 | 1 |
| 211 | K | 0 | 0 | No value | 0 | 0 | 1 |
| 212 | P | 0 | 0 | No value | 0 | 0 | 1 |
| 213 | S | 0 | 0 | No value | 0 | 0 | 1 |
| 214 | A | 1.426 | 0 | 0 | 0.932 | 0.841 | 0.359 |
| 215 | S | 3.9 | 0 | 0 | 2.316 | 1.879 | 0.17 |
| 216 | E | 0 | 0 | No value | 0 | 0 | 1 |
| 217 | L | 3.133 | 0 | 0 | 1.52 | 1.352 | 0.245 |
| 218 | F | 0 | 0 | No value | 0 | 0 | 1 |
| 219 | L | 7.839 | 0 | 0 | 5.407 | 2.815 | 0.093 |
| 220 | G | 1.943 | 0 | 0 | 1.18 | 0.963 | 0.327 |
| 221 | I | 1.744 | 0 | 0 | 1.079 | 0.935 | 0.334 |
| 222 | L | 0 | 0 | No value | 0 | 0 | 1 |
| 223 | I | 0 | 0 | No value | 0 | 0 | 1 |
| 224 | P | 1.881 | 0 | 0 | 1.037 | 1.134 | 0.287 |
| 225 | K | 0 | 0 | No value | 0 | 0 | 1 |
| 226 | L | 0 | 0 | No value | 0 | 0 | 1 |
| 227 | S | 0 | 0 | No value | 0 | 0 | 1 |
| 228 | S | 0 | 0 | No value | 0 | 0 | 1 |

| | | | | | | | |
|---|---|---|---|---|---|---|---|
| 229 | R | 0 | 0 | No value | 0 | 0 | 1 |
| 230 | T | 0 | 0 | No value | 0 | 0 | 1 |
| 231 | I | 0 | 0 | No value | 0 | 0 | 1 |
| 232 | Q | 0 | 0 | No value | 0 | 0 | 1 |
| 233 | Q | 3.606 | 0 | 0 | 1.603 | 1.484 | 0.223 |
| 234 | A | 3.324 | 0 | 0 | 1.455 | 1.554 | 0.213 |
| 235 | V | 9.822 | 0 | 0 | 4.376 | 3.413 | 0.065 |
| 236 | G | 0 | 0 | No value | 0 | 0 | 1 |
| 237 | V | 0 | 0 | No value | 0 | 0 | 1 |
| 238 | V | 0 | 0 | No value | 0 | 0 | 1 |
| 239 | G | 0 | 0 | No value | 0 | 0 | 1 |
| 240 | C | 0 | 0 | No value | 0 | 0 | 1 |
| 241 | I | 0 | 0 | No value | 0 | 0 | 1 |
| 242 | I | 0 | 0 | No value | 0 | 0 | 1 |
| 243 | M | 0 | 0 | No value | 0 | 0 | 1 |
| 244 | P | 0 | 0 | No value | 0 | 0 | 1 |
| 245 | H | 0 | 0 | No value | 0 | 0 | 1 |
| 246 | N | 0 | 0 | No value | 0 | 0 | 1 |
| 247 | V | 0 | 0 | No value | 0 | 0 | 1 |
| 248 | F | 0 | 0 | No value | 0 | 0 | 1 |
| 249 | L | 0 | 0 | No value | 0 | 0 | 1 |
| 250 | H | 0 | 0 | No value | 0 | 0 | 1 |
| 251 | S | 0 | 0 | No value | 0 | 0 | 1 |
| 252 | A | 0 | 0 | No value | 0 | 0 | 1 |
| 253 | L | 0 | 0 | No value | 0 | 0 | 1 |
| 254 | V | 1.917 | 0 | 0 | 1.237 | 0.834 | 0.361 |
| 255 | Q | 0 | 0 | No value | 0 | 0 | 1 |
| 256 | S | 0 | 0 | No value | 0 | 0 | 1 |
| 257 | R | 0 | 0 | No value | 0 | 0 | 1 |
| 258 | E | 0 | 0 | No value | 0 | 0 | 1 |
| 259 | I | 12.387 | 0 | 0 | 5.669 | 5.406 | 0.02 |
| 260 | D | 0 | 0 | No value | 0 | 0 | 1 |
| 261 | H | 0 | 0 | No value | 0 | 0 | 1 |
| 262 | N | 0 | 2.549 | Infinity | 1.747 | 0.791 | 0.374 |
| 263 | K | 0 | 0 | No value | 0 | 0 | 1 |
| 264 | K | 0 | 0 | No value | 0 | 0 | 1 |
| 265 | . | 0 | 9.06 | Infinity | 2.311 | 4.965 | 0.026 |
| 266 | R | 1.963 | 0 | 0 | 1.191 | 0.943 | 0.332 |
| 267 | V | 0 | 0 | No value | 0 | 0 | 1 |
| 268 | Q | 0 | 0 | No value | 0 | 0 | 1 |
| 269 | E | 0 | 0 | No value | 0 | 0 | 1 |
| 270 | A | 0 | 0 | No value | 0 | 0 | 1 |
| 271 | L | 1.451 | 0 | 0 | 0.983 | 0.759 | 0.384 |
| 272 | R | 0 | 0 | No value | 0 | 0 | 1 |
| 273 | Y | 0 | 0 | No value | 0 | 0 | 1 |
| 274 | Y | 0 | 0 | No value | 0 | 0 | 1 |
| 275 | S | 0 | 0 | No value | 0 | 0 | 1 |
| 276 | I | 0 | 0 | No value | 0 | 0 | 1 |

| | | | | | | | |
|---|---|---|---|---|---|---|---|
| 277 | E | 0 | 0 | No value | 0 | 0 | 1 |
| 278 | S | 0 | 0 | No value | 0 | 0 | 1 |
| 279 | T | 2.004 | 0 | 0 | 1.102 | 1.174 | 0.278 |
| 280 | T | 0 | 0 | No value | 0 | 0 | 1 |
| 281 | A | 0 | 0 | No value | 0 | 0 | 1 |
| 282 | L | 0 | 0 | No value | 0 | 0 | 1 |
| 283 | V | 0 | 0 | No value | 0 | 0 | 1 |
| 284 | I | 0 | 0 | No value | 0 | 0 | 1 |
| 285 | S | 6.339 | 0 | 0 | 3.928 | 2.601 | 0.107 |
| 286 | F | 0 | 0 | No value | 0 | 0 | 1 |
| 287 | V | 2.025 | 4.376 | 2.161 | 3.372 | 0.154 | 0.695 |
| 288 | I | 0 | 0 | No value | 0 | 0 | 1 |
| 289 | N | 0 | 0 | No value | 0 | 0 | 1 |
| 290 | L | 0 | 0 | No value | 0 | 0 | 1 |
| 291 | F | 0 | 0 | No value | 0 | 0 | 1 |
| 292 | V | 0 | 0 | No value | 0 | 0 | 1 |
| 293 | T | 4.524 | 0 | 0 | 2.278 | 2.497 | 0.114 |
| 294 | T | 6.503 | 0 | 0 | 3.37 | 3.629 | 0.057 |
| 295 | V | 3.263 | 0 | 0 | 1.378 | 1.142 | 0.285 |
| 296 | F | 0 | 0 | No value | 0 | 0 | 1 |
| 297 | A | 0 | 0 | No value | 0 | 0 | 1 |
| 298 | K | 0 | 0 | No value | 0 | 0 | 1 |
| 299 | G | 0 | 0 | No value | 0 | 0 | 1 |
| 300 | F | 6.037 | 0 | 0 | 2.912 | 2.56 | 0.11 |
| 301 | Y | 8.263 | 0 | 0 | 3.57 | 2.706 | 0.1 |
| 302 | G | 0 | 0 | No value | 0 | 0 | 1 |
| 303 | T | 0 | 0 | No value | 0 | 0 | 1 |
| 304 | E | 0 | 0 | No value | 0 | 0 | 1 |
| 305 | L | 1.178 | 0 | 0 | 0.925 | 0.462 | 0.497 |
| 306 | A | 0 | 0 | No value | 0 | 0 | 1 |
| 307 | N | 0 | 0 | No value | 0 | 0 | 1 |
| 308 | S | 0 | 0 | No value | 0 | 0 | 1 |
| 309 | I | 0 | 0 | No value | 0 | 0 | 1 |
| 310 | G | 0 | 0 | No value | 0 | 0 | 1 |
| 311 | L | 0 | 0 | No value | 0 | 0 | 1 |
| 312 | V | 0 | 0 | No value | 0 | 0 | 1 |
| 313 | N | 0 | 0 | No value | 0 | 0 | 1 |
| 314 | A | 0 | 0 | No value | 0 | 0 | 1 |
| 315 | G | 0 | 0 | No value | 0 | 0 | 1 |
| 316 | Q | 0 | 0 | No value | 0 | 0 | 1 |
| 317 | Y | 0 | 0 | No value | 0 | 0 | 1 |
| 318 | L | 0 | 0 | No value | 0 | 0 | 1 |
| 319 | Q | 3.691 | 0 | 0 | 1.578 | 1.543 | 0.214 |
| 320 | D | 7.486 | 0 | 0 | 2.899 | 3.57 | 0.059 |
| 321 | K | 0 | 0 | No value | 0 | 0 | 1 |
| 322 | Y | 0 | 0 | No value | 0 | 0 | 1 |
| 323 | G | 1.951 | 0 | 0 | 1.174 | 0.962 | 0.327 |
| 324 | G | 0 | 0 | No value | 0 | 0 | 1 |

| | | | | | | | |
|---|---|---|---|---|---|---|---|
| 325 | G | 1.91 | 0 | 0 | 1.201 | 0.888 | 0.346 |
| 326 | F | 0 | 0 | No value | 0 | 0 | 1 |
| 327 | F | 0 | 0 | No value | 0 | 0 | 1 |
| 328 | P | 0 | 0 | No value | 0 | 0 | 1 |
| 329 | I | 0 | 0 | No value | 0 | 0 | 1 |
| 330 | L | 0 | 0 | No value | 0 | 0 | 1 |
| 331 | Y | 0 | 0 | No value | 0 | 0 | 1 |
| 332 | I | 0 | 0 | No value | 0 | 0 | 1 |
| 333 | W | 0 | 0 | No value | 0 | 0 | 1 |
| 334 | G | 3.127 | 0 | 0 | 1.508 | 1.357 | 0.244 |
| 335 | I | 4.186 | 0 | 0 | 1.685 | 1.671 | 0.196 |
| 336 | G | 1.908 | 0 | 0 | 1.207 | 0.889 | 0.346 |
| 337 | L | 0 | 0 | No value | 0 | 0 | 1 |
| 338 | L | 0 | 0 | No value | 0 | 0 | 1 |
| 339 | A | 0 | 0 | No value | 0 | 0 | 1 |
| 340 | A | 0 | 0 | No value | 0 | 0 | 1 |
| 341 | G | 0 | 0 | No value | 0 | 0 | 1 |
| 342 | Q | 0 | 0 | No value | 0 | 0 | 1 |
| 343 | S | 7.48 | 0 | 0 | 2.72 | 3.815 | 0.051 |
| 344 | S | 0 | 0 | No value | 0 | 0 | 1 |
| 345 | T | 0 | 0 | No value | 0 | 0 | 1 |
| 346 | I | 0 | 0 | No value | 0 | 0 | 1 |
| 347 | T | 0 | 0 | No value | 0 | 0 | 1 |
| 348 | G | 1.454 | 0 | 0 | 0.984 | 0.768 | 0.381 |
| 349 | T | 3.129 | 0 | 0 | 1.345 | 1.582 | 0.208 |
| 350 | Y | 0 | 0 | No value | 0 | 0 | 1 |
| 351 | A | 0 | 0 | No value | 0 | 0 | 1 |
| 352 | G | 3.997 | 0 | 0 | 2.466 | 1.837 | 0.175 |
| 353 | Q | 0 | 0 | No value | 0 | 0 | 1 |
| 354 | F | 0 | 0 | No value | 0 | 0 | 1 |
| 355 | I | 0 | 0 | No value | 0 | 0 | 1 |
| 356 | M | 0 | 0 | No value | 0 | 0 | 1 |
| 357 | G | 0 | 0 | No value | 0 | 0 | 1 |
| 358 | G | 0 | 0 | No value | 0 | 0 | 1 |
| 359 | F | 0 | 0 | No value | 0 | 0 | 1 |
| 360 | L | 0 | 0 | No value | 0 | 0 | 1 |
| 361 | N | 0 | 0 | No value | 0 | 0 | 1 |
| 362 | L | 0 | 0 | No value | 0 | 0 | 1 |
| 363 | R | 5.852 | 0 | 0 | 2.731 | 2.948 | 0.086 |
| 364 | L | 0 | 0 | No value | 0 | 0 | 1 |
| 365 | K | 0 | 0 | No value | 0 | 0 | 1 |
| 366 | K | 0 | 0 | No value | 0 | 0 | 1 |
| 367 | W | 0 | 0 | No value | 0 | 0 | 1 |
| 368 | L | 4.046 | 0 | 0 | 3.066 | 1.509 | 0.219 |
| 369 | R | 0 | 0 | No value | 0 | 0 | 1 |
| 370 | A | 0 | 0 | No value | 0 | 0 | 1 |
| 371 | L | 2.048 | 0 | 0 | 1.403 | 0.694 | 0.405 |
| 372 | I | 2.45 | 0 | 0 | 1.318 | 1.215 | 0.27 |

| | | | | | | | |
|---|---|---|---|---|---|---|---|
| 373 | T | 0 | 0 | No value | 0 | 0 | 1 |
| 374 | R | 0 | 0 | No value | 0 | 0 | 1 |
| 375 | S | 0 | 0 | No value | 0 | 0 | 1 |
| 376 | C | 0 | 0 | No value | 0 | 0 | 1 |
| 377 | A | 0 | 0 | No value | 0 | 0 | 1 |
| 378 | I | 1.758 | 0 | 0 | 1.085 | 0.94 | 0.332 |
| 379 | I | 0 | 0 | No value | 0 | 0 | 1 |
| 380 | P | 0 | 0 | No value | 0 | 0 | 1 |
| 381 | T | 0 | 0 | No value | 0 | 0 | 1 |
| 382 | M | 0 | 0 | No value | 0 | 0 | 1 |
| 383 | I | 0 | 0 | No value | 0 | 0 | 1 |
| 384 | V | 0 | 0 | No value | 0 | 0 | 1 |
| 385 | A | 1.925 | 0 | 0 | 1.104 | 1.05 | 0.306 |
| 386 | L | 0 | 0 | No value | 0 | 0 | 1 |
| 387 | V | 3.186 | 0 | 0 | 1.656 | 1.225 | 0.268 |
| 388 | F | 0 | 0 | No value | 0 | 0 | 1 |
| 389 | D | 0 | 0 | No value | 0 | 0 | 1 |
| 390 | . | 1.994 | 5.36 | 2.688 | 3.569 | 0.63 | 0.427 |
| 391 | S | 7.301 | 0 | 0 | 4.004 | 3.278 | 0.07 |
| 392 | E | 0 | 0 | No value | 0 | 0 | 1 |
| 393 | D | 0 | 0 | No value | 0 | 0 | 1 |
| 394 | S | 1.925 | 0 | 0 | 1.287 | 0.752 | 0.386 |
| 395 | L | 0 | 0 | No value | 0 | 0 | 1 |
| 396 | D | 0 | 0 | No value | 0 | 0 | 1 |
| 397 | V | 0 | 0 | No value | 0 | 0 | 1 |
| 398 | L | 0 | 0 | No value | 0 | 0 | 1 |
| 399 | N | 0 | 0 | No value | 0 | 0 | 1 |
| 400 | E | 0 | 0 | No value | 0 | 0 | 1 |
| 401 | W | 0 | 0 | No value | 0 | 0 | 1 |
| 402 | L | 0 | 0 | No value | 0 | 0 | 1 |
| 403 | N | 0 | 0 | No value | 0 | 0 | 1 |
| 404 | V | 0 | 0 | No value | 0 | 0 | 1 |
| 405 | L | 3.19 | 0 | 0 | 1.832 | 1.133 | 0.287 |
| 406 | Q | 3.346 | 0 | 0 | 1.544 | 1.4 | 0.237 |
| 407 | S | 2.976 | 0 | 0 | 1.742 | 1.085 | 0.298 |
| 408 | I | 4.96 | 0 | 0 | 1.675 | 2.215 | 0.137 |
| 409 | Q | 0 | 0 | No value | 0 | 0 | 1 |
| 410 | I | 0 | 0 | No value | 0 | 0 | 1 |
| 411 | P | 0 | 0 | No value | 0 | 0 | 1 |
| 412 | F | 7.043 | 0 | 0 | 2.205 | 1.987 | 0.159 |
| 413 | A | 0 | 0 | No value | 0 | 0 | 1 |
| 414 | L | 0 | 0 | No value | 0 | 0 | 1 |
| 415 | I | 1.756 | 0 | 0 | 1.083 | 0.937 | 0.333 |
| 416 | P | 0 | 0 | No value | 0 | 0 | 1 |
| 417 | L | 0 | 0 | No value | 0 | 0 | 1 |
| 418 | L | 2.005 | 0 | 0 | 1.21 | 0.984 | 0.321 |
| 419 | C | 0 | 3.379 | Infinity | 1.428 | 1.728 | 0.189 |
| 420 | L | 4.313 | 0 | 0 | 2.94 | 1.438 | 0.23 |

| | | | | | | | |
|---|---|---|---|---|---|---|---|
| 421 | V | 1.812 | 0 | 0 | 1.205 | 0.803 | 0.37 |
| 422 | S | 9.857 | 0 | 0 | 4.875 | 4.219 | 0.04 |
| 423 | K | 0 | 0 | No value | 0 | 0 | 1 |
| 424 | E | 0 | 0 | No value | 0 | 0 | 1 |
| 425 | Q | 0 | 0 | No value | 0 | 0 | 1 |
| 426 | I | 0 | 0 | No value | 0 | 0 | 1 |
| 427 | M | 0 | 0 | No value | 0 | 0 | 1 |
| 428 | G | 0 | 0 | No value | 0 | 0 | 1 |
| 429 | T | 0 | 0 | No value | 0 | 0 | 1 |
| 430 | F | 2.609 | 0 | 0 | 1.399 | 1.192 | 0.275 |
| 431 | K | 0 | 0 | No value | 0 | 0 | 1 |
| 432 | I | 4.701 | 0 | 0 | 2.555 | 2.309 | 0.129 |
| 433 | G | 1.633 | 3.132 | 1.918 | 2.168 | 0.202 | 0.653 |
| 434 | P | 0 | 0 | No value | 0 | 0 | 1 |
| 435 | I | 0 | 3.074 | Infinity | 1.681 | 1.235 | 0.266 |
| 436 | L | 0 | 0 | No value | 0 | 0 | 1 |
| 437 | . | 0 | 6.063 | Infinity | 3.151 | 2.719 | 0.099 |
| 438 | M | 0 | 0 | No value | 0 | 0 | 1 |
| 439 | V | 3.006 | 0 | 0 | 1.66 | 1.196 | 0.274 |
| 440 | . | 0 | 6.576 | Infinity | 3.174 | 2.974 | 0.085 |
| 441 | W | 0 | 0 | No value | 0 | 0 | 1 |
| 442 | L | 0 | 0 | No value | 0 | 0 | 1 |
| 443 | V | 1.906 | 0 | 0 | 1.242 | 0.832 | 0.362 |
| 444 | A | 0 | 0 | No value | 0 | 0 | 1 |
| 445 | A | 0 | 0 | No value | 0 | 0 | 1 |
| 446 | L | 1.492 | 0 | 0 | 1.113 | 0.57 | 0.45 |
| 447 | V | 0 | 0 | No value | 0 | 0 | 1 |
| 448 | . | 0 | 4.623 | Infinity | 4.382 | 0.225 | 0.635 |
| 449 | V | 1.52 | 0 | 0 | 1.062 | 0.691 | 0.406 |
| 450 | I | 0 | 0 | No value | 0 | 0 | 1 |
| 451 | N | 0 | 0 | No value | 0 | 0 | 1 |
| 452 | G | 0 | 0 | No value | 0 | 0 | 1 |
| 453 | Y | 0 | 0 | No value | 0 | 0 | 1 |
| 454 | L | 0 | 0 | No value | 0 | 0 | 1 |
| 455 | L | 5.033 | 0 | 0 | 3.127 | 1.601 | 0.206 |
| 456 | L | 3.702 | 0 | 0 | 1.98 | 1.704 | 0.192 |
| 457 | D | 0 | 0 | No value | 0 | 0 | 1 |
| 458 | F | 0 | 0 | No value | 0 | 0 | 1 |
| 459 | F | 0 | 0 | No value | 0 | 0 | 1 |
| 460 | . | 0 | 11.774 | Infinity | 2.531 | 2.954 | 0.086 |
| 461 | N | 0 | 0 | No value | 0 | 0 | 1 |
| 462 | E | 0 | 0 | No value | 0 | 0 | 1 |
| 463 | V | 3.97 | 0 | 0 | 2.536 | 1.707 | 0.191 |
| 464 | . | 4.204 | 5.359 | 1.275 | 4.754 | 0.053 | 0.818 |
| 465 | G | 0 | 0 | No value | 0 | 0 | 1 |
| 466 | V | 0 | 0 | No value | 0 | 0 | 1 |
| 467 | A | 2.088 | 0 | 0 | 1.157 | 1.116 | 0.291 |
| 468 | F | 0 | 0 | No value | 0 | 0 | 1 |

| | | | | | | | |
|---|---|---|---|---|---|---|---|
| 469 | T | 0 | 0 | No value | 0 | 0 | 1 |
| 470 | T | 3.168 | 0 | 0 | 1.351 | 1.595 | 0.207 |
| 471 | V | 1.908 | 0 | 0 | 1.241 | 0.832 | 0.362 |
| 472 | V | 0 | 3.933 | Infinity | 1.247 | 2.296 | 0.13 |
| 473 | C | 0 | 0 | No value | 0 | 0 | 1 |
| 474 | G | 0 | 0 | No value | 0 | 0 | 1 |
| 475 | F | 0 | 0 | No value | 0 | 0 | 1 |
| 476 | T | 0 | 0 | No value | 0 | 0 | 1 |
| 477 | G | 2.909 | 0 | 0 | 1.503 | 1.318 | 0.251 |
| 478 | A | 1.924 | 0 | 0 | 1.105 | 1.051 | 0.305 |
| 479 | Y | 7.755 | 0 | 0 | 2.387 | 2.118 | 0.146 |
| 480 | . | 3.204 | 7.665 | 2.392 | 5.456 | 0.467 | 0.494 |
| 481 | A | 1.95 | 2.887 | 1.481 | 2.343 | 0.069 | 0.792 |
| 482 | F | 0 | 0 | No value | 0 | 0 | 1 |
| 483 | I | 0 | 0 | No value | 0 | 0 | 1 |
| 484 | I | 0 | 0 | No value | 0 | 0 | 1 |
| 485 | Y | 0 | 0 | No value | 0 | 0 | 1 |
| 486 | L | 0 | 0 | No value | 0 | 0 | 1 |
| 487 | I | 4.444 | 0 | 0 | 1.682 | 1.772 | 0.183 |
| 488 | S | 0 | 0 | No value | 0 | 0 | 1 |
| 489 | R | 0 | 0 | No value | 0 | 0 | 1 |
| 490 | G | 0 | 0 | No value | 0 | 0 | 1 |
| 491 | F | 11.922 | 3.8 | 0.319 | 6.783 | 0.833 | 0.361 |
| 492 | T | 3.85 | 2.567 | 0.667 | 3.282 | 0.107 | 0.744 |
| 493 | C | 0 | 0 | No value | 0 | 0 | 1 |
| 494 | F | 0 | 0 | No value | 0 | 0 | 1 |
| 495 | S | 2.03 | 3.202 | 1.577 | 2.5 | 0.1 | 0.752 |
| 496 | . | 0 | 7.81 | Infinity | 5.699 | 1.252 | 0.263 |
| 497 | C | 2.601 | 0 | 0 | 1.409 | 1.167 | 0.28 |
| 498 | C | 0 | 0 | No value | 0 | 0 | 1 |
| 499 | . | 3.96 | 5.692 | 1.437 | 4.719 | 0.122 | 0.727 |
| 500 | S | 6.648 | 0 | 0 | 3.803 | 3.051 | 0.081 |
| 501 | K | 0 | 0 | No value | 0 | 0 | 1 |
| 502 | Q | 0 | 0 | No value | 0 | 0 | 1 |
| 503 | I | 0 | 0 | No value | 0 | 0 | 1 |
| 504 | E | 0 | 0 | No value | 0 | 0 | 1 |
| 505 | V | 0 | 0 | No value | 0 | 0 | 1 |
| 506 | E | 0 | 0 | No value | 0 | 0 | 1 |

**Supplementary table S4.** Primers used for the cloning of *P. trichocarpa NRAMP3.*

| Primer names | Primer sequences |
|---|---|
| PtNramp3aLgtw | 5'-TACAAAAAAGCAGGCTTCATGCCTTCACCAGAAGAAGAC-3' |
| PtNramp3aRgtwnostp | 5'-CAAGAAAGCTGGGTCGGTGGGACAGTACCAAGTGG-3' |
| PtNramp3aRgtwstp | 5'-CAAGAAAGCTGGGTCCTGTTTAGGTGGGACAGTACCA-3' |
| PtNramp3bLlonggtw | 5'-GGGGACAAGTTTGTACAAAAAAGCAGGCTTCATGCCTGTAGAAGAAAAC-3' |
| PtNramp3bRlonggtwnostp | 5'-GGGGACCACTTTGTACAAGAAAGCTGGGTCTGTAAAGCCCCTAGAATG-3' |

| | |
|---|---|
| PtNramp3bRlonggtwstp | 5'-GGGGACCACTTTGTACAAGAAAGCTGGGTCTTATGTAAAGCCCCTAG-3' |
| U5 | 5'-GGGGACAAGTTTGTACAAAAAAGCAGGCTTC-3' |
| U3 | 5'-GGGGACCACTTTGTACAAGAAAGCTGGGTC-3' |

**Supplementary table S5.** Composition of media used for poplar transformation and *in vitro* propagation.

| Composition | MS30 | M1 | M2 | M3 | MS1/2 |
|---|---|---|---|---|---|
| Macro-elements (10X)[1] | 100 mL/L | 100 mL/L | 100 mL/L | 100 mL/L | 50 mL/L |
| Micro-elements (1000X)[2] | 1 mL/L | 1 mL/L | 1 mL/L | 1 mL/L | 1 mL/L |
| Ethylenediaminetetra-acetic acid ferric monosodium salt(Fe) | 40 mg/L | 40 mg/L | 40 mg/L | 40 mg/L | 40 mg/L |
| Myo-inositol | 100 mg/L | 100 mg/L | 100 mg/L | 100 mg/L | 100 mg/L |
| M.E.S | | 250 mg/L | 250 mg/L | 250 mg/L | |
| Vitamins (100X)[3] | 10 mL/L | 10 mL/L | 10 mL/L | 10 mL/L | 10 mL/L |
| L-Glutamine | 200 mg/L | 200 mg/L | 200 mg/L | 200 mg/L | 200 mg/L |
| Sucrose | 30 g/L | 30 g/L | 30 g/L | 30 g/L | 20 g/L |
| pH | 5.9 – 6 | 5.8 | 5.8 | 5.8 | 5.9 - 6 |
| Agar | | 7 g/L | 7 g/L | 7 g/L | 7 g/L |
| $N_6$-(2-Isopentenyl) adenine (2ip) | | 5 µM | 5 µM | | |
| α-Naphthaleneacetic Acid (NAA) | | 10 µM | 10 µM | | |
| Ticarpen | | | 500 mg/L | 500 mg/L | |
| Cefotaxime | | | 250 mg/L | 250 mg/L | |
| Thidiazuron (TDZ) | | | | 0.1 µM | |

[1]**Macro-elements (10X):** 16.5 g $NH_4NO_3$, 19 g $KNO_3$, 4.4 g $CaCl_2.2H_2O$, 3.7 g $MgSO_4.7H_2O$, 1.7 g $KH_2PO_4$ dissolved in 1000 ml $H_2O$.

[2]**Micro-elements (1000X):** 620 mg $H_3BO_3$, 1690 mg $MnSO_4$.H2O, 1060 mg $ZnSO_4.7H_2O$, 83 mg KI, 25 mg $Na_2MoO_4.2H_2O$, 2.5 mg $CuSO_4.5H_2O$, 2.5 mg $CoCl_2.6H_2O$ dissolved in 100 ml $H_2O$.

[3]**Vitamins (100X):** 50 mg Nicotinic acid, 50 mg Pyridoxine hydrochloride, 50 mg Thiamine hydrochloride, 50 mg Calcium pantothenate, 50 mg L-cysteine chlorohydrate, 5 ml of Biotine (50 mg/ 50 ml NaOH) dissolved in 500 ml $H_2O$.

**Supplementary table S6.** Primers used for the RT-qPCR.

| Genes | Forward primers | Reverse primers |
|---|---|---|
| *PotriNRAMP3.1* | TGACATTTGCGTGGATGTTT | ATTGTGGTCGATCTCCCTTG |
| *PotriNRAMP3.2* | CGGCCCCATTCTTAAGATGG | TGTAAAGCCCCTAGAAATGAGAT |
| *PtEF1* | CCACACCTGTCACATTGCTG | ACCAGCATCACCGTTCTTCAG |
| *PtPP2A* | TTCCTGATGTGCGACTGAAC | CTCCAATGCCTATCCTCTGC |
| *PtUBQ* | CCATATCCAAGGTATTGCTCTCC | CGTCTCATACTTGTTCCTGTGG |
| *AtACTIN* | GGTAACATTGTGCTCAGTGGTGG | GGTAACATTGTGCTCAGTGGTGG |

### Supplementary Data

**Supplementary data S1.** NRAMP3 coding sequences identified in *Populus* and *Salix* genomes

>P. trichocarpa_NRAMP3.1_CDS
ATGCCTTCACCAGAAGAAGACCCACAACCTTTATTAAAAGACCAAGAAGAAACAGCTTATGATT
CTGACGGGAAAGTCCTTTCATTTGGGATTGATTATGACACAGAAAGCGGTGGCTCAACGGTGGT
GCCATCATTTTCATGGAGAAAATTATGGTTGTTCACTGGTCCTGGGTTTTTAATGTGCATTGCTTT
TTTGGACCCTGGCAATTTGGAAGGGGATCTTCAGGCTGGTGCAATTGCAGGGTATTCTTTGCTTT
GGCTTCTCTTATGGGCTACTGCTATGGGTTTGTTGGTGCAGTTGCTGTCAGCAAGGCTTGGAGTG
GCTACAGGAAGGCATTTGGCTGAGCTATGTAGAGAAGAGTATCCAACTTGGGCTCGAATGATTT
TGTGGATTATGGCTGAGTTGGCTTTGATTGGTGCTGATATACAAGAAGTTATTGGGAGTGCTATT
GCTATTCAGATTTTGAGTAATGGGGTTTTGCCTTTGTGGGCTGGTGTTATTATTACTGCTTCCGAT
TGCTTTATCTTCCTATTTCTTGAGAACTACGGTGTGAGGAAATTGGAGGCTGCTTTTGGGATTCTC
ATTGGAATAATGGCAGTGACATTTGCGTGGATGTTTGCTGATGCAAAACCCAGTGCCCCCGAACT
TTTTCTGGGCATCTTAATTCCAAAACTTAGCTCCAAAACAATAAAACAGGCTGTTGGAGTTGTGG
GTTGCATTATCATGCCTCACAATGTGTTCTTGCATTCTGCTCTTGTACAGTCAAGGGAGATCGAC
CACAATAAGAAAGGCCAGGTTCAAGAAGCTCTCAGATACTACTCCATAGAGTCAACTGCTGCCC
TTGCAATATCATTCATGATCAATTTGTTTGTGACGACCATTTTTGCTAAAGGTTTCCACGGGACA
GAACTGGCCAATAGTATTGGCCTTGTAAATGCAGGGCAATATCTTCAAGATAAATACGGGGGTG
GATTTTTCCCAATTTTATACATCTGGGGTATTGGGTTATTAGCAGCTGGCCAAAGTAGCACCATT
ACTGGCACTTATGCAGGGCAGTTTATCATGGGAGGTTTCCTGAACTTGGGGTTAAAGAAATGGCT
GAGGGCATTGATTACTCGAAGCTGTGCTATCATCCCAACTATAATTGTTGCACTTGTTTTTGATAC
TTCTGAAGACTCACTAGATGTTCTGAATGAATGGCTAAATATGCTTCAGTCTATTCAGATTCCTTT
TGCACTCATCCCTCTTCTTTGCTTGGTCTCTAAGGAGCAAATCATGGGCACTTTCACAGTTGGCCC
CATTCTTAAGATGGTTTCTTGGCTTGTAGCTGCCTTGGTGATGCTAATCAATGGTTACCTTTTGCT
TGACTTTTTCTCCAATGAAGTAACTGGAGTAGTGTTTACCACTGTGGTATGCGCTTTTACAGGAG
CATATGTTACGTTTATAATTTATCTCATTTCTAGGGAAGTTACCATTTCCACTTGGTACTGTCCCA
CC

>P. trichocarpa_NRAMP3.2_CDS
ATGCCTGTAGAAGAAAACTACCAACCTTTATTGCAAGAAGAAGAAGAAAGAGCTTATGATTCTG
ATGAGAAAGTGCTCATAATTGGGGTTGATTCTGACACGGAAAGCGGTGGCTCAACGGTGTTGCC
ACCGTTTTCATGGAAAAAGTTATGGTTGTTTACTGGTCCTGGGTTTTTAATGTCCATTGCGTTTTT
GGATCCTGGGAATTTGGAAGGGGATCTTCAGGCTGGTGCAATCGCAGGCTACTCTTTGCTTTGGC
TTCTTTTATGGGCTACTGCTATGGGGTTGTTGGTGCAGTTGCTTTCAGCGAGGCTTGGAGTGGCT
ACAGGGAGGCATTTAGCTGAGCTGTGTAGAGAAGAGTATCCAACCTGGGCTTCAATGGTTTTGT
GGATTATGGCTGAGTTGGCTTTGATTGGTGCTGATATACAAGAGGTTATTGGAAGTGCTATTGCT
ATTAAGATCTTGAGTAATGGGTTTGTGCCTTTGTGGGCTGGTGTTACTATTACTGCTTGTGATTGC
TTCATCTTCCTATTTCTAGAGAACTACGGTGTGAGAAAATTGGAGGCTGTATTTGCGGTCCTTATT
GGAATAATGGCAGTTACATTTGGATGGATGTTTGCAGATGCAAAACCCAGTGCCTCCGAACTTTT
TCTGGGTATCTTAATTCCAAAACTTAGCTCCAGAACAATACAACAGGCTGTTGGAGTTGTGGGTT
GCATTATCATGCCTCACAATGTGTTCTTGCATTCTGCTCTTGTACAGTCAAGGGAGATCGACCAC
AATAAGAAAGACCGGGTTCAAGAAGCTCTCAGATACTACTCCATAGAGTCAACCACTGCCCTTG
TAATATCGTTCGTAATCAATTTGTTTGTGACGACTGTTTTTGCTAAAGGTTTCTACGGGACAGAA
CTAGCCAATAGTATTGGCCTTGTAAATGCAGGGCAATATCTTCAAGACAAATACGGGGGTGGAT
TTTTCCCAATTTTATACATCTGGGGTATTGGATTATTAGCAGCTGGCCAAAGCAGCACCATTACT
GGCACTTATGCAGGACAGTTTATCATGGGAGGTTTCCTGAACTTGAGGTTAAAGAAATGGCTTA
GGGCATTGATCACTCGAAGCTGTGCTATCATCCCAACTATGATTGTTGCACTTGTTTTTGATACCT
CCGAAGACTCACTAGATGTTCTGAATGAATGGCTAAATGTGCTGCAGTCAATACAGATTCCTTTT

GCACTCATCCCTCTTCTCTGCTTAGTATCCAAGGAGCAAATCATGGGCACTTTCAAAATCGGCCC
CATTCTTAAGATGGTAGCTTGGCTTGTGGCTGCCCTGGTGATGGTAATCAATGGTTACCTTTTGCT
CGACTTTTTCTTCAATGAAGTGACCGGAGTAGCGTTTACCACTGTAGTATGCGGTTTTACAGGTG
CATATGTTGCGTTTATAATTTATCTCATTTCTAGGGGCTTTACATGTTTCTCCCGGTGCTGTCCAT
CTAAACAGATAGAAGTAGAG

>P. alba_NRAMP3.1_CDS
ATGCCTTTACCAGAAGAAGACCCGCAACCTTTATTAAAAGACCAAGAAGAAACAGCTTATGATT
CTGACGGGAAAGTCCTTTCATTTGGCATTGATTATGACACGGAAAGCGGTGGCTCAACGGCGGT
GTCATCATTTTCATGGAGAAAATTATGGTTGTTCACTGGTCCTGGGTTTTTAATGTGCATTGCTTT
TTTGGACCCTGGCAATTTGGAAGGGGATCTTCAGGCTGGTGCAATTGCAGGGTATTCTTTGCTTT
GGCTTCTCTTATGGGCTACTGCTATGGGTTTGTTGGTGCAGTTGCTGTCAGCAAGGCTTGGAGTG
GCTACAGGAAGGCATTTGGCTGAGCTATGTAGAGAAGAGTATCCAACTTGGGCTCGAATGATTT
TGTGGATTATGGCTGAGTTGGCTTTGATTGGTGCTGATATACAAGAAGTTATTGGGAGTGCTATT
GCTATTCAGATTTTGAGTAATGGGGTTTTGCCTTTGTGGGCTGGTGTTATTATTACTGCTTTCGAT
TGCTTTATCTTCCTATTTCTTGAGAACTACGGTGTGAGGAAATTGGAGGCTGCTTTTGGGATCCTC
ATTGGAATAATGGCAGTGACATTTGCGTGGATGTTTGCTGATGCAAAACCCAGTGCCCCTGAACT
TTTTCTAGGCATCTTAATTCCAAAACTTAGCTCCAAAACAATAAAACAGGCTGTTGGAGTTGTGG
GTTGCATTATCATGCCTCACAATGTGTTCTTGCATTCTGCTCTTGTACAGTCAAGGGAGATTGACC
ACAATAAGAAAGGCCAGGTTCAAGAAGCTCTCAGATACTACTCCATAGAGTCAACTGCTGCCCT
TGCAATATCATTCATGATCAATTTGTTTGTGACGACCATTTTTGCTAAAGGTTTCCACGGGACAG
AACTGGCCAATAGTATTGGCCTTGTAAATGCAGGGCAATATCTTCAAGATAAATACGGGGGTGG
ATTTTTCCCAATTTTATACATCTGGGGTATTGGGTTATTAGCAGCTGGCCAAAGTAGCACCATTA
CTGGCACTTATGCAGGGCAGTTTATCATGGGAGGTTTCCTGAACTTGGGATTAAAGAAATGGCTG
AGGGCATTGATTACTCGAAGCTGTGCTATCATCCCAACTATAATTGTTGCACTTGTTTTTGATACT
TCTGAAGACTCACTAGATGTTCTGAATGAATGGCTAAATATGCTTCAGTCTATTCAGATTCCTTTT
GCACTCATCCCTCTTCTTTGCTTGGTCTCCAAGGAGCAAATCATGGGCACTTTCACAGTTGGCCC
CATTCTTAAGATGTTTTCTTGGCTTGTAGCTGCCTTGGTGATGCTAATCAATGGTTACCTTTTGCT
TGACTTTTTCTCCAATGAAGTAACTGGAGTAGTGTTTACCACTGTGGTATGCTCTTTTACAGGAG
CATATGTTACGTTTATAATTTATCTCATTTCTAGGGAAGTATCCATTTCCACTTGGTACTGTCCCA
CC

>P. alba_NRAMP3.2_CDS
ATGCCTGTAGAAGAAAACCAACAACCTTTATTGCAAGAAGAAGAAGAAAGAGCTTATGATTCTG
ATGAGAAAGTGCTCATAATTGGGGTTGATTCTGACACGGAAAGCGGTGGCTCAACGGTGTTGCC
ACCGTTTTCATGGAAAAAGTTATGGTTGTTTACTGGTCCTGGGTTTTTAATGTCCATTGCGTTTTT
GGATCCTGGGAATTTGGAAGGGGATCTTCAGGCTGGTGCTATCGCAGGCTACTCTTTGCTTTGGC
TTCTTTTATGGGCTACTGCTATGGGTTTGTTGGTGCAGTTGCTTTCAGCGAGGCTTGGAGTGGCTA
CAGGGAGGCATTTAGCTGAGCTGTGTAGAGAAGAGTATCCAACCTGGGCGTCAATGGTTTTGTG
GATTATGGCTGAGTTGGCTTTGATTGGTGCTGATATACAAGAGGTTATTGGAAGTGCTATTGCTC
TTAAGATCTTGAGTAATGGGTTTTTGCCTCTGTGGGCTGGTGTTACTATTACTGCTTGTGATTGCT
TCATCTTCCTATTTCTAGAGAACTACGGTGTGAGAAAATTGGAGGCTGTATTTGCGGTCCTTATT
GGAATAATGGCAGTTACATTTGGATGGATGTTTGCAGATGCAAAACCCAGTGCCTCCGAACTTTT
TCTGGGTATCTTAATTCCCAAACTTAGCTCCAGAACAATACAACAGGCTGTTGGAGTTGTGGGTT
GCATTATCATGCCTCACAATGTGTTCTTGCATTCTGCTCTTGTGCAGTCAAGGGAGATCGACCAC
AATAAGAAAGGCCGGGTTCAAGAAGCTCTCAGATACTACTCCATAGAGTCAACCACTGCCCTTG
TAATATCATTCGTAATCAATTTGTTTGTGACGACTGTTTTTGCTAAAGGTTTCTATGGGACAGAAC
TGGCCAATAGTATTGGCCTTGTAAATGCAGGGCAATATCTTCAAGACAAATACGGGGGTGGATT
TTTCCCAATTTTATACATCTGGGGTATTGGATTATTAGCAGCTGGCCAAAGCAGCACCATTACTG
GCACTTATGCAGGACAGTTTATCATGGGAGGTTTCCTGAACTTGAGGTTAAAGAAATGGTTAAG

GGCATTGATCACTCGAAGCTGTGCTATCATCCCAACTATGATTGTTGCACTTGTTTTTGATACCTC
TGAAGACTCACTAGATGTTCTGAATGAATGGCTAAATGTGCTGCAGTCAATACAGATTCCTTTTG
CACTCATCCCTCTTCTCTGCTTGGTATCCAAGGAGCAAATCATGGGCACTTTCAAAATCGGTCCC
ATTCTTAAGGTATCTTGGCTTGTGGCTGCCCTGGTGATAGTAATCAATGGTTACCTTTTGCTCGAC
TTTTTCGTCAATGAAGTGGCCGGAGTAGCGTTTACCACTGTACTATGCGGTTTTACAGGTGCATA
TGTAGCGTTTATAATTTATCTCATTTCTAGGGGCTTTACATGTTTCTTCTGGTGCTGTCAATCTAA
ACAGATAGAAGTAGAG

>P. cathayana_NRAMP3.1_CDS
ATGCCTTTACCAGAAGAAGACCCACAACCTTTATTAAAAGACCAAGAAGAAACAGCTTATGATT
CTGACGGGAAAGTCCTTTCATTTGGGATTGATTATGACACCGAAAGCGGTGGCTCAACGGTGGT
GCCATCATTTTCATGGAGAAAATTATGGTTGTTCACTGGTCCTGGGTTTTTAATGTGCATTGCTTT
TTTGGACCCTGGCAATTTGGAAGGGGATCTTCAGGCTGGTGCAATTGCAGGGTATTCTTTGCTTT
GGCTTCTCTTATGGGCTACTGCTATGGGTTTGTTGGTGCAGTTGCTGTCAGCAAGGCTTGGAGTG
GCTACAGGAAGGCATTTGGCTGAGCTATGTAGAGAAGAGTATCCAACTTGGGCTCGAATGATTT
TGTGGATTATGGCTGAGTTGGCTTTGATTGGTGCTGATATACAAGAAGTTATTGGGAGTGCTATT
GCTATTCAGATTTTGAGTAATGGGGTTTTGCCTTTGTGGGCTGGTGTTATTATTACTGCTTCCGAT
TGCTTTATCTTCCTATTTCTTGAGAACTACGGTGTGAGGAAATTGGAGGCTGCTTTTGGGATTCTC
ATTGGAATAATGGCAGTGACATTTGCGTGGATGTTTGCTGATGCAAAACCCAGTGCCCCCGAACT
TTTTCTGGGCATCTTAATTCCAAAACTTAGCTCCAAAACAATAAAACAGGCTGTTGGAGTTGTGG
GTTGCATTATCATGCCTCACAATGTGTTCTTGCATTCTGCTCTTGTACAGTCAAGGGAGATCGAC
CACAATAAGAAAGGCCAGGTTCAAGAAGCTCTCAGATACTACTCCATAGAGTCAACTGCTGCCC
TTGCAATATCATTCATGATCAATTTGTTTGTGACGACCGTTTTTGCTAAAGGTTTCCACGGGACA
GAACTGGCCAATAGTATTGGCCTTGTAAATGCAGGGCAATATCTTCAAGATAAATACGGGGGTG
GATTTTTCCCAATTTTATACATCTGGGGTATTGGGTTATTAGCAGCTGGCCAAAGTAGCACCATT
ACTGGCACTTATGCAGGGCAGTTTATCATGGGAGGTTTCCTGAACTTGGGGTTAAAGAAATGGCT
GAGGGCATTGATTACTCGAAGCTGTGCTATCATCCCAACTATAATTGTTGCACTTGTTTTTGATAC
TTCTGAAGACTCACTAGATGTTCTGAATGAATGGCTAAATATGCTTCAGTCTATTCAGATTCCTTT
TGCACTCATCCCTCTTCTTTGCTTGGTCTCTAAGGAGCAAATCATGGGCACTTTCACAGTTGGCCC
CATTCTTCAGATGGTTTCTTGGCTTGTAGCTGCCTTGGTGATGCTAATCAATGGTTACCTTTTGCT
TGACTTTTTCTCCAATGAAGTGACTGGAGTAGTGTTTACCACTGTGGTATGCGCTTTTACAGGAG
CATATGTTACGTTTATAATTTATCTCATTTCTAGGGAAGTTACCATTTCCACTTGGTACTGTCCTA
CC

>P. cathayana_NRAMP3.2_CDS
ATGCCTGTAGAAGAAAACCACCAACCTTTATTGCAAGAAGAAGAAGAAAGAGCTTATGATTCTG
ATGAGAAAGTGCTCATAATTGGGGTCGATTCTGACACGGAAAGCGGTGGCTCAACGGTGTTGCC
ACCGTTTTCATGGAAAAAGTTATGGTTGTTTACTGGTCCTGGGTTTTTAATGTCCATTGCGTTTTT
GGATCCTGGGAATTTGGAAGGGGATCTTCAGGCTGGTGCAATCGCAGGCTACTCTTTGCTTTGGC
TTCTTTTATGGGCTACTGCTATGGGGTTGTTGGTGCAGTTGCTTTCAGCGAGGCTTGGAGTGGCT
ACAGGGAGACATTTAGCTGAGCTGTGTAGAGAAGAGTATCCAACCTGGGCTTCAATGGTTTTGT
GGATTATGGCTGAGTTGGCTTTGATTGGTGCTGATATACAAGAGGTTATTGGAAGTGCTATTGCT
ATTAAGATCTTGAGTAATGGGTTTGTGCCTTTGTGGGCTGGTGTTACTATTACTGCTTGTGATTGC
TTCATCTTCCTATTTCTAGAGAACTACGGTGTGAGAAAATTGGAGGCTGTATTTGCGGTCCTTATT
GGAATAATGGCAGTTACATTTGGATGGATGTTTGCAGATGCAAAACCCAGTGCCTCCGAACTTTT
TCTGGGTATCTTAATTCCAAAACTTAGCTCCAGAACAATACAACAGGCTGTTGGAGTTGTGGGTT
GCATTATCATGCCTCACAATGTGTTCTTGCATTCTGCTCTTGTACAGTCAAGGGAGATCGACCAC
AATAAGAAAGACCGGGTTCAAGAAGCTCTCAGATACTACTCCATAGAGTCAACCACTGCCCTTG
TAATATCGTTCGTAATCAATTTGTTTGTGACGACTGTTTTTGCTAAAGGTTTCTACGGGACAGAA
CTGGCCAATAGTATTGGCCTTGTAAATGCAGGGCAATATCTTCAAGACAAATACGGGGGTGGAT

TTTTCCCAATTTTATACATCTGGGGTATTGGATTATTAGCAGCTGGCCAAAGCAGCACCATTACT
GGCACTTATGCAGGACAGTTTATCATGGGAGGTTTCCTGAACTTGAGGTTAAAGAAATGGCTAA
GGGCATTGATCACTCGAAGCTGTGCTATCATCCCAACTATGATTGTTGCACTTGTTTTTGATACCT
CCGAAGACTCACTAGATGTTCTGAATGAATGGCTAAATGTGCTGCAGTCAATACAGATTCCTTTT
GCACTCATCCCTCTTCTCTGCTTAGTATCCAAGGAGCAAATCATGGGCACTTTCAAAATCGGCCC
CATTCTTAAGATGGTAGCTTGGCTTGTGGCTGCCCTGGTGATGGTAATCAATGGTTACCTTTTGCT
CGACTTTTTCTTCAATGAAGTGACCGGAGTAGCATTTACCACAGTAGTATGCGGTTTTACAGGTG
CATATGCTGCGTTTATAATTTATCTCATTTCTAGGGGCTTTACATGTTTCTCCCGGTGCTGTCCAT
CTAAACAGATAGAAGTAGAG

>P. simonii_NRAMP3.1_CDS
ATGCCTTTACCAGAAGAAGACCCACGACCTTTATTAAAAGACCAAGAAGAAACAGCTTATGATT
CTGACGGGAAAGTCCTTTCATTTGGGATTGATTATGACACCGAAAGCGGTGGCTCAACGGTGGT
GCCATCATTTTCATGGAGAAAATTATGGTTGTTCACTGGTCCTGGGTTTTTAATGTGCATTGCTTT
TTTGGACCCTGGCAATTTGGAAGGGGATCTTCAGGCTGGTGCAATTGCAGGGTATTCTTTGCTTT
GGCTTCTCTTATGGGCTACTGCTATGGGTTTGTTGGTGCAGTTGCTGTCAGCAAGGCTTGGAGTG
GCTACAGGAAGGCATTTGGCTGAGCTATGTAGAGAAGAGTATCCAACTTGGGCTCGAATGATTT
TGTGGATTATGGCTGAGTTGGCTTTGATTGGTGCTGATATACAAGAAGTTATTGGGAGTGCTATT
GCTATTCAGATTTTGAGTAATGGGGTTTTGCCTTTGTGGGCTGGTGTTATTATTACTGCTTCCGAT
TGCTTTATCTTCCTATTTCTTGAGAACTACGGTGTGAGGAAATTGGAGGCTGCTTTTGGGATTCTC
ATTGGAATAATGGCAGTGACATTTGCGTGGATGTTTGCTGATGCAAAACCCAGTGCCCCCGAACT
TTTTCTGGGCATCTTAATTCCAAAACTTAGCTCCAAAACAATAAAACAGGCTGTTGGAGTTGTGG
GTTGCATTATCATGCCTCACAATGTGTTCTTGCATTCTGCTCTTGTACAGTCAAGGGAGATTGACC
ACAATAAGAAAGGCCAGGTTCAAGAAGCTCTCAGATACTACTCCATAGAGTCAACTGCTGCCCT
TGCAATATCATTCATGATCAATTTGTTTGTGACGACTGTTTTTGCTAAAGGTTTCCACGGGACAG
AACTGGCCAATAGTATTGGCCTTGTAAATGCAGGGCAATATCTTCAAGATAAATACGGGGGTGG
ATTTTTCCCAATTTTATACATCTGGGGTATTGGGTTATTAGCAGCTGGCCAAAGTAGCACCATTA
CTGGCACTTATGCAGGGCAGTTTATCATGGGAGGTTTCCTGAACTTGGGGTTAAAGAAATGGCTG
AGGGCATTGATTACTCGAAGCTGTGCTATCATCCCAACTACAATTGTTGCACTTGTTTTTGATACT
TCTGAAGACTCACTAGATGTTCTGAATGAATGGCTAAATATGCTTCAGTCTATTCAGATTCCTTTT
GCACTCATCCCTCTTCTTTGCTTGGTCTCTAAGGAGCAAATCATGGGCACTTTCACAGTTGGCCCC
ATTCTTCAGATGGTTTCTTGGCTTGTAGCTGCCTTGGTGATGCTAATCAATGGTTACCTTTTGCTT
GACTTTTTCTCCAATGAAGTAACTGGAGTAGTGTTTACCACTGTGGTATGCGCTTTTACAGGAGC
ATATGTTACGTTTATAATTTATCTCATTTCTAGGGAAGTTACCATTTCCACTTGGTACTGTCCCAC
C

>P. simonii_NRAMP3.2_CDS
ATGCCTGTAGAAGAAAACCACCAACCTTTATTGCAAGAAGAAGAAGAAAGAGCTTATGATTCTG
ATGAGAAAGTGCTCATAATTGGGGTTGATTCTGACACGGAAAGCGGTGGCTCAACGGTGTTGCC
ACCGTTTTCATGGAAAAAGTTATGGTTGTTTACTGGTCCTGGGTTTTTAATGTCCATTGCGTTTTT
GGATCCTGGGAATTTGGAAGGGGATCTTCAGGCTGGTGCAATCGCAGGCTACTCTTTGCTTTGGC
TTCTTTTATGGGCTACTGCTATGGGGTTGTTGGTGCAGTTGCTTTCAGCGAGGCTTGGAGTGGCT
ACAGGGAGGCATTTAGCTGAGCTGTGTAGAGAAGAGTATCCAACCTGGGCTTCAATGGTTTTGT
GGATTATGACTGAGTTGGCTTTGATTGGTGCTGATATACAAGAGGTTATTGGAAGTGCTATTGCT
ATTAAGATCTTGAGTAATGGGTTTGTGCCTTTGTGGGCTGGTGTTACTATTACTGCTTGTGATTTC
ATCTTCCTATTTCTAGAGAACTACGGTGTGAGAAAATTGGAGGCTGTATTTGCGGTCCTTATTGG
AATAATGGCAGTTACATTTGGATGGATGTTTGCAGATGCAAAACCCAGTGCTTCCGAACTTTTTC
TGGGTATCTTAATTCCAAAACTTAGCTCCAGAACAATACAACAGGCTGTTGGAGTTGTGGGTTGC
ATTATCATGCCTCACAATGTGTTCTTGCATTCTGCTCTTGTACAGTCAAGGGAGATCGACCACAA
TAAGAAAGACCGGGTTCAAGAAGCTCTCAGATACTACTCCATAGAGTCAACCACTGCCCTTGTA

ATATCGTTCGTAATCAATTTGTTTGTGACGACTGTTTTTGCTAAAGGTTTCTACGGGACAGAACT
GGCCAATAGTATTGGCCTTGTAAATGCAGGGCAATATCTTCAAGACAAATACGGGGGTGGATTT
TTCCCAATTTTATACATCTGGGGTATTGGATTATTAGCAGCTGGCCAAAGCAGCACCATTACTGG
CACTTATGCAGGACAGTTTATCATGGGAGGTTTCCTGAACTTGAGGTTAAAGAAATGGCTAAGG
GCATTGATCACTCGAAGCTGTGCTATCATCCCAACTATGATTGTTGCACTTGTTTTTGATACCTCC
GAAGACTCACTAGATGTTCTGAATGAATGGCTAAATGTGCTGCAGTCAATACAGATTCCTTTTGC
ACTCATCCCTCTTCTCTGCTTAGTATCCAAGGAGCAAATCATGGGCACTTTCAAAATCGGCCCCA
TTCTTAAGATGGTATCTTGGCTTGTGGCTGCCCTGGTGATGGTAATCAATGGTTACCTTTTACTCG
ACTTTTTCTTCAATGAAGTGACCGGAGTAGCGTTTACCACTGTAGTATGCGGTTTTACAGGTGCA
TATGTTGCGTTTATAATTTATCTCATTTCTAGGGGCTTTACATGTTTCTCCCGGTGCTGTCCATCTA
AACAGATAGAAGTAGAG

>P. lasiocarpa _NRAMP3.1_CDS
ATGCCTTTACCAGAAGAAGACCCAAAACCTTTATTAAAAGACCAAGAAGAAACAGCTTATGATT
CTGACGGGAAAGTCCTTTCATTTGGGATTGATTATGGCACCGAAAGCGGTGGCTCAACGGTGGT
GCCATCATTTTCATGGAGAAAATTATGGTTGTTCACTGGTCCTGGGTTTTTAATGTGCATTGCTTT
TTTGGACCCTGGCAATTTGGAAGGGGATCTTCAGGCTGGTGCAATTGCTGGGTATTCTTTGCTTT
GGCTTCTCTTATGGGCTACTGCTATGGGTTTGTTGGTGCAGTTGCTGTCAGCAAGGCTTGGAGTG
GCTACAGGAAGGCATTTGGCTGAGCTATGTAGAGAAGAGTATCCAACTTGGGCTCGAATGATTT
TGTGGATTATGGCTGAGTTGGCTTTGATTGGTGCTGATATACAAGAAGTTATTGGGAGTGCTATT
GCTATTCAGATTTTGAGTAATGGGGTTTTGCCTTTGTGGGCTGGTGTTATTATTACTGCTTCCGAT
TGCTTTATCTTCCTATTTCTTGAGAACTACGGTGTGAGGAAATTGGAGGCTGCTTTTGGGATTCTC
ATTGGAATAATGGCAGTGACATTTGCGTGGATGTTTGCTGATGCAAAACCCAGTGCCCCCGAACT
TTTTCTGGGCATCTTAATTCCAAAACTTAGCTCCAAAACAATAAAACAGGCTGTTGGAGTTGTGG
GTTGCATTATCATGCCTCACAATGTGTTCTTGCATTCTGCTCTTGTACAGTCAAGGGAGATTGACC
ACAATAAGAAAGGCCAGGTTCAAGAAGCTCTCAGATACTACTCCATAGAGTCAACTGCTGCCCT
TGCAATATCATTCATGATCAATTTGTTTGTGACGACCGTTTTTGCTAAAGGTTTCCATGGGACAG
AACTGGCCAATAGTATTGGCCTTGTAAATGCAGGGCAATATCTTCAAGATAAATACGGAGGTGG
ATTTTTCCCAATTTTTTACATCTGGGGTATTGGGTTATTAGCAGCTGGCCAAAGTAGCACCATTAC
TGGCACTTATGCAGGGCAGTTTATCATGGGAGGTTTCCTGAACTTGGGGTTAAAGAAATGGCTG
AGGGCATTGATTACTCGAAGCTGTGCTATCATCCCAACTATAATTGTTGCACTTGTATTTGATACT
TCTGAAGACTCACTAGATGTTCTGAATGAATGGCTAAATATGCTTCAGTCTATTCAGATTCCTTTT
GCACTCATCCCTCTTCTTTGCTTGGTCTCCAAGGAGCAAATCATGGGCACTTTCACAGTTGGCCC
CATTCTTCAGATGGTTTCTTGGCTTGTAGCTGCCTTGGTGATGCTAATCAATGGTTACCTTTTGCT
TGACTTTTTCTCCAATGAAGTAACTGGAGTAGTGTTTACCACTGTGGTATGCGCTTTTACAGGAG
CATATGTTACGTTTATAATTTATCTCATTTCTAGGGAAGTTACCATTTCCACTTGGTACTGTCCCA
CC

>P. lasiocarpa _NRAMP3.2_CDS
ATGCCTGTAGAAGAAAACCACCAACCTTTATTGCAAGAAGAAGAAGAAAGAGCTTATGATTCTG
ATGAGAAAGTGCTCATAATTGGGGTTGATTCTGACACGGAAAGCGGTGGCTCAACGGTGTTGCC
ACCGTTTTCATGGAAAAAGTTATGGTTGTTTACTGGTCCTGGGTTTTTAATGTCCATTGCGTTTTT
GGATCCTGGGAATTTGGAAGGGGATCTTCAGGCTGGTGCAATCGCAGGCTACTCTTTGCTTTGGC
TTCTTTTATGGGCTACTGCTATGGGGTTGTTGGTGCAGTTGCTTTCAGCGAGGCTTGGAGTGGCT
ACAGGGAGGCATTTAGCTGAGCTGTGTAGAGAAGAGTATCCAACCTGGGCTTCAATGGTTTTGT
GGATTATGGCTGAGTTGGCTTTGATTGGTGCTGATATACAAGAGGTTATTGGAAGTGCTATTGCT
ATTAAGATCTTGAGTAATGGGTTTGTGCCTTTGTGGGCTGGTGTTACTATTACTGCTTGTGATTGG
TTCATCTTCCTATTTCTAGAGAACTACGGCGTGAGAAAATTGGAGGCTGTATTTGCGGTCCTTAT
TGGAATAATGGCAGTTACATTTGGATGGATGTTTGCAGATGCAAAACCCAGTGCCTCCGAACTTT
TTCTGGGTATCTTAATTCCAAAACTTAGCTCCAGAACAATACAACAGGCTGTTGGAGTTGTGGGT

TGCATTATCATGCCTCACAATGTGTTCTTGCATTCTGCTCTTGTACAGTCAAGGGAGATCGACCA
CAATAAGAAAGACCGGGTTCAAGAAGCTCTCAGATACTACTCCATAGAGTCAACCACTGCCCTT
GTAATATCATTCGTAATCAATTTGTTTGTGACGACTGTTTTTGCTAAAGGTTTCTACGGGACAGA
ACTGGCCAATAGTATTGGCCTTGTAAATGCAGGGCAATATCTTCAAGACAAATACGGGGGTGGA
TTTTTCCCAATTTTATACATCTGGGGTATTGGATTATTAGCAGCTGGCCAAAGCAGCACCATTAC
TGGCACTTATGCAGGACAGTTTATCATGGGAGGTTTCCTGAACTTGAGGTTAAAGAAATGGCTA
AGGGCATTGATCACTCGAAGCTGTGCTATCATCCCAACTATGATTGTTGCACTTGTTTTTGATACC
TCCGAAGACTCACTAGATGTTCTGAATGAATGGCTAAATGTGCTGCAGTCAATACAGATTCCTTT
TGCACTCATCCCTCTTCTCTGCTTAGTATCCAAGGAGCAAATCATGGGCACTTTCAAAATCGACC
CCATTCTTAAGATGGTATCTTGGCTTGTGGCTGCCCTGGTGATGGTAATCAATGGTTACCTTTTGC
TCGACTTTTTCTTCAATGAAGTGACCGGAGTAGCGTTTACCACTGTAGTATGCGGTTTTACAGGT
GCATATGTTGCGTTTATAATTTATCTCATTTCTAGGGGCTTTACATGTTTCTCCCGGTGCTGTCCA
TCTAAACAGATAGAAGTAGAG

>P. maximowiczii _NRAMP3.1_CDS
ATGCCTTTACCAGAAGAAGACCCACAACCTTTATTAAAAGACCAAGAAGAAACAGCTTATGATT
CTGACGGGAAAGTCCTTTCATTTGGGATTGATTATGACACCGAAAGCGGTGGCTCAACGGTGGT
GCCATCATTTTCATGGAGAAAATTATGGTTGTTCACTGGTCCTGGGTTTTTAATGTGCATTGCTTT
TTTGGACCCTGGCAATTTGGAAGGGGATCTTCAGGCTGGTGCAATTGCAGGGTATTCTTTGCTTT
GGCTTCTCTTATGGGCTACTGCTATGGGTTTGTTGGTGCAGTTGCTGTCAGCAAGGCTTGGAGTG
GCTACAGGAAGGCATTTGGCTGAGCTATGTAGAGAAGAGTATCCAACTTGGGCTCGAATGATTT
TGTGGATTATGGCTGAGTTGGCTTTGATTGGTGCTGATATACAAGAAGTTATTGGGAGTGCTATT
GCTATTCAGATTTTGAGTAATGGGGTTTTGCCTTTGTGGGCTGGTGTTATTATTACTGCTTCCGAT
TGCTTTATCTTCCTATTTCTTGAGAACTACGGTGTGAGGAAATTGGAGGCTGCTTTTGGGATTCTC
ATTGGAATAATGGCAGTGACATTTGCGTGGATGTTTGCTGATGCAAAACCCAGTGCCCCCGAACT
TTTTCTAGGCATCTTAATTCCAAAACTTAGCTCCAAAACAATAAAACAGGCTGTTGGAGTTGTGG
GTTGCATTATCATGCCTCACAATGTGTTCTTGCATTCTGCTCTTGTACAGTCAAGGGAGATTGACC
ACAATAAGAAAGGCCAAGTTCAAGAAGCTCTCAGATACTACTCCATAGAGTCAACTGCTGCCCT
TGCAATATCATTCATGATCAATTTGTTTGTGACGACTGTTTTTGCTAAAGGTTTCCACGGGACAG
AACTGGCCAATAGTATTGGCCTTGTAAATGCAGGGCAATATCTTCAAGATAAATACGGGGGTGG
ATTTTTCCCAATTTTATACATCTGGGGTATTGGGTTATTAGCAGCTGGCCAAAGTAGCACCATTA
CTGGCACTTATGCAGGGCAGTTTATCATGGGAGGTTTCCTGAACTTGGGGTTAAAGAAATGGCTG
AGGGCATTGATTACTCGAAGCTGTGCTATCATCCCAACTATAATTGTTGCACTTGTTTTTGATACT
TCTGAAGACTCACTAGATGTTCTGAATGAATGGCTAAATATGCTTCAGTCTATTCAGATTCCTTTT
GCACTCATCCCTCTTCTTTGCTTGGTCTCTAAGGAGCAAATCATGGGCACTTTCACAGTTGGCCCC
ATTCTTAAGATGGTTTCTTGGCTTGTAGCTGCCTTGGTGATGCTAATCAATGGTTACCTTTTGCTT
GACTTTTTCTCCAATGAAGTGACTGGAGTAGTGTTTACCACTGTGGTATGCGCTTTTACAGGAGC
ATATGTTACGTTTATAATTTATCTCATTTCTAGGGAAGTTACCATTTCCACTTGGTACTGTCCTAC
C

>P. maximowiczii _NRAMP3.2_CDS
ATGCCTGTAGAAGAAAACCACCAACCTTTATTGCAAGAAGAAGAAGAAAGAGCTTATGATTCTG
ATGAGAAAGTGCTCATAATTGGGGTTGATTCTGACACGGAAAGCGGTGGCTCAACGGTGTTGCC
ACCGTTTTCATGGAAAAAGTTATGGTTGTTTACTGGTCCTGGGTTTTTAATGTCCATTGCGTTTTT
GGATCCTGGGAATTTGGAAGGGGATCTTCAGGCTGGTGCAATCGCAGGCTACTCTTTGCTTTGGC
TTCTTTTATGGGCTACTGCTATGGGGTTGTTGGTGCAGTTGCTTTCAGCGAGGCTTGGAGTGGCT
ACAGGGAGACATTTAGCTGAGCTGTGTAGAGAAGAGTATCCAACCTGGGCTTCAATGGTTTTGT
GGATTATGGCTGAGTTGGCTTTGATTGGTGCTGATATACAAGAGGTTATTGGAAGTGCTATTGCT
ATTAAGATCTTGAGTAATGGGTTTGTGCCTTTGTGGGCTGGTGTTACTATTACTGCTTGTGATTGC
TTCATCTTCCTATTTCTAGAGAACTACGGTGTGAGAAAATTGGAGGCTGTATTTGCGGTCCTTATT

GGAATAATGGCAGTTACATTTGGATGGATGTTTGCAGATGCAAAACCCAGTGCCTCCGAACTTTT
TCTGGGTATCTTAATTCCAAAACTTAGCTCCAGAACAATACAACAGGCTGTTGGAGTTGTGGGTT
GCATTATCATGCCTCACAATGTGTTCTTGCATTCTGCTCTTGTACAGTCAAGGGAGATCGACCAC
AATAAGAAAGACCGGGTTCAAGAAGCTCTCAGATACTACTCCATAGAGTCAACCACTGCCCTTG
TAATATCGTTCGTAATCAATTTGTTTGTGACGACTGTTTTTGCTAAAGGTTTCTACGGGACAGAA
CTGGCCAATAGTATTGGCCTTGTAAATGCAGGGCAATATCTTCAAGACAAATACGGGGGTGGAT
TTTTCCCAATTTTATACATCTGGGGTATTGGATTATTAGCAGCTGGCCAAAGCAGCACCATTACT
GGCACTTATGCAGGACAGTTTATCATGGGAGGTTTCCTGAACTTGAGGTTAAAGAAATGGCTAA
GGGCATTGATCACTCGAAGCTGTGCTATCATCCCAACTATGATTGTTGCACTTGTTTTTGATACCT
CCGAAGACTCACTAGATGTTCTGAATGAATGGCTAAATGTGCTGCAGTCAATACAGATTCCTTTT
GCACTCATCCCTCTTCTCTGCTTAGTATCCAAGGAGCAAATCATGGGCACTTTCAAAATCGGCCC
CATTCTTAAGATGGTAGCTTGGCTTGTGGCTGCCCTGGTGATGGTAATCAATGGTTACCTTTTGCT
CGACTTTTTCTTCAATGAAGTGACTGGAGTAGCGTTTACCACTGTAGTATGCGGTTTTACAGGTG
CATATGCTGCGTTTATAATTTATCTCATTTCTAGGGGCTTTACATGTTTCTCCCGGTGTTGTCCAT
CTAAACAGATAGAAGTAGAG

>P. euphratica_NRAMP3.1_CDS
ATGCCTTTACCAGAAGAAGACCCCCAACCTTTATTAGAAGACCAAGAAGAAACAGCTTATGATT
CTAACGGGAAAGTCCTTTCGTTTGGGATTGATTATGACACCGAAAGTGGTGGCTCAACGGTGGT
GCCGTCATTTTCATGGAGAAAATTATGGTTGTTCACTGGTCCTGGGTTTTTAATGTGCATTGCTTT
TTTGGACCCTGGCAATTTGGAAGGGGATCTTCAGGCTGGTGCAATTGCAGGATATTCTTTGCTTT
GGCTTCTCTTATGGGCTACTGCTATGGGTTTGTTGGTGCAGTTGCTGTCAGCGAGGCTTGGAGTG
GCTACAGGAAGGCATTTGGCTGAGCTATGGAGAGAAGAGTAnCCAACTTGGGCTCGAATGATTT
TGTGGATTATGGCTGAGTTGGCTTTGATTGGTGCTGATATACAAGAAGTTATTGGGAGTGCTATT
GCTATTCAGATTTTGAGTAATGGGGTTTTGCCTTTGTGGGCTGGTGTTATTATTACTGCCTCCGAT
AGCTTTATCTTCCTATTTCTTGAGAACTACGGTGTGAGGAAATTGGAGGCTGCTTTTGGGATTCTC
ATTGGAATAATGGCAGTGACATTTGCGTGCATGTTTGCTGATGCAAAACCCAGTGCCCCCGAACT
TTTTCTAGGCATCTTAATTCCAAAACTTAGCTCCAAAACAATAAAACAGGCAGTTGGAGTTGTGG
GTTGCATTATCATGCCTCACAATGTGTTCTTGCATTCTGCTCTTGTACAGTCAAGGGAGATTGACC
ACAATAAGAAAGGCCGGGTTCAAGAAGCTCTCAGATACTACTCCATAGAGTCAACTGCTGCCCT
TGCAATATCATTCATGAACAATTTGTTTGTGACAACCGTGTTTGCTAAAGGTTTCCACAGGACAG
AACTGGCCAATAGTATTGGCCTTGTAAATGCAGGGCAATATCTTCAGGATAAATACGGGGGTGG
ATTTTTCCCAATTTTATACATCTGGGGTATTGGGTTATTAGCAGCTGGCCAAAGTAGCACCATTA
CTGGCACTTATGCAGGGCAGTTTATCATGGGAGGTTTCCTGAACTTGGGGTTAAAGAAATGGCTG
AGGGCATTGATTACTCGAAGCTGTGCTATCGTCCCAACTATAATTGTTGCACTTGTTTTTGATACT
TCTGAAGACTCACTAGATGTTCTGAATGAATGGCTAAATATGCTTCAGTCTATTCAGATTCCTTTT
GCACTCATCCCTCTTCTTTGCTTGGTCTCTAAGGAGCAAATCATGGGCACTTTCACAGTTGGCCCC
ATTCTTAAGATGGTTTCTTGGTTTGTAGCTGCCTTGGTGATGCTAATCAATGGTTACCTTTTGCTT
GACTTTTTCTCCAATGAAGTAACTGGAGTAGTGTTTATCACTGTGGTATGCGCTTTTACAGGAGC
ATATGTTACGTTTACAATTTATCTCATTTCTAGGGAAGTTACCATTTCCACTTGGTACTGTCCCTC
C

>P. euphratica_NRAMP3.2_CDS
ATGCCTGTAGAAGAAAACCAACAACTTTTATTGCAAGAAGAAGAAGAAGATAGAGCTTATGATT
CTGATGAGAAAGTGCTCATAATTGGGGTTGATTCTGACACTGAAAGCAGTGGCTCAACGGTGTT
GCCACCGTTTTCATGGAAAAAGTTATGGTTATTTACTGGTCCTGGGTTTTTAATGTCCATTGCGTT
TTTGGATCCTGGAAATTTGGAAGGGGATCTTCAGGCTGGTGCAATCGCAGGCTACTCTTTGCTTT
GGCTTCTTTTATGGGCTACTGCTATGGGGTTGTTGGTGCAGTTGCTTTCAGCGAGGCTTGGAGTG
GCTACAGGAAGGCATTTAGCTGAGCTGTGTAGAGAAGAGTATCCAACCTGGGCTTCAATGGTTT
TGTGGATTATGGCTGAGTTGGCTTTGATTGGTGCTGATATACAAGAGGTTATTGGAAGTGCTATT

GCTATTAAGATCTTGAGTAATGGGTTTTTCCTTTGTGGGCTGGTGTTACTATTACTGCTTGTGAT
TGCTTCATCTTCCTATTTCTAGAGAACTACGGTGTGAGAAAATTGGAGGCTGTATTTGCGGTACT
TATTGGAATAATGGCAGTTACATTTGGATGGATGTTTGGAGATGCAAAACCCAGTGCCTCCGAA
CTTTTTCTGGGTATCTTAATTCCAAAACTTAGCTCCAGAACAATACAACAAGCTGTTGGAGTTGT
GGGTTGCATTATCATGCCTCACAATGTGTTCTTGCATTCTGCTCTTGTACAGTCAAGGGAGATCG
ACCACAGTAAGAAAGACCGGGTTCAAGAAGCTCTCAGATACTACTCCATAGAGTCAACCACTGC
CCTTGTAATATCGTTCCTAATCAATTTGTTTGTGACAACTGTTTTTGCTAAAGGTTTCTACGGGAC
AGAACTGGCCAATAGTATTGGCCTTGTAAATGCAGGGCAATATCTTCAAGACAAATACGGGGGT
GGATTTTTCCCAATTTTATACATCTGGGGTATTGGATTATTAGCAGCTGGCCAAAGCAGCACCAT
TACTGGCACTTATGCAGGACAGTTTATCATGGGAGGTTTCCTGAACTTGAGGTTAAAGAAATGGC
TAAGGGCATTGATCACTCGAAGCTGTGCTATCATCCCAACTATGATTGTTGCACTTGTTTTTGATA
CCTCAGAAGACTCACTAGATGTTCTGAATGAATGGCTAAATGTGCTTCAGTCAATACAGATTCCT
TTTGCACTCATCCCTCTTCTCTGCTTAGTATCCAAGGAGCAAATCATGGGCACTTTCAAAATCGG
CCCCATTCTTCAGATGGTATCTTGGCTTGTGGCTGCCCTGGTGATAGTAATCAATGGTTACCTTTT
GCTTGACTTTTTCGTAAATGAAGTGACCGGAGTAGCGTTTACCACTGTAGTATGCGGTTTTACAG
GTGCATACGTTGTGTTTATAATTTATCTCATTTCTAGGGGCGTTACATGTTTCTCCTGGTGCTGTC
CATCAAAACAGATAGAAGTAGAG

>P. ussuriensis_NRAMP3.1_CDS
ATGCCTTTACCAGAAGAAGACCCACAACCTTTATTAAAAGACCAAGAAGAAACAGCTTATGATT
CTGACGGGAAAGTCCTTTCATTTGGGATTGATTATGACACCGAAAGCGGTGGCTCAACGGTGGT
GCCATCATTTTCATGGAGAAAATTATGGTTGTTCACTGGTCCTGGGTTTTTAATGTGCATTGCTTT
TTTGGACCCTGGCAATTTGGAAGGGGATCTTCAGGCTGGTGCAATTGCAGGGTATTCTTTGCTTT
GGCTTCTCTTATGGGCTACTGCTATGGGTTTGTTGGTGCAGTTGCTGTCAGCAAGGCTTGGAGTG
GCTACAGGAAGGCATTTGGCTGAGCTATGTAGAGAAGAGTATCCAACTTGGGCTCGAATGATTT
TGTGGATTATGGCTGAGTTGGCTTTGATTGGTGCTGATATACAAGAAGTTATTGGGAGTGCTATT
GCTATTCAGATTTTGAGTAATGGGGTTTTGCCTTTGTGGGCTGGTGTTATTATTACTGCTTCCGAT
TGCTTTATCTTCCTATTTCTTGAGAACTACGGTGTGAGGAAATTGGAGGCTGCTTTTGGGATTCTC
ATTGGAATAATGGCAGTGACATTTGCGTGGATGTTTGCTGATGCAAAACCCAGTGCCCCCGAACT
TTTTCTGGGCATCTTAATTCCAAAACTTAGCTCCAAAACAATAAAACAGGCTGTTGGAGTTGTGG
GTTGCATTATCATGCCTCACAATGTGTTCTTGCATTCTGCTCTTGTACAGTCAAGGGAGATTGACC
ACAATAAGAAAGGCCAGGTTCAAGAAGCTCTCAGATACTACTCCATAGAGTCAACTGCTGCCCT
TGCAATATCATTCATGATCAATTTGTTTGTGACGACTGTTTTTGCTAAAGGTTTCCACGGGACAG
AACTGGCCAATAGTATTGGCCTTGTAAATGCAGGGCAATATCTTCAAGATAAATACGGGGGTGG
ATTTTTCCCAATTTTATACATCTGGGGTATTGGGTTATTAGCAGCTGGCCAAAGTAGCACCATTA
CTGGCACTTATGCAGGGCAGTTTATCATGGGAGGTTTCCTGAACTTGGGGTTAAAGAAATGGCTG
AGGGCATTGATTACTCGAAGCTGTGCTATCATCCCAACTATAATTGTTGCACTTGTTTTTGATACT
TCTGAAGACTCACTAGATGTTCTGAATGAATGGCTAAATATGCTTCAGTCTATTCAGATTCCTTTT
GCACTCATCCCTCTTCTTTGCTTGGTCTCTAAGGAGCAAATCATGGGCACTTTCACAGTTGGCCCC
ATTCTTCAGATGGTTTCTTGGCTTGTAGCTGCCTTGGTGATGCTAATCAATGGTTACCTTTTGCTT
GACTTTTTCTCCAATGAAGTGACTGGAGTAGTGTTTACCACTGTGGTATGCGCTTTTACAGGAGC
ATATGTTACGTTTATAATTTATCTCATTTCTAGGGAAGTTACCATTTCCACTTGGTACTGTCCTAC
C

>P. ussuriensis_NRAMP3.2_CDS
ATGCCTGTAGAAGAAAACCACCAACCATTATTGCAAGAAGAAGAAGAAAGAGCTTATGATTCTG
ATGAGAAAGTGCTCATAATTGGGGTTGATTCTGACACGGAAAGCGGTGGCTCAACGGTGTTGCC
ACCGTTTTCATGGAAAAAGTTATGGTTGTTTACTGGTCCTGGGTTTTTAATGTCCATTGCGTTTTT
GGATCCTGGGAATTTGGAAGGGGATCTTCAGGCTGGTGCAATCGCAGGCTACTCTTTGCTTTGGC
TTCTTTTATGGGCTACTGCTATGGGGTTGTTGGTGCAGTTGCTTTCAGCGAGGCTTGGAGTGGCT

ACAGGGAGACATTTAGCTGAGCTGTGTAGAGAAGAGTATCCAACCTGGGCTTCAATGGTTTTGT
GGATTATGGCTGAGTTGGCTTTGATTGGTGCTGATATACAAGAGGTTATTGGAAGTGCTATTGCT
ATTAAGATCTTGAGTAATGGGTTTGTGCCTTTGTGGGCTGGTGTTACTATTACTGCTTGTGATTGC
TTCATCTTCCTATTTCTAGAGAACTACGGTGTGAGAAAATTGGAGGCTGTATTTGCGGTCCTTATT
GGAATAATGGCAGTTACATTTGGATGGATGTTTGCAGATGCAAAACCCAGTGCCTCCGAACTATT
TCTGGGTATCTTAATTCCAAAACTTAGCTCCAGAACAATACAACAGGCTGTTGGAGTTGTGGGTT
GCATTATCATGCCTCACAATGTGTTCTTGCATTCTGCTCTTGTACAGTCAAGGGAGATCGACCAC
AATAAGAAAGACCGGGTTCAAGAAGCTCTCAGATACTACTCCATAGAGTCAACCACTGCCCTTG
TAATATCGTTCGTAATCAATTTGTTTGTGACGACTGTTTTTGCTAAAGGTTTCTACGGACAGAA
CTGGCCAATAGTATTGGCCTTGTAAATGCAGGGCAATATCTTCAAGACAAATACGGGGGTGGAT
TTTTCCCAATTTTATACATCTGGGGTATTGGATTATTAGCAGCTGGCCAAAGCAGCACCATTACT
GGCACTTATGCAGGACAGTTTATCATGGGAGGTTTCCTGAACTTGAGGTTAAAGAAATGGCTAA
GGGCATTGATCACTCGAAGCTGTGCTATCATCCCAACTATGATTGTTGCACTTGTTTTTGATACCT
CCGAAGACTCACTAGATGTTCTGAATGAATGGCTAAATGTGCTGCAGTCAATACAGATTCCTTTT
GCACTCATCCCTCTTCTCTGCTTAGTATCCAAGGAGCAAATCATGGGCACTTTCAAAATCGGCCC
CATTCTTAAGATGGTAGCTTGGCTTGTGGCTGCCCTGGTGATGGTAATCAATGGTTACCTTTTGCT
CGACTTTTTCTTCAATGAAGTGACCGGAGTAGCGTTTACCACTGTAGTATGCGGTTTTACAGGTG
CATATGCTGCGTTTATAATTTATCTCATTTCTAGGGGCTTTACATGTTTCTCCCGGTGCTGTCCAT
CTAAACAGATAGAAGTAGAG

>P. nigra_NRAMP3.1_CDS
ATGCCTGTACCAGAAGAAGACCCACAACCTTTATTAAAAGACCAAGAAGAAACAGCTTATGATT
CTGACGGGAAAGTTCTTTCATTTGGGATTGATTATGACACCGAAAGCGGTGGCTCAACGGTGGT
GCCATCATTTTCATGGAGAAAATTATGGTTGTTCACTGGTCCTGGGTTTTTAATGTGCATTGCTTT
TTTGGACCCTGGCAATTTGGAAGGGGATCTTCAGGCTGGTGCAATTGCAGGGTATTCTTTGCTTT
GGCTTCTCTTATGGGCTACTGCTATGGGTTTGTTGGTGCAGTTGCTGTCAGCAAGGCTTGGAGTG
GCTACAGGAAGGCATTTAGCTGAGCTATGTAGAGAAGAGTATCCAACTTGGGCTCGAATGATTT
TGTGGATTATGGCTGAGTTGGCTTTGATTGGTGCTGATATACAAGAAGTTATTGGGAGTGCTATT
GCTATTCAGATTTTGAGTAATGGGGTTTTGCCTTTGTGGGCTGGTGTTATTATTACTGCTTCTGAT
TGCTTTATCTTCCTATTTCTTGAGAACTACGGTGTGAGGAAATTGGAGGCTGCTTTTGGGATTCTC
ATTGGAATAATGGCAGTGACATTTGCGTGGATGTTTGCTGATGCAAAACCCAGTGCCCCCGAACT
TTTTCTAGGCATCTTAATTCCAAAACTTAGCTCCAAAACAATAAAACAGGCTGTTGGAGTTGTGG
GTTGCATTATCATGCCTCACAATGTGTTCTTGCATTCTGCTCTTGTACAGTCAAGGGAGATTGACC
ACAATAAGAAAGGCCAGGTTCAAGAAGCTCTTAGATACTACTCCATAGAGTCAACTGCTGCCCT
TGCAATATCATTCATGATCAATTTGTTTGTGACGACCGTTTTTGCTAAAGGTTTTCACGGACAG
AACTGGCCAATAGTATTGGCCTTGTAAATGCAGGGCAATATCTTCAAGACAAATACGGGGGTGG
GTTTTTCCCAATTTTATACATCTGGGGTATTGGGTTATTAGCAGCTGGCCAAAGCAGCACCATTA
CTGGCACTTATGCAGGGCAGTTTATCATGGGAGGTTTCCTGAACTTGGGGTTAAAGAAATGGCTG
AGGGCATTGATTACTCGAAGCTGTGCTATCATCCCAACTATAATTGTTGCACTTGTTTTTGATACT
TCTGAAGACTCGCTAGATGTTCTGAATGAATGGCTAAATATGCTTCAGTCTATTCAGATTCCTTTC
GCACTCATCCCTCTTCTTTGCTTGGTCTCGAAGGAGCAAATCATGGGCACTTTCACAGTTGGCCC
CATTCTTCAGATGGTTTCTTGGCTTGTAGCTGCCTTGGTGATGCTAATCAATGGTTACCTTTTGCT
TGACTTTTTCTCCAATGAAGTAACTGGAGTAGCGTTTACCACTGTGGTATGCGCTTTTACAGGAG
CATATGTTGCGTTTATAATTTATCTCATTTCTAGGGAAGTTACCATTTCCACTTGGTACTGTCCCA
CC

>P. nigra_NRAMP3.2_CDS
ATGCCTGTAGAAGAAAACCACCAACCTTTATTGCAAGAAGAAGAAGAAAGAGCTTATGATTCTG
ATGAGAAAGTGCTCATAATTGGGGTTGATTCTGACACGGAAAGCGGTAGCTCAACGGTGTTGCC
ACCGTTTTCATGGAAAAAGTTATGGTTGTTTACTGGTCCTGGGTTTTTAATGTCCATTGCGTTTTT

GGATCCTGGGAATTTGGAAGGGGATCTTCAGGCTGGTGCAATCGCAGGCTACTCTTTGCTTTGGC
TTCTTTTATGGGCTACTGCTATGGGGTTGTTGGTGCAGTTGCTTTCAGCGAGGCTTGGAGTGGCT
ACAGGGAGGCATTTAGCTGAGCTGTGTAGAGAAGAGTATCCAACCTGGGCTTCAATGGTTTTGT
GGATTATGGCTGAGTTGGCTTTGATTGGTGCTGATATACAAGAGGTTATTGGAAGTGCTATTGCT
ATTAAGATCTTGAGTAATGGGTTTGTGCCTTTGTGGGCTGGTGTTACTATTACTGCTTGTGATTGG
TTCATATTCCTATTTCTAGAGAACTACGGTGTGAGAAAATTGGAGGCTGTATTTGCGGTCCTTAT
TGGAATAATGGCAGTTACATTTGGATGGATGTTTGCAGATGCAAAACCCAGTGCCTCCGAACTTT
TTCTAGGTATCTTAATTCCAAAACTTAGCTCCAGAACAATACAACAGGCTGTTGGAGTTGTGGGT
TGCATTATCATGCCTCACAATGTGTTCTTGCATTCTGCTCTTGTACAGTCAAGGGAGATCGACCA
CAATAAGAAAGACCGGGTTCAAGAAGCTCTCAGATACTACTCCATAGAGTCAACCACTGCCCTT
GTAATATCGTTCGTAATCAATTTGTTTGTGACGACTGTTTTTGCTAAAGGTTTCTACGGGACAGA
ACTGGCCAATAGTATTGGCCTTGTAAATGCAGGGCAATATCTTCAAGACAAATACGGGGGTGGA
TTTTTCCCAATTTTATACATCTGGGGTATTGGATTATTAGCAGCTGGCCAAAGCAGCACCATTAC
TGGGACCTATGCAGGACAGTTTATCATGGGAGGTTTCCTGAACTTGAGGTTAAAGAAATGGCTA
AGGGCACTGATCACTCGAAGCTGTGCTATCATCCCAACTATGATTGTTGCACTTGTTTTTGATAC
CTCCGAAGACTCACTAGATGTTCTGAATGAATGGCTAAATGTGCTGCAGTCAATACAGATTCCTT
TTGCACTCATCCCTCTTCTCTGCTTAGTATCCAAGGAGCAAATCATGGGCACTTTCAAAATCGGC
CCCACTCTTCAGATGGTATCTTGGCTTGTGGCTGCCCTGGTGATGGTAATCAATGGTTACCTTTTA
CTCGACTTTTTCTTCAATGAAGTGACCGGAGTAGCGTTTACCACTGTAGTATGCGGTTTTACAGG
TGCATATGTTGCGTTTATAATTTATCTCATATCTAGGGGCTTTACGTGTTTCTCCCGGTGCTGTTC
ATCTAAACAGATAGAAGTAGAG

>P. deltoides_NRAMP3.1_CDS
ATGCCTTTACCAGAAGAAGACCCACAACCTTTATTAAAAGACCAAGAAGAAACAGCTTATGATT
CTGACGGGAAAGTCCTTTCATTTGGGATTGATTATGACACCGAAAGCGGTAGCTCAACGGTGGT
GCCATCATTTTCATGGAGAAAATTATGGTTGTTCACTGGTCCTGGGTTTTTAATGTGCATTGCTTT
TTTGGACCCTGGCAATTTGGAAGGGGATCTTCAGGCTGGTGCAATTGCAGGATATTCTTTGCTTT
GGCTTCTCTTATGGGCTACTGCTATGGGTTTGTTGGTGCAGTTGCTGTCAGCAAGGCTTGGAGTG
GCTACAGGAAGGCATTTGGCTGAGCTATGTAGAGAAGAGTATCCAACTTGGGCTCGAATGATTT
TGTGGATTATGGCTGAGTTGGCTTTGATTGGTGCTGATATACAAGAAGTTATTGGGAGTGCTATT
GCTATTCAGATTTTGAGTAATGGGGTTTTGCCTTTGTGGGCTGGTGTTATTATTACTGCTTCCGAT
TGCTTTATCTTCCTATTTCTTGAGAACTACGGTGTGAGGAAATTGGAGGCTGCTTTTGGGATTCTC
ATTGGAATAATGGCAGTGACATTTGCGTGGATGTTTGCTGATGCAAAACCCAGTGCCCCCGAACT
TTTTCTGGGCATTTTAATTCCAAAACTTAGCTCCAAAACAATAAAACAGGCTGTTGGAGTTGTGG
GTTGCATTATCATGCCTCACAATGTGTTCTTGCATTCTGCTCTTGTACAGTCAAGGGAGATTGACC
ACAATAAGAAAGGCCAGGTTCAAGAAGCTCTCAGATACTACTCCATAGAGTCAACTGCTGCCCT
TGCAATATCATTCATGATCAATTTGTTTGTGACGACCGTTTTTGCTAAAGGTTTTCATGGGACAG
AACTGGCCAATAGTATTGGCCTTGTAAATGCAGGGCAATATCTTCAAGATAAATACGGGGGTGG
ATTTTTCCCAATTTTATACATCTGGGGTATTGGGTTATTAGCAGCTGGCCAAAGTAGCACCATTA
CTGGCACTTATGCAGGGCAGTTTATCATGGGAGGTTTCCTGAACTTGGGGTTAAAGAAATGGCTG
AGGGCATTGATTACTCGAAGCTGTGCTATCATCCCAACTATAATTGTTGCACTTGTTTTTGATACT
TCTGAAGACTCACTAGATGTTCTGAATGAATGGCTAAATATGCTTCAGTCTATTCAGATTCCTTTT
GCACTCATCCCTCTTCTTTGCTTGGTCTCTAAGGAGCAAATCATGGGCACTTTCACAGTTGGCCCC
ATTCTTCAGATGGTTTCTTGGCTTGTAGCTGCCTTGGTGATGTTAATCAATGGTTACCTTTTGCTT
GACTTTTTCTCCAATGAAGTAACTGGAGTAGTGTTTACCACTGTGGTATGCGCTTTTACAGGAGC
ATATGTTACGTTTATAATTTATCTCATTTCTAGGGAAGTTACCATTTCCACTTGGTACTGTCCCAC
C

>P. deltoides_NRAMP3.2_CDS

ATGCCAGTAGAAGAAAACCACCAACCTTTATTGCAAGAAGAAGAAGAAAGAGCTTATGATTCTG
ATGAGAAAGTGCTCATAATTGGGGTTGATTCTGACACGGAAAGCGGTGGCTCAACGGTGTTGCC
ACCGTTTTCATGGAAAAAGTTATGGTTGTTTACTGGTCCTGGGTTTTTAATGTCCATTGCGTTTTT
GGATCCTGGAAATTTGGAAGGGGATCTTCAGGCTGGTGCAATCGCAGGCTACTCTTTGCTTTGGC
TTCTTTTATGGGCTACTGCTATGGGGTTGTTGGTGCAGTTGCTTTCAGCGAGGCTTGGAGTGGCT
ACAGGGAGGCATTTAGCTGAGCTGTGTAGAGAAGAGTATCCAACCTGGGCTTCAATGGTTTTGT
GGATTATGGCTGAGTTGGCTTTGATTGGTGCTGATATACAAGAGGTTATTGGAAGTGCTATTGCT
ATTAAGATCTTGAGTAATGGGTTTGTGCCTTTGTGGGCTGGTGTTACTATTACTGCTTGTGATTGC
TTCATCTTCCTATTTCTAGAGAACTACGGTGTGAGAAAATTGGAGGCTGTATTTGCGGTCCTTATT
GGAATAATGGCAGTTACATTTGGATGGATGTTTGCAGATGCAAAACCCAGTGCCTCCGAACTTTT
TCTGGGTATCTTAATTCCAAAACTTAGCTCCAGAACAATACAACAGGCTGTTGGAGTTGTGGGTT
GCATTATCATGCCTCACAATGTGTTCTTGCATTCTGCTCTTGTACAGTCAAGGGAGATTGACCAC
AATAAGAAAGGCCGGGTTCAAGAAGCTCTCAGATACTACTCCATAGAGTCAACCACTGCCCTTG
TAATATCGTTCGTAATCAATTTGTTTGTGACGACTGTTTTTGCTAAAGGTTTCTACGGGACAGAA
CTGGCCAATAGTATTGGCCTTGTAAATGCAGGGCAATATCTTCAAGACAAATACGGGGGTGGAT
TTTTCCCAATTTTATACATCTGGGGTATTGGATTATTAGCAGCTGGCCAAAGCAGCACCATTACT
GGCACTTATGCAGGACAGTTTATCATGGGAGGTTTCCTGAACTTGAGGTTAAAGAAATGGCTAA
GGGCATTGATCACTCGAAGCTGTGCTATAATCCCAACTATGATTGTTGCACTTGTTTTTGATACCT
CCGAAGACTCACTAGATGTTCTGAATGAATGGCTAAATGTGCTGCAGTCAATACAGATTCCTTTT
GCACTCATCCCTCTTCTCTGCTTAGTATCCAAGGAGCAAATCATGGGCACTTTCAAAATCGGCCC
CATTCTTCAGATGGTAGCTTGGCTTGTGGCTGCCCTGGTGATGGTAATCAATGGTTACCTTTTACT
CGACTTTTTCTTCAATGAAGTGACCGGAGTAGCGTTTACCACTGTAGTATGCGGTTTTACAGGTG
CATATGCTGCGTTTATAATTTATCTCATATCTAGGGGCTTTACATGTTTCTCCCGGTGCTGTCCAT
CCAAACAGATAGAAGTAGAG

>P. tremula_NRAMP3.1_CDS
ATGCCTTTACCAGAAGAAGACCCACAACCTTTATTAAAAGACCAAGAAGAAACAGCTCATGATT
CTGACGGGAAAGTCCTTTCATTTGGGATTGATTATGACACCGAAAGCGGTGGCTCAACGGTGGT
GCCATCATTTTCATGGAGAAAATTATGGTTGTTCACTGGTCCTGGGTTTTTAATGTGCATTGCTTT
TTTGGACCCTGGCAATTTGGAAGGGGATCTTCAGGCTGGTGCAATTGCAGGGTATTCTTTGCTTT
GGCTTCTCTTATGGGCTACTGCTATGGGTTTGTTGGTGCAGTTGCTGTCAGCAAGGCTTGGAGTG
GCTACAGGAAGGCATTTGGCTGAGCTATGTAGAGAAGAGTATCCAACCTGGGCTCGAATGATTT
TGTGGATTATGGCTGAGTTGGCTTTGATTGGTGCTGATATACAAGAAGTTATTGGGAGTGCTATA
GCTATTCAGATTTTGAGTAATGGGGTTTTGCCTTTGTGGGCTGGTGTTATTATTACTGCTTCCGAT
TGCTTTATCTTCCTATTTCTTGAGAACTACGGTGTGAGGAAACTGGAGGCTGCTTTTGGGATCCTC
ATTGGAATAATGGCAGTGGCATTTGCGTGGATGTTTGCTGATGCAAAACCCAGTGCCCCTGAACT
TTTTCTGGGCATCTTAATTCCAAAACTTAGCTCCAAAACAATAAAACAGGCTGTCGGAGTTGTGG
GTTGCATTATCATGCCTCACAATGTGTTCTTGCATTCTGCTCTTGTACAGTCAAGGGAGATTGACT
GCAATAAGAAAGGCCAGGTTCAAGAAGCTCTCAGATACTACTCCATAGAGTCAACTGCTGCCCT
TGCAATATCATTCATGATCAATTTGTTTGTGACGACCATTTTTGCTAAAGGTTTCCACGGGACAG
AACTGGCCAATAGTATTGGCCTTGTAAATGCAGGGCAATATCTTCAAGATAAATACGGGGGTGG
ATTTTTCCCAATTTTATACATCTGGGGTATTGGGTTATTAGCAGCTGGCCAAAGTAGCACCATTA
CTGGCACTTATGCAGGGCAGTTTATCATGGGAGGTTTCCTGAACTTGGGATTAAAGAAATGGCTG
AGGGCATTGATTACTCGAAGCTGTGCTATCATCCCAACTATAATTGTTGCACTTGTTTTTGATACT
TCTGAAGACTCACTAGATGTTCTGAATGAATGGCTAAATATGCTTCAGTCTATTCAGATTCCTTTT
GCACTCATCCCTCTTCTTTGCTTGGTCTCCAAGGAGCAAATCATGGGCACTTTCACAGTCGGCCC
CATTCTTAAGATGGTTTCTTGGCTTGTAGCTGCCTTGGTGATGCTAATCAATGGTTACCTTTTGCT
TGACTTTTTCTCCAATGAAGTAACTGGAGTAGTGTTTACCACTGTGGTATGCGCTTTTACAGGAG
CATATGTTACGTTTATAATTTATCTCATTTCTAGGGAAGTATCCATTTCCACTTGGTACTGTCCCA
CC

>P. tremula_NRAMP3.2_CDS
ATGTCTGTAGAAGAAAACCAACAACCTTTATTGCAAGAAGAAGAAGAAAGAGCTTATGATTCTG
ATGAGAAAGTGCTCATAATTGGGGTTGATTCTGACACGGAAAGCGGTGGCTCAACGGTGTTGCC
ACCGTTTTCATGGAAAAAGTTATGGTTGTTTACTGGTCCTGGATTTTTAATGTCCATTGCGTTTTT
GGATCCTGGGAATTTGGAAGGGGATCTTCAGGCTGGTGCAATCGCAGGCTACTCTTTGCTTTGGC
TTCTTTTCTGGGCTACTGCTATGGGGTTGTTGGTGCAGTTGCTTTCAGCGAGGCTTGGAGTGGCTA
CAGGGAGGCATTTAGCTGAGCTGTGTAGAGAAGAGTATCCAACCTGGGCTTCAATGGTTTTGTG
GATTATGGCTGAGTTGGCTTTGATTGGTGCTGATATACAAGAGGTTATTGGAAGTGCTATTGCTC
TTAAGATCTTGAGTAATGGGTTTTTGCCTTTGTGGGCTGGTGTTACTATTACTGCTTGTGATTGCT
TCATCTTCCTATTTCTAGAGAACTACGGTGTGAGAAAATTGGAGGCTGTATTTGCGGTCCTTATT
GGAATAATGGCAGTTACATTTGGATGGATGTTTGCAGATGCAAAACCCAGTGCCTCCGAACTTTT
TCTGGGTATCTTAATTCCCAAACTTAGCTCCAGAACAATACAACAGGCTGTTGGAGTTGTGGGTT
GCATTATCATGCCTCACAATGTGTTCTTGCATTCTGCTCTTGTACAGTCAAGGGAGATCGACCAC
AATAAGAAAGGCCGGGTTCAAGAAGCTCTCAGATACTACTCCATAGAGTCAACCACTGCCCTTG
TAATATCGTTCGTAATCAATTTGTTTGTGACGACTGTTTTTGCTAAAGGTTTCTATGGGACAGAAC
TGGCCAATAGTATTGGCCTTGTAAATGCAGGGCAATATCTTCAAGACAAATACGGGGGTGGATT
TTTCCCAATTTTATACATCTGGGGCATTGGATTATTAGCAGCTGGCCAAAGCAGCACCATTACTG
GCACTTATGCAGGGCAGTTTATCATGGGAGGTTTCCTGAACTTGAGGTTAAAGAAATGGTTAAG
GGCATTGATCACTCGAAGCTGTGCTATCATCCCAACTATGATTGTTGCACTTGTTTTTGATGCCTC
CGAAGACTCACTAGATGTTCTGAATGAATGGCTAAATGTGCTGCAATCAATACAGATTCCTTTTG
CACTCATTCCTCTTCTCTTCTTGGTATCCAAGGAGCAAATCATGGGCACTTTCAAAATCGGTCCC
ATTCTTAAGATGGTATCTTGGCTTGTGGCTGCCCTGGTGATAGTAATCAATGGTTACCTTTTGCTC
GACTTTTTCGTCAATGAAGTGACCGGAGTAGCGTTTACCACTGTAGTATGCGGTTTTACAGGTGC
ATATGTTGCGTTTATAATTTATCTCATTTCTAGGGGCTTTAAATGTTTCTCCTGGTGCTGTCAATC
TAAACAGATAGAAGTAGAG

>P. tremuloides_NRAMP3.1_CDS
ATGCCTTTACCAGAAGAAGACCCACAACCTTTATTAAAAGACCAAGAAGAAACAGCTTATGATT
CTGACGGGAAAGTCCTTTCATTTGGGATTGATTATGACACCGAAAGCGGTGGCTCAACGGTGTTG
CCACCGTTTTCATGGAAAAAGTTATGGTTGTTTACTGGTCCTGGGTTTTTAATGTCCATTGCTTTT
TTGGACCCTGGCAATTTGGAAGGGGATCTTCAGGCTGGTGCAATTGCAGGGTATTCTTTGCTTTG
GCTTCTCTTATGGGCTACTGCTATGGGTTTGTTGGTGCAGTTGCTGTCAGCAAGGCTTGGAGTGG
CTACAGGAAGGCATTTGGCTGAGCTATGTAGAGAAGAGTATCCAACCTGGGCTCGAATGATTTT
GTGGATTATGGCTGAGTTGGCTTTGATTGGTGCTGATATACAAGAAGTTATTGGGAGTGCTATAG
CTATTCAGATTTTGAGTAATGGGGTTTTGCCTTTGTGGGCTGGTGTTATTATTACTGCTTCCGATT
GCTTTATCTTCCTATTTCTTGAGAACTACGGTGTGAGGAAATTGGAGGCTGCTTTTGGGATCCTC
ATTGGAATAATGGCAGTGACATTTGCGTGGATGTTTGCTGATGCAAAACCCAGTGCCCCTGAACT
TTTTCTGGGCATCTTAATTCCAAAACTTAGCTCCAAAACAATAAAACAGGCTGTCGGAGTTGTGG
GTTGCATTATCATGCCTCACAATGTGTTCTTGCATTCTGCTCTTGTACAGTCAAGGGAGATTGACC
ACAATAAGAAAGGCCAGGTTCAAGAAGCTCTCAGATACTACTCCATAGAGTCAACTGCTGCCCT
TGCAATATCATTCATGATCAATTTGTTTGTGACGACCATTTTTGCTAAAGGTTTCCACGGGACAG
AACTGGCCAATAGTATTGGCCTTGTAAATGCAGGGCAATATCTTCAAGATAAATACGGGGGTGG
ATTTTTCCCAATTTTATACATCTGGGGTATTGGGTTATTAGCAGCTGGCCAAAGTAGCACCATTA
CTGGCACTTATGCAGGGCAGTTTATCATGGGAGGTTTCCTGAACTTGGGATTAAAGAAATGGCTG
AGGGCATTGATTACTCGAAGCTGTGCTATCATCCCAACTATAATTGTTGCACTTGTTTTTGATACC
TCTGAAGACTCACTAGATGTTCTGAATGAATGGCTAAATATGCTTCAGTCTATTCAGATTCCTTTT
GCACTCATCCCTCTTCTTTGCTTGGTCTCCAAGGAGCAACTCATGGGCACTTTCACAGTTGGCCCC
ATTCTTAAGATGGTTTCTTGGCTTGTAGCTGCCTTGGTGATGCTAATCAATGGTTACCTTTTGCTT
GACTTTTTCTCCAATGAAGTAACTGGAGTATTGTTTACCACTGTGGTATGCGCTTTTACAGGAGC
ATATGTTACGTTTATAATTTATCTCATTTCTAGGGAAGTATCCATTTCCACTTGGTACTGTCCCAC
C

>P. tremuloides_NRAMP3.2_CDS
ATGCCTGTAGAAGAAAACCAACAACCTTTATTGCAAGAAGAAGAAGAAAGAGCTTATGATTCTG
ATGAGAAAGTGCTCATAATTGGGGTTGATTCTGACACGGAAAGCGGTGGCTCAACGGTGTTGCC
ACCGTTTTCATGGAAAAAGTTATGGTTGTTTACTGGTCCTGGGTTTTTAATGTCCATTGCGTTTTT
GGATCCTGGGAATTTGGAAGGGGATCTTCAGGCTGGTGCAATCGCAGGCTACTCTTTGCTTTGGC
TTCTTTTATGGGCTACTGCTATGGGGTTGTTGGTGCAGTTGCTTTCAGCGAGGCTTGGAGTGGCT
ACAGGGAGGCATTTAGCTGAGCTGTGTAGAGAAGAGTATCCAACCTGGGCTTCAATGGTTTTGT
GGATTATGGCTGAGTTGGCTTTGATTGGTGCTGATATACAAGAGGTTATTGGAAGTGCTATTGCT
CTTAAGATCTTGAGTAATGGGTTTTTGCCTTTGTGGGCTGGTGTTACTATTACTGCTTGTGATTGC
TTCATCTTCCTATTTCTAGAGAACTACGGTGTGAGAAAATTGGAGGCTGTATTTGCGGTCCTTATT
GGACTAATGGCAGTTACATTTGGATGGATGTTTGCAGATGCAAAACCCAGTGCCTCCGAACTTTT
TCTGGGTATCTTAATTCCCAAACTTAGCTCCAGAACAATACAACAGGCTGTTGGAGTTGTGGGTT
GCATTATCATGCCTCACAATGTGTTCTTGCATTCTGCTCTTGTACAGTCAAGGGAGATCGACCAC
AATAAGAAAGGCCGGGTTCAAGAAGCTCTCAGATACTACTCCATAGAGTCAACCACTGCCCTTG
TAATATCGTTCGTAATCAATTTGTTTGTGACGACTGTTTTTGCTAAAGGTTTCTATGGGACAGAAC
TGGCCAATAGTATTGGCCTTGTAAATGCAGGGCAATATCTTCAAGACAAATACGGGGGTGGATT
TTTCCCAATTTTATACATCTGGGGTATTGGATTATTAGCAGCTGGCCAAAGCAGCACCATTACTG
GCACTTATGCAGGACAGTTTATCATGGGAGGTTTCCTGAACTTGAGGTTAAAGAAATGGTTAAG
GGCATTGATCACTCGAAGCTGTGCTATCATCCCAACTATGATTGTTGCACTTGTTTTTGATGCCTC
CGAAGACTCACTAGATGTTCTGAATGAATGGCTAAATGTGCTGCAGTCAATACAGATTCCTTTTG
CACTCATCCCTCTTCTCTGCTTGGTATCCAAGGAGCAAATCATGGGCACTTTCAAAATCGGCCCC
ATTCTTAAGATGGTATCTTGGCTTGTGGCTGCCCTGGTGATAGTAATCAATGGTTACCTTTTGCTC
GACTTTTTCGTCAATGAAGTGGCCGGAGTAGCGTTTACCACTGTAGTATGCGGTTTTACAGGTGC
ATATGTTGCGTTTATAATTTATCTCATTTCTAGGGGCTTTACATGTTTCTCCTGGTGCTGTCCATCT
AAACAGATAGAAGTAGAG

>P. grandidentata_NRAMP3.1_CDS
ATGCCTTTACCAGAAGAAGACCCACAACCTTTATTAAAAGACCAAGAAGAAACAGCTTATGATT
CTGACGGGAAAGTCCTTTCATTTGGGATTGATTATGACACCGAAAGCGGTGGCTCAACGGTGGT
GCCATCATTTTCATGGAGAAAATTATGGTTGTTCACTGGTCCTGGGTTTTTAATGTGCATTGCTTT
TTTGGACCCTGGCAATTTGGAAGGGGATCTTCAGGCTGGTGCAATTGCAGGGTATTCTTTGCTTT
GGCTTCTCTTATGGGCTACTGCTATGGGTTTGTTGGTGCAGTTGCTGTCAGCAAGGCTTGGGGTG
GCTACAGGAAGGCATTTGGCTGAGCTATGTAGAGAAGAGTATCCAACTTGGGCTCGAATGATTT
TGTGGATTATGGCTGAGTTGGCTTTGATTGGTGCTGATATACAAGAAGTTATTGGGAGTGCTATT
GCCATTCAGATTTTGAGTAATGGGGTTTTGCCTTTGTGGGCTGGTGTTATTATTACTGCTTCCGAT
TGCTTTATCTTCCTATTTCTTGAGAACTACGGTGTGAGGAAATTGGAGGCTGCTTTTGGGATCCTC
ATTGGAATAATGGCAGTGACATTTGCGTGGATGTTTGCTGATGCAAAACCCAGTGCCCCTGAACT
TTTTCTGGGCATCTTAATTCCAAAACTTAGCTCCAAAACAATAAAACAGGCTGTCGGAGTTGTGG
GTTGCATTATCATGCCTCACAATGTGTTCTTGCATTCTGCTCTTGTACAGTCAAGGGAGATTGACC
ACAATAAGAAAGGCCAGGTTCAAGAAGCTCTCAGATACTACTCCATAGAGTCAACTGCTGCCCT
TGCAATATCATTCATGATCAATTTGTTTGTGACGACCATTTTTGCTAAAGGTTTCCACGGGACAG
AACTGGCCAATAGTATTGGCCTTGTAAATGCAGGGCAATATCTTCAAGATAAATACGGGGGTGG
ATTTTTCCCAATTTTATACATCTGGGGTATCGGGTTATTAGCAGCTGGCCAAAGTAGCACCATTA
CTGGCACTTATGCAGGGCAGTTTATCATGGGAGGTTTCCTGAACTTGGGATTAAAGAAATGGCTG
AGGGCATTGATTACTCGAAGCTGTGCTATCATCCCAACTATAATTGTTGCACTTGTTTTTGATACT
TCTGAAGACTCACTAGATGTTCTGAATGAATGGCTAAATATGCTTCAGTCTATTCAGATTCCTTTT
GCACTCATCCCTCTTCTTTGCTTGGTCTCCAAGGAGCAACTCATGGGCACTTTTACAGTTGGCCCC
ATTCTTAAGATGGTTTCTTGGCTTGTAGCTGCCTTGGTGATGCTAATCAATGGTTACCTTTTGCTT
GACTTTTTCTCCAATGAAGTAACTGGAGTAGTGTTTACCACTGTGGTATGCGCTTTTACAGGAGC

GTATGTTACGTTTATAATTTATCTCATTTCTAGGGAAGTATCCATTTCCACTTGGTACTGTCCCAC
C

>P. grandidentata_NRAMP3.2_CDS
ATGCCTGTAGAAGAAAACCAACAACCTTTATTGCAAGAAGAAGAAGAAAGAGCTTATGATTCTG
ATGAGAAAGTGCTCATAATTGGGGTTGATTCTGACACGGAAAGCGGTGGCTCAACGGTGTTGCC
ACCGTTTTCATGGAAAAAGTTATGGTTGTTTACTGGTCCTGGGTTTTTAATGTCAATTGCGTTTTT
GGATCCTGGGAATTTGGAAGGGGATCTTCAGGCTGGTGCAATCGCAGGCTACTCTTTGCTTTGGC
TTCTTTTATGGGCTACTGCTATGGGGTTGTTGGTGCAGTTGCTTTCAGCGAGGCTTGGAGTGGCT
ACAGGGAGGCATTTAGCTGAGCTGTGTAGAGAAGAGTATCCAACCTGGGCTTCAATGATTTTGT
GGATTATGGCTGAGTTGGCTTTGATTGGTGCTGATATACAAGAGGTTATTGGAAGTGCTATTGCT
CTTAAGATCTTGAGTAATGGGTTTTTGCCTTTGTGGGCTGGTGTTACTATTACTGCTTGTGATTGC
TTCATCTTCCTATTTCTAGAGAACTACGGTGTGAGAAAATTGGAGGCTGTATTTGCGGTCCTTATT
GGAATAATGGCAGTTACATTTGGATGGATGTTTGCAGATGCAAAACCCAGTGCCTCCGAACTTTT
TCTGGGTATCTTAATTCCCAAACTTAGCTCCAGAACAATACAACAGGCTGTTGGAGTTGTGGGTT
GCATTATCATGCCTCACAATGTGTTCTTGCATTCTGCTCTTGTACAGTCAAGGGAGATCGACCAC
AATAAGAAAGGCCGGGTTCAAGAAGCTCTCAGATACTACTCCATAGAGTCAACCACTGCCCTTG
TAATATCGTTCGTAATCAATTTGTTTGTGACGACTGTTTTTGCTAAAGGTTTCTATGGGACAGAAC
TGGCCAATAGTATTGGCCTTGTAAATGCAGGGCAATATCTTCAAGACAAATACGGGGGTGGATT
TTTCCCAATTTTATACATCTGGGGTATTGGATTATTAGCAGCTGGCCAAAGCAGCACCATTACTG
GCACTTATGCAGGACAGTTTATCATGGGAGGTTTCCTGAACTTGAGGTTAAAGAAATGGTTAAG
GGCATTGATCACTCGAAGCTGTGCTATCATCCCAACTATGATTGTTGCCCTTGTTTTTGATGCCTC
CGAAGACTCACTAGATGTTCTGAATGAATGGCTAAATGTGCTGCAGTCAATACAGATTCCTTTTG
CACTCATCCCTCTTCTCTGCTTGGTATCCAAGGAGCAAATCATGGGCACTTTCAAAATCGGCCCC
ATTCTTAAGATGGTATCTTGGCTTGTGGCTGCCCTGGTGATAGTAATCAATGGTTACCTTTTGCTC
GACTTTTTCGTCAATGAAGTGGCCGGAGTAGCGTTTACCACTGTAGTATGCGGTTTTACAGGTGC
ATATGTTGCATTTATAATTTATCTCATTTCTAGGGGCTTTACATGTTTCTCCCGGTGCTGTCCATCT
AAACAGATAGAAGTAGAG

>S. purpurea_NRAMP3_CDS
ATGTCTTTAGATGAAAACCAGCAACCTTTATTGCAAGAAGAAGAAGAGAGAGCTTATGATTCTG
ATGAGAAAGTGCTCGTAATTGGGATTGATTCTGACGCCGAAAGCGGTGGAACGGTGTTGCCACC
GTTTTCATGGAAAAAGTTATGGTTGTTTACTGGTCCTGGGTTTTTAATGTCCATTGCGTTTTTGGA
TCCTGGAAATTTGGAAGGGGATCTTCAGGCTGGTGCAATTGCAGGCTACTCTTTGCTTTGGCTCC
TTTTTTGGGCCACTGCTATGGGGTTGTTGGTGCAGTTGCTGTCAGCCAGGCTTGGAGTGGCTACA
GGGAGGCATTTAGCTGAGCTGTGTAGAGAAGAGTATCCAACCTGGGCTAGAATGATTTTGTGGA
TTATGGCTGAGTTAGCTTTGATTGGTGCTGATATACAAGAGGTTATTGGAAGTGCTATTGCTATT
AAGATCTTGAGTAACGGGGTTGTGCCTTTATGGGCTGGTGTTACTATTACTGCTTGTGACAGCTT
CATCTTCCTATTTCTAGAGAACTACGGTGTGAGAAAATTGGAGGCTGTATTTGCGGTCCTCATTG
GAGTCATGGCAGTTACATTTGGATGGATGTTTGCTGATGCCAAACCCAGTGCCTCCGAACTTTTT
CTGGGTATCTTTATTCCAAAACTTAGCTCCAGAACAATACAGCAGGCTGTAGGAGTTGTGGGTTG
CATTATCATGCCTCACAATGTGTTCCTGCATTCTGCTCTTGTACAGTCAAGGGAGATCGACCACA
ATAATAAAGTCCAGGTTCAAGAAGCTCTCAGATACTACTCCATAGAGTCAACCACTGCCCTTGTG
ATATCGTTCGTAATCAATTTGTTCGTGACGACTGTGTTTGCTAAAGGTTTCTATGGGACAGAACT
GGCCAATAGTATTGGCCTTGTAAATGCAGGGCAATATCTTCAAGACAAATACGGGGGTGGATTT
TTCCCAATCTTATACATCTGGGGTATTGGATTATTAGCTGCTGGCCAAAGTAGCACCATTACTGG
GACCTATGCAGGACAGTTTATAATGGGAGGTTTCCTGAACATGAGGTTAAAGAAATGGCTAAGG
GCATTGATCACTCGAAGTTGTGCTATCATCCCAACGATAATTGTTGCTCTTATTTTTGATACCTCT
GAAGATTCTCTAGATGTTCTGAATGAATGGCTAAATGTGCTGCAGTCAATACAGATTCCTTTTGC
ACTCATCCCTCTTCTCTGTTTGGTCTCCAAGGAGCGAATCATGGGCACTTTCAAAATTGGCTCCAT

CCTTAAGGTCGTTTCTTGGCTTGTGGCGGCCCTGGTGATAGTAATCAATGGTTACCTTTTGCTCGA
CTTTTTCTTCAATGAAGTGACTGGAGTAGCGTTTACCACTGTAGTATGCACTTTTACAGCTGCATA
TGCTGCGTTTATAATTTATCTCACTTCCAGGGGCGTTACATGTTCTTCCTGGCGCGGCCCACCTAA
ACAGATAGAAGTAGAG

>S. suchowensis_NRAMP3_CDS
ATGTCTTTAGATGAAAACCAGCAACCATTATTGCAAGAAGAAGAAGAGAGAGCTTATGATTCTG
ATGAGAAAGTGCTCGTAATTGGGATTGATTCTGACGCCGAAAGCGGTAGAACGGTGTTGCCACC
GTTTTCATGGAAAAAGTTATGGTTGTTTACTGGTCCTGGGTTCTTAATGTCCATTGCGTTTTTGGA
TCCTGGAAATTTGGAAGGGGATCTTCAGGCTGGTGCAATTGCAGGCTACTCTTTGCTTTGGCTCC
TTTTATGGGCCACTGCTATGGGGTTGTTGGTGCAGTTGCTGTCTGCCAGGCTTGGAGTGGCTACA
GGGAGGCATTTAGCTGAGCTGTGTAGAGAAGAGTATCCAACCTGGGCTAGAATGATTTTGTGGA
TTATGGCTGAGTTAGCTTTGATTGGTGCTGATATACAAGAGGTTATTGGAAGTGCTATTGCTATT
AAGATCTTGAGTAACGGGGTTGTGCCTTTATGGGCTGGTGTTACTATTACTGCTTGTGATTGCTTC
ATCTTCCTATTTCTAGAGAACTACGGTGTGAGAAAATTGGAGGCTGTATTTGCGGTCCTCATTGG
AGTCATGGCAGTTACATTTGGATGGATGTTTGCTGATGCCAAACCCAGTGCCTCCGAACTTTTTC
TGGGTATCTTAATTCCAAAACTTAGCTCCAGAACAATACAGCAGGCTGTAGGAGTTGTGGGTTGC
ATTATCATGCCTCACAATGTGTTCCTGCATTCTGCTCTTGTACAGTCAAGGGAGATCGACCACAA
TAATAAAGTCCAGGTTCAAGAAGCTCTCAGATACTACTCCATAGAGTCAACCACTGCCCTTGTGA
TATCGTTCGTAATCAATTTGTTCGTGACGACTGTGTTTGCTAAAGGTTTCTATGGGACAGAACTG
GCCAATAGTATTGGCCTTGTAAATGCAGGGCAATATCTTCAAGACAAATACGGGGGTGGATTTTT
TCCAATCTTATACATCTGGGGTATTGGATTATTAGCTGCTGGCCAAAGTAGCACCATTACTGGGA
CGTATGCAGGACAGTTTATCATGGGAGGTTTCCTGAACATGAGGTTAAAGAAATGGCTAAGGGC
ATTGATCACTCGAAGTTGTGCTATCATCCCAACGATAATTGTTGCGCTTATTTTTGATACCTCTGA
AGATTCTCTAGATGTTCTGAATGAATGGCTAAATGTGCTGCAGTCAATACAGATTCCTTTTGCAC
TCATCCCTCTTCTCTGTTTGGTCTCCAAGGAGCGAATCATGGGCACTTTCAAAATTGGCTCCATCC
TTAAGGTCGTTTCTTGGCTTGTGGCGGCCCTGGTGATAGTAATCAATGGTTACCTTTTGCTCGACT
TTTTCTTCAATGAAGTGACTGGAGTAGCGTTTACCACTGCAGTATGCACTTTCACAGCTGCATAT
GCTGCGTTTATAATTTACCTCACTTCCAGGGGCATTACATGTTCTTCCTGGCGCGGCCCACCTAA
ACAGATAGAAGCAGAG

>S. brachista_NRAMP3_CDS
ATGTCTTTAGATGAAAACCAGCAACCATTATTGCAAGAAGAAGAAGAGAGAGCTTATGATTCTG
ATCAGAAAGTGCTCGTAATTGGGATTGATTCTGACGCCGAAAGCGGTGGCACGGTGTTACCACC
GTTTTCATGGAAAAAGTTATGGTTGTTTACTGGTCCTGGGTTTTTAATGTCCATTGCGTTTTTGGA
TCCTGGAAATTTGGAAGGGGATCTTCAGGCTGGTGCGATTGCAGGCTACTCTTTGCTTTGGCTCC
TTTTATGGGCCACTGCTATGGGGTTGTTGGTGCAGTTGCTGTCAGCCAGGCTTGGAGTGGCTACA
GGGAGGCATTTAGCTGAGCTGTGTAGAGAAGAGTATCCAACCTGGGCTAGAATGATTTTGTGGA
TTATGGCTGAGTTAGCTTTGATTGGTGCTGATATACAAGAGGTTATTGGAAGTGCTATTGCTATT
AAGATCTTGAGTAACGGGGTTGTGCCTTTATGGGCTGGTGTTACTATTACTGCTTGTGATTGCTTC
ATCTTCCTATTTCTAGAGAACTACGGTGTGAGAAAATTGGAGGCTGTATTTGCGGTCCTAATTGG
AGTAATGGCAGTTACATTTGGAATGATGTTTGCTGATGCCAAACCCAGTGCCTCCGAACTTTTTC
TGGGTATCTTAATCCCAAAACTTAGCTCCAGAACAATACAACAGGCTGTAGGAGTTGTGGGTTG
CATTATCATGCCTCACAATGTGTTCCTGCATTCTGCTCTTGTACAGTCAAGGGAGATCGACCACA
ATAATAAAGTCCAGGTTCAAGAAGCTCTCAGATACTACTCCATAGAGTCAACCACTGCCCTTGTA
ATATCGTTCGTAATCAATTTGTTCGTGACGACTGTGTTTGCTCAAGGTTTCTATGGGACAGAACT
GGCCAATAGTATTGGCCTTGTAAATGCAGGGCAATATCTTCAAGACAAATACGGGGGTGGATTT
TTCCCAATCTTATACATCTGGGGTATTGGATTATTAGCTGCTGGCCAAAGTAGCACCATTACAGG
GACCTATGCAGGACAGTTTATCATGGGAGGTTTCCTGAACATGAGGTTAAAGAAATGGCTGAGG
GCATTGATCACTCGAAGTTGTGCTATCATCCCAACGATAATTGTTGCGCTTGTTTTTGAAACCTCT

GAAGAGTCTCTAGATGTTCTGAATGAATGGCTAAATGTGCTGCAGTCAATACAGATTCCTTTTGC
ACTCATCCCTCTTCTCTGTTTGGTCTCCAAGGAGCGAATCATGGGCACTTTCAAAATTGGCTCCAT
TCTTAAGGTCGTTTCTTGGCTTGTGGCGGCCCTGGTGATAATAATCAATGGTTACCTTTTGCTCGA
CTTTTTCTTCAATGAAGTGACTGGAGTAGCGTATACCACTGCAGTATGCACTTTTACAGCTGCAT
ATGCTGCGTTTATAATTTATCTCACTTCTAGGGGCGTTACATGCTCTTCCTGGCGCGGCCCACCTA
AACAGAATCAACAGATAGAAGTAGAG

>S. eriocephala_NRAMP3_CDS
ATGTCTTTAGATGAAAACCAGCAAGCTTTATTGCAAGAAGAAGAAGAGAGAGCTTATGATTCTG
ATGAGAAAGTGCTCGTAATTGGGGTTTATTCTGACGCCGAAAGCGGTGGCACGGTGTTGCCACC
GTTTTCATGGAAAAAGTTATGGTTGTTTACTGGTCCTGGGTTCTTAATGTCCATTGCGTTTTTGGA
TCCTGGAAATTTGGAAGGGGATCTTCAGGCTGGTGCAATTGCAGGCTACTCTTTGCTTTGGCTCC
TCTTATGGGCTACTGCTATGGGGTTGTTGGTGCAGTTGCTGTCAGCCAGGCTTGGAGTGGCTACA
GGGAGGCATTTAGCTGAGCTGTGTAGAGAAGAGTATCCAACCTGGGCTAGAATGATTTTGTGGA
TTATGGCTGAGTTAGCTTTGATTGGAGCTGATATACAAGAGGTTATTGGAAGTGCTATTGCTATT
AAGATCTTGAGTAACGGGGTTGTGCCTTTATGGGCTGGTGTTACTATTACTGCTTGTGATTGCTTC
ATCTTCCTATTTCTAGAGAACTACGGTGTGAGAAAATTGGAGGCTGTATTTGCGGTCCTAATTGG
AGTCATGGCAGTTACATTTGGATGGATGTTTGCTGATGCGAAACCCAGTGCCTCCGAACTTTTTC
TGGGTATCTTAATTCCAAAACTTAGCTCCAGAACAATACAGCAGGCTGTAGGAGTTGTGGGTTGC
ATTATCATGCCTCACAATGTGTTCCTGCATTCTGCTCTTGTACAGTCAAGGGAGATCGACCACAA
TAATAAAATCCAGGTTCAAGAAGCTCTCAGATACTACTCCATAGAGTCAACCACTGCCCTTGTAA
TATCGTTCGTAATCAATTTGTTCGTGACGACTGTGTTTGCTAAAGGTTTCTATGGGACAGAACTG
GCCAATAGTATTGGCCTTGTAAATGCAGGGCAATATCTTCAAGACAAATACGGGGGTGGATTTTT
CCCAATCTTATACATCTGGGGTATTGGATTATTAGCTGCTGGCCAAAGTAGCACCATTACTGGGA
CCTATGCAGGACAGTTTATCATGGGAGGTTTCCTGAACATGAGGTTAAAGAAATGGATAAGGGC
ATTGATCACTCGAAGTTGTGCTATCATCCCAACGATAATTGTTGCGCTTATTTTTGATACCTCTGA
AGATTCTCTAGATGTTCTGAATGAATGGCTAAATGTGCTGCAGTCAATACAGATTCCTTTTGCAC
TCATCCCTCTTCTCTGTTTGGTCTCCAAGGAGCGAATCATGGGCACTTTCAAAATTGGCTCCATCC
TTAAGGTCGTTTCTTGGCTTGTGGCGGCCCTGGTGATGGTAATCAATGGTTACCTTTTGCTCGACT
TTTTCTTCAATGAAGTGACCGGAGTAGCGTTTACCACTGCAGTATGCACTTTTACAGCTGCATAT
GCTGCGTTTATAATTTATCTCACTTCCAGGGGCGTTACATGTTCTTCCTGGCGCGGCCCACCTAAA
CAGATAGAAGTAGAG

>S. sachalinensis_NRAMP3_CDS
ATGTCTTTAGATGAAAACCAGCAACCTTTATTGCAAGAAGAAGAAGAGAGAGCTTATGATTCTG
ATGAGAAAGTGCTCGTAATTGGGATTGATTCTGACGCCGAAAGCGGTGGAACGGTGTTGCCACC
GTTTTCATGGAAAAAGTTATGGTTGTTTACTGGTCCTGGGTTTTTAATGTCCATTGCGTTTTTGGA
TCCTGGAAATTTGGAAGGGGATCTTCAGGCTGGTGCAATTGCAGGCTACTCTTTGCTTTGGCTCC
TTTTATGGGCCACTGCTATGGGGTTGTTGGTGCAGTTGCTGTCAGCCAGGCTTGGAGTGGCTACA
GGGAGGCATTTAGCTGAGCTGTGTAGAGAAGAGTATCCAACCTGGGCTAGAATGATTTTGTGGA
TTATGGCTGAGTTAGCTTTGATTGGTGCTGATATACAAGAGGTTATTGGAAGTGCTATTGCTATT
AAGATCTTGAGTAACGGGGTTGTGCCTTTATGGGCTGGTGTTACTATTACTGCTTGTGATTGCTTC
ATCTTCCTATTTCTAGAGAACTACGGTGTGAGAAAATTGGAGGCTGTATTTGCGGTCCTCATTGG
AGTCATGGCAGTTACATTTGGATGGATGTTTGCTGATGCCAAACCCAGTGCCTCCGAACTTTTTC
TGGGTATCTTAATTCCAAAACTTAGCTCCAGAACAATACAGCAGGCTGTAGGAGTTGTGGGTTGC
ATTATCATGCCTCACAATGTGTTCCTGCATTCTGCTCTTGTACAGTCAAGGGAGATCGACCACAA
TAATAAAGTCCAGGTTCAAGAAGCTCTCAGATACTACTCCATAGAGTCAACCACTGCCCTTGTGA
TATCGTTCGTAATCAATTTGTTCGTGACGACTGTGTTTGCTAAAGGTTTCTATGGGACAGAACTG
GCCAATAGTATTGGCCTTGTAAATGCAGGGCAATATCTTCAAGACAAATACGGGGGTGGATTTTT
CCCAATCTTATACATCTGGGGTATTGGATTATTAGCTGCTGGCCAAAGTAGCACCATTACTGGGA

CCTATGCAGGACAGTTTATCATGGGAGGTTTCCTGAACATGAGGTTAAAGAAATGGCTAAGGGC
ATTGATCACTCGAAGTTGTGCTATCATCCCAACGATAATTGTTGCGCTTATTTTTGATACCTCTGA
AGATTCTCTAGATGTTCTGAATGAATGGCTAAATGTGCTGCAGTCAATACAGATTCCTTTTGCAC
TCATCCCTCTTCTCTGTTTGGTCTCCAAGGAGCGAATCATGGGCACTTTCAAAATTGGCTCCATCC
TTAAGGTCGTTTCTTGGCTTGTGGCGGCCCTGGTGATAGTAATCAATGGTTACCTTTTGCTCGACT
TTTTCTTCAATGAAGTGATTGGAGTAGCGTTTACCACTGCAGTATGCACTTTCACAGCTGCATAT
GCTGCGTTTATAATTTACCTCACTTCCAGGGGCATTACATGTTCTTCCTGGCGCGGCCCACCTAA
ACAGATAGAAGCAGAG

>S. dasyclados_NRAMP3_CDS

ATGGCAGTTACATTTGGATGGATGTTTGCTGATGCCAAACCCAGTGCCTCCGAACTTTTTCTGGG
TATCTTAATTCCAAAACTTAGCTCCAGAACAATACAACAGGCTGTTGGGGTTGTGGGTTGCATTA
TCATGCCTCACAATGTGTTCCTGCATTCTGCTCTTGTACAGTCAAGGGAGATCGACCACAATAAT
AAAGTCCAGGTTCAAGAAGCTGTCAGATACTACTCCATAGAGTCAACCACTGCCCTTGTGATATC
GTTCGTAATCAATTTGTTCGTGACGACTGTGTTTGCTCAAGGTTTCTATGGGACAGAACTGGCCA
ATAGTATTGGCCTTGTAAATGCAGGGCAATATCTTCAAGACAAATACGGGGGTGGATTTTTCCCA
ATCTTATACATCTGGGGTATTGGATTATTAGCTGCTGGCCAAAGTAGCACCATTACTGGGACCTA
TGCAGGACAGTTTATCATGGGAGGTTTCCTGAACATGAGGTTAAAGAAATGGCTAAGGGCATTG
ATCACTCGAAGTTGTGCTATCATCCCAACGATAATTGTTGCGCTTATTTTTGATACCTCTGAAGAT
TCTCTAGATGTTCTGAATGAATGGCTAAATGTGCTGCAGTCAATACAGATTCCTTTTGCACTCAT
CCCTCTTCTCTGTTTGGTCTCCAAGGAGCGAATCATGGGCACTTTCAAAATTGGCTCCATCCTTAA
GGTCGTTTCTTGGCTTGTGGCGGCCCTGGTGATAGTAATCAATGGCTACCTTTTGCTCGACTTTTT
CTTCAATGAAGTGACCGGAGTAGCGTTTACCACCGCAGTATGCACTTTTACAGCTGCATATGCTG
CGTTTATAATTTATCTCACTTCTAGGGGCATTACATGCTCTTCCTGGCGCGGCCCACCTAAACAG
ATAGAAGTAGAG

>S._viminalis_NRAMP3_CDS

ATGTCTTTAGATGAAAACCAGCAACCTTTATTGCAAGAAGAAGAAGAGAGAGCTTATGAT
TCTGATGAGAAAGTGCTCGTAATTGGGATTGATTCTGACGCCGAAAGCAGTGGCACGGTG
TTGCCACCGTTTTCATGGAAAAAGTTATGGTTGTTTACTGGTCCTGGGTTTTTAATGTCC
ATTGCGTTTTTGGATCCTGGAAATTTGGAAGGGGATCTTCAGGCTGGTGCAATTGCAGGC
TACTCTTTGCTTTGGCTCCTTTTATGGGCTACTGCTATGGGGTTGTTGGTGCAGTTGCTG
TCAGCCAGGCTTGGAGTGGCTACAGGGAGGCATTTAGCTGAGCTGTGTAGAGAAGAGTAT
CCAACCTGGGCTAGAATGATTTTGTGGATTATGGCTGAGTTAGCTTTGATTGGTGCTGAT
ATACAAGAGGTTATTGGAAGTGCTATTGCTATTAAGATCTTGAGTAACGGGGTTGTGCCT
TTATGGGCTGGTGTTACTATTACTGCTTGTGATTGCTTCATCTTCCTATTTCTAGAGAAC
TACGGTGTGAGAAAATTGGAGGCTGTATTTGCGGTCCTAATTGGAGTAATGGCAGTTACA
TTTGGATGGATGTTTGCTGATGCCAAACCCAGTGCCTCCGAACTTTTTCTGGGTATCTTA
ATTCCAAAACTTAGCTCCAGAACAATACAGCAGGCTGTTGGGGTTGTGGGTTGCATTATC
ATGCCTCACAATGTGTTCCTGCATTCTGCTCTTGTACAGTCAAGGGAGATCGACCACAAT
AATAAAGTCCAGGTTCAAGAAGCTGTCAGATACTACTCCATAGAGTCAACCACTGCCCTT
GTGATATCGTTCGTAATCAATTTGTTCGTGACGACTGTGTTTGCTCAAGGTTTCTATGGG
ACAGAACTGGCCAATAGTATTGGCCTTGTAAATGCAGGGCAATATCTTCAAGACAAATAC
GGGGGTGGATTTTTCCCAATCTTATACATCTGGGGTATTGGATTATTAGCTGCTGGCCAA
AGTAGCACCATTACTGGGACCTATGCAGGACAGTTTATCATGGGAGGTTTCCTGAACATG
AGGTTAAAGAAATGGCTAAGGGCATTGATCACTCGAAGTTGTGCTATCATCCCAACGATA
ATTGTTGCGCTTATTTTTGATACCTCTGAAGATTCTCTAGATGTTCTGAATGAATGGCTA
AATGTGCTGCAGTCAATACAGATTCCTTTTGCACTCATCCCTCTTCTCTGTTTGGTCTCC

AAGGAGCGAATCATGGGCACTTTCAAAATTGGCTCCATCCTTAAGGTCGTTTCTTGGCTT
GTGGCGGCCCTGGTGATAGTAATCAATGGTTACCTTTTGCTCGACTTTTTCTTCNNNNNN NNG

>S. fargesii_NRAMP3_CDS

ATGTCCATTGCGTTTTTGGATCCTGGAAATTTGGAAGGGGATCTTCAGGCTGGTGCGATT
GCAGGCTACTCTTTGCTTTGGCTCCTTTTATGGGCCACTGCTATGGGGTTGTTGGTGCAG
TTGCTGTCAGCCAGGCTTGGAGTGGCTACAGGGAGGCATTTAGCTGAGCTGTGTAGAGAA
GAGTATCCAACCTGGGCTAGAATGATTTTGTGGATTATGGCTGAGTTAGCTTTGATTGGT
GCTGATATACAAGAGGTTATTGGAAGTGCTATTGCTATTAAGATCTTGAGTAACGGGGTT
GTGCCTTTATGGGCTGGTGTTACTATTACTGCTTGTGATTGCTTCATCTTCCTATTTCTA
GAGAACTACGGTGTGAGAAAATTGGAGGCTGTATTTGCGGTCCTAATTGGAGTAATGGCA
GTTACATTTGGAATGATGTTTGCTGATGCCAAACCCAGTGCCTCCGAACTTTTTCTGGGT
ATCTTAATCCCAAAACTTAGCTCCAGAACAATACAACAGGCTGTAGGAGTTGTGGGTTGC
ATTATCATGCCTCACAATGTGTTCCTGCATTCTGCTCTTGTACAGTCAAGGGAGATCGAC
CACAATAATAAAGTCCAGGTTCAAGAAGCTCTCAGATACTACTCCATAGAGTCAACCACT
GCCCTTGTAATATCGTTCGTAATCAATTTGTTCGTGACGACTGTGTTTGCTCAAGGTTTC
TATGGGACAGAACTGGCCAATAGTATTGGCCTTGTAAATGCAGGGCAATATCTTCAAGAC
AAATACGGGGGTGGATTTTTCCCAATCTTATACATCTGGGGTATTGGATTATTAGCTGCT
GGCCAAAGTAGCACCATTACAGGGACCTATGCAGGACAGTTTATCATGGGAGGTTTCCTG
AACATGAGGTTAAAGAAATGGCTGAGGGCATTGATCACTCGAAGTTGTGCTATCATCCCA
ACGATAATTGTTGCGCTTGTTTTTGAAACCTCTGAAGAGTCTCTAGATGTTCTGAATGAA
TGGCTAAATGTGCTGCAGTCAATACAGATTCCTTTTGCACTCATCCCTCTTCTCTGTTTG
GTCTCCAAGGAGCGAATCATGGGCACTTTCAAAATTGGCTCCATTCTTAAGGTCGTTTCT
TGGCTTGTGGCGGCCCTGGTGATAATAATCAATGGTTACCTTTTGCTCGACTTTTTCTTC
AATGAAGTGACTGGAGTAGCGTATACCACTGCAGTATGCACTTTTACAGCTGCATATGCT
GCGTTTATAATTTATCTCACTTCTAGGGGCGTTACATGCTCTTCCTGGCGCGGCCCACCT
AAACAGAATCAACAGATAGAAGTAGAG

**Supplementary data S2.** Partial genomic sequence of *NRAMP3.1* and *RNAMP3.2* retrieved from *Populus mexicana* genome.

>P. mexicana_NRAMP3.1_genomic
nnnnnnnnnnnnnnnnnnnnnnnnnnnnnnnnnnnnnnnnnnnnnnnnnnnnnnnnnnnnnnnnnnnnnnnnnnnnnnnnnnnnnnnnnn
nnnnnnnnnnnnnnnnnnnnnnnnnnnnnnnnnnnnnnnnnnnnnnnnnnnnnnnnnnnnnnnnnnnnnnnnnnnnnnnnnnnnnnnnn
nnnnnnnnnnnnnnnnnnnnnnnnnnnnnnnnnnnnnnnnnnnnnnnnnnnnnnnnnnnnnnnnnnnnnnnnnnnnnnnnnnnnnnnnn
nnnnnnnnnnnnnnnnnnnnnnnnnnnnnnnnnnnnnnnnnnnnnnnnnnnnnnnnnnnnnnnnnnnnnnnnnnnnnnnnnnnnnnnnn
nnnnnnnnnnnnnnnnnnnnnnnnnnnnnnnnnnnnnnnnnnnnnnnnnnnnnnnnnnnnnnnnnnnnnnnnnnnnnnnnnnnnnnnnn
nnnnnnnnnnnnnnnnnnnnnnnnnnnnnnnnnnnnnnnnnnnnnnnnnnnnnnnnnnnnnnnnnnnnnnnnnnnnnnnnnnnnnnnnn
nnnnnnnnnnnnnnnnnnnnnnnnnnnnnnnnnnnnnnnnnnnnnnnnnnnnnnnnnnnnnnnnnnnnnnnnnnnnnnnnnnnnnnnnn
nnnnnnnnnnnnnnnnnnnnnnnnnnnnnnnnnnnnnnnnnnnnnnnnnnnnnnnnnnnnnnnnnnnnnnnnnnnnnnnnnnnnnnnn
nnnnnnnnnnnnnnnnnnnnnnnnnnnnnnnnnnnnnnnnnnnnnnnnnnnnnnnnnnnnnnnnnnnnnnnnnnnnnnnnnnnnnnnnn
nnnnnnnnnnnnnnnnnnnnnnnnnnnnnnnnnnnnnnnnnnnnnnnnnnnnnnnnnnnnnnnnnnnnnnnnnnnnnnnnnnnnnnnnn
nnnnnnnnnnnnnnnnnnnnnnnnnnnnnnnnnnnnnnnnnnnnnnnnnnnnnnnnnnnnnnnnnnnnnnnnnnnnnnnnnnnnnnnnn
nnnnnnnnnnnnnnnnnnnnnnnnnnnnnnnnnnnnnnnnnnnnnnnnnnnnnnnnnnnnnnnnnnnnnnnnnnnnnnnnnnnnnnnnnn
nnnnnnnnnnnnnnnnnnnnnnnnnnnnnnnnnnnnnnnnnnnnnnnnnnnnnnnnnnnnnnnnnnnnnnnnnnnnnnnnnnnnnnnnn
nnnnnnnnnnnnnnnnnnnnnnnnnnnnnnnnnnnnnnnnnnnnnnnnnnnnnnnnnnnnnnnnnnnnnnnnnnnnnnnnnnnnnnnnn
nnnnnnnnnnnnnnnnnnnnnnCTGGCCAAAAATTTCAATTTACAAGCTCCAGTTTCTCTTGTTTCTCTTCA
TTTGTCCGACTATAGTACTGCTTTGTATCTAGGAGACAGTTAATTGGATTCTCTATTACAGGCATC
TTAATTCCAAAACTTAGCTCCAAAACAATAAAACAGGCTGTTGGAGTTGTGGGTTGCATTATCAT

GCCTCACAATGTGTTCTTGCATTCTGCTCTTGTACAGTCAAGGGAGATTGACCACAATAAGAAAG
GCCAGGTTCAAGAAGCTCTCAGATACTACTCCATAGAGTCAACTGCTGCCCTTGCAATATCATTC
ATGATCAATTTGTTTGTGACGACTGTTTTTGCTAAAGGTTTCCATGGGACAAAACTGGCCAATAG
TATTGGCCTTGTAAATGCAGGGCAATATCTTCAAGATAAATACAGGGGTGGATTTTTCCCAATTT
TATACATCTGGGGTATTGGGTTATTAGCAGCTGGTCAAAGTAGCACCATTACTGGCACCTATGCA
GGGCAGTTTATCATGGGAGGTTTCCTGAACTTGGGATTAAAGAAATGGCAGAGGGCATTGATTA
CTCGAAGCTGTGCTATCATCCCAACTATAATTGTTGCACTTGTTTTTGATACTTCTGAAGACTCAC
TAGATGTTCTGAATGAATGGCTAAATATGCTTCAGTCTATTCAGATTCCTTTTGCACTCATCCCTC
TTCTTTGCTTGGTCTCCAAGGAGCAAATCATGGGCACCTTCACAGTTGGCCCCATTCTTAAGGTA
TGTTGACTAGTCTCTGTAACTCTGTTGTCCAATGTTTCTGTCATTAAACTCAAGATAACCCATTAA
ATTGGTGACAGTAGATTGGATCAAGGAAGCAGTTGCCATTTTCAATTTGTTTGGAAGTTTGTGTA
Annnnnnnnnnnnnnnnnnnnnnnnnnnnnnnnnnnnnnnnnnnnnnnnnnnnnnnnnnnnnnnnnnnnnnnnnnnnnnnnnnnnnnnn
nnnnnnnnnnnnnnnnnnnnnnnnnnnnnnnnnnnnnnnnnnnnnnnnnnnnnnnnnnnnnnnnnnnnnnnnnnnnnnnnnnnnnnnn
nnnnnnnnnnnnnnnnnnnnnnnnnnnnnnnnnnnnnnnnnnnnnnnnnnnnnnnnnnnnnnnnnnnnnnnnnnnnnnnnnnnnnnn
nnnnnnnnnnnnnnnnnnnnnnnnnnnnnnnnnnnnnnnnnnnnnnnnnnnnnnnnnnnnnnnnnnnnnnnnnnnnnnnnnnnnnnn
nnnnnnnnnnnnnnnnnnnnnnnnnnnnnnnnnnnnnnnnnnnnnnnnnnnnnnnnnnnnnnnnnnnnnnnnnnnnnnnnnnnnnnn
nnnnnnnnnnnnnnnnnnnnnnnnnnnnnnnnnnnnnnnnnnnnnnnnnnnnnnnnnnnnnnnnnnnnnnnnnnnnnnnnnnnnnnn
nnnnnnnnnnnnnnnnnnnnnnnnnnnnnnnnnnnnnnnnnnnnnnnnnnnnnnnnnnnnnnnnnnnnnnnnnnnnnnnnnnnnnnn
nnnnnnnnnnnnnnnnnnnnnnnnnnnnnnnnnnnnnnnnnnnnnnnnnnnnnnnnnnnnnnnnnnnnnnnnnnnnnnnnnnnnnnn
nnnnnnnnnnnnnnnnnnnnnnnnnnnnnnnnnnnnnnnnnnnnnnnnnnnnnnnnnnnnnnnnnnnnnnnnnnnnnnnnnnnnnnn
nnnnnnnnnnnnnnnnnnn

>P. mexicana_NRAMP3.2_genomic
nnnnnnnnnnnnnnnnnnnnnnnnnnnnnnnnnnnnnnnnnnnnnnnnnnnnnnnnnnnnnnnnnnnnnnnnnnnnnnnnnnnnnnnn
nnnnnnnnnnnnnnnnnnnnnnnnnnnnnnnnnnnnnnnnnnnnnnnnnnnnnnnnnnnnnnnnnnnnnnnnnnnnnnnnnnnnnnn
nnnnnnnnnnnnnnnnnnnnnnnnnnnnnnnnnnnnnnnnnnnnnnnnnnnnnnnnnnnnnnnnnnnnnnnnnnnnnnnnnnnnnnnn
nnnnnnnnnnnnnnnnnnnnnnnnnnnnnnnnnnnnnnnnnnnnnnnnnnnnnnnnnnnnnnnnnnnnnnnnnnnnnnnnnnnnnnnn
nnnnnnnnnnnnnnnnnnnnnnnnnnnnnnnnnnnnnnnnnnnnnnnnnnnnnnnnnnnnnnnnnnnnnnnnnnnnnnnnnnnnnnnn
nnnnnnnnnnnnnnnnnnnnnnnnnnnnnnnnnnnnnnnnnnnnnnnnnnnnnnnnnnnnnnnnnnnnnnnnnnnnnnnnnnnnnnnn
nnnnnnnnnnnnnnnnnnnnnnnnnnnnnnnnnnnnnnnnnnnnnnnnnnnnnnnnnnnnnnnnnnnnnnnnnnnnnnnnnnnnnnn
nnnnnnnnnnnnnnnnnnnnnnnnnnnnnnnnnnnnnnnnnnnnnnnnnnnnnnnnnnnnnnnnnnnnnnnnnnnnnnnnnnnnnnn
nnnnnnnnnnnnnnnnnnnnnnnnnnnnnnnnnnnnnnnnnnnnnnnnnnnnnnnnnnnnnnnnnnnnnnnnnnnnnnnnnnnnnnn
nnnnnnnnnnnnnnnnnnnnnnnnnnnnnnnnnnnnnnnnnnnnnnnnnnnnnnnnnnnnnnnnnnnnnnnnnnnnnnnnnnnnnnnn
nnnnnnnnnnnnnnnnnnnnnnnnnnnnnnnnnnnnnnnnnnnnnnnnnnnnnnnnnnnnnnnnnnnnnnnnnnnnnnnnnnnnnnnn
nnnnnnnnnnnnnnnnnnnnnnnnnnnnnnnnnnnnnnnnnnnnnnnnnnnnnnnnnnnnnnnnnnnnnnnnnnnnnnnnnnnnnnnn
nnnnnnnnnnnnnnnnnnnnnnnnnnnnnnnnnnnnnnnnnnnnnnnnnnnnnnnnnnnnnnnnnnnnnnnnnnnnnnnnnnnnnnnn
nnnnnnnnnnnnnnnnnnnnnnnnnnnnnnnnnnnnnnnnnnnnnnnnnnnnnnnnnnnnnnnnnnnnnnnnnnnnnnnnnnnnnnnn
nnnnnnnnnnnnnnnnnnnnnnnnnnnnnnnnnnnnnnnnnnnnnnnnnnnnnnnnnnnnnnnnnnnnnnnnnnnnnnnnnnnnnnnn
nnnnnnnnnnnnnnnnnnnnnnnnnnnnnnnnnnnnnnnnnnnnnnnnnnnnnnnnnnnnnnnnnnnnnnnnnnnnnnnnnnnnnnnn
nnnnnnnnnnnnnnnnnnnnnnnnnnnnnnnnnnnnnnnnnnnnnnnnnnnnnnnnnnnnnnnnnnnnnnnnnnnnnnnnnnnnnnnn
nnnnnnnnnnnnnnnnnnnnnnnnnnnnnnnnnnnnnnnnnnnnnnnnnnnnnnnnnnnnnnnnnnnnnnnnnnnnnnnnnnnnnnnn
nnnnnnnnnnnAATTTGTTTGTGACTACTGTTTTTGCTAAAGGTTTCTATGGGACAGAACTGGCCAAT
AGTATTGGCCTTGTAAATGCAGGGCAATATCTTCAAGACAAATATGGGGGTGGATTTTTCCCAAT
TTTATACATCTGGGGTATTGGATTACTAGCAGCTGGCCAAAGCAGCACCATTACCGGAACTTATG
CAGGACAGTTTATCATGGGAGGTTTCCTGAACTTGAGGTTAAAGAAATGGCTGAGGGCATTGAT
CACTCGAAGCTGTGCTATCATCCCAACTATGATTGTTGCACTTGTTTTTGATACCTCCGAAGACTC
ACTAGATGTTCTGAATGAATGGCTAAATGTGCTGCAGTCAATACAGATTCCTTTTGCACTCATCC
CTCTTCTCTGCTTGGTATCCAAGGAGCAAATCATGGGCACTTTCAAAATCAACCCCATTCTTAAG
GTATGTTGACTAGTCTGTTATTCTGCGGTATTTGTCTTGAAGCTTAAGATGGCCCTTTAAATTGGT
GACAGTTGACTAAATCAAGGAACCAATTGCCTTTTTCAATTnTTTGGAATCTCCATATAACCAAA
TCTCTCAACTTGTGTCGTTCTTCAATACTCCAATGCATCATAAAGTCAAAAACATGTGACATTTTG
TTAATGTGTAAACATGTCTGAAATTATGCATGTGATGGATGAATCTTGGGTAAAAAATTGnnnnnnn

nnnnnnnnnnnnnnnnnnnnnnnnnnnnnnnnnnnnnnnnnnnnnnnnnnnnnnnnnnnnnnnnnnnnnnnnnnnnnnnnnnnnnnn
nnnnnnnnnnnnnnnnnnnnnnnnnnnnnnnnnnnnnnnnnnnnnnnnnnnnnnnnnnnnnnnnnnnnnnnnnnnnnnnnnnnnnn
nnnnnnnnnnnnnnnnnnnnnnnnnnnnnnnnnnnnnnnnnnnnnnnnnnnnnnnnnnnnnnnnnnnnnnnnnnnnnnnnnnnnn
nnnnnnnnnnnnnnnnnnnnnnnnnnnnnnnnnnnnnnnnnnnnnnnnnnnnnnnnnnnnnnnnnnnnnnnnnnnnnnnnnnnnn
nnnnnnnnnnnnnnnnnnnnnnnnnnnnnnnnnnnnnnnnnnnnnnnnnnnnnnnnnnnnnnnnnnnnnnnnnnnnnnnnnnnnn
nnnnnnnnnnnnnnnnnnnnnnnnnnnnnnnnnnnnnnnnnnnnnnnnnnnnnnnnnnnnnnnnnnnnnnnnnnnnnnnnnnnnn
nnnnnnnnnnnnnnnnnnnnnnnnnnnnnnnnnnnnnnnnnnnnnnnnnnnnnnnnnnnnnnnnnnnnnnnnnnnnnnnnnnnnn
nnnnnnnnnnnnnnnnnnnnnnnnnnnnnnnnnnnnnnnnnnnnnnnnnnnnnnnnnnnnnnnnnnnnnnnnnnnnnnnnnnnnn
nnnnn

**Supplementary data S3.** Protein sequences used to construct the phylogenetic tree shown in figure 2. $ at the end of the name indicate incomplete protein sequences.

>P. trichocarpa_NRAMP3.1
MPSPEEDPQPLLKDQEETAYDSDGKVLSFGIDYDTESGGSTVVPSFSWRKLWLFTGPGFLMCIAFLDP
GNLEGDLQAGAIAGYSLLWLLLWATAMGLLVQLLSARLGVATGRHLAELCREEYPTWARMILWIM
AELALIGADIQEVIGSAIAIQILSNGVLPLWAGVIITASDCFIFLFLENYGVRKLEAAFGILIGIMAVTFA
WMFADAKPSAPELFLGILIPKLSSKTIKQAVGVVGCIIMPHNVFLHSALVQSREIDHNKKGQVQEALR
YYSIESTAALAISFMINLFVTTIFAKGFHGTELANSIGLVNAGQYLQDKYGGGFFPILYIWGIGLLAAG
QSSTITGTYAGQFIMGGFLNLGLKKWLRALITRSCAIIPTIIVALVFDTSEDSLDVLNEWLNMLQSIQIP
FALIPLLCLVSKEQIMGTFTVGPILKMVSWLVAALVMLINGYLLLDFFSNEVTGVVFTTVVCAFTGAY
VTFIIYLISREVTISTWYCPT

>P. trichocarpa_NRAMP3.2
MPVEENYQPLLQEEEERAYDSDEKVLIIGVDSDTESGGSTVLPPFSWKKLWLFTGPGFLMSIAFLDPG
NLEGDLQAGAIAGYSLLWLLLWATAMGLLVQLLSARLGVATGRHLAELCREEYPTWASMVLWIMA
ELALIGADIQEVIGSAIAIKILSNGFVPLWAGVTITACDCFIFLFLENYGVRKLEAVFAVLIGIMAVTFG
WMFADAKPSASELFLGILIPKLSSRTIQQAVGVVGCIIMPHNVFLHSALVQSREIDHNKKDRVQEALR
YYSIESTTALVISFVINLFVTTVFAKGFYGTELANSIGLVNAGQYLQDKYGGGFFPILYIWGIGLLAAG
QSSTITGTYAGQFIMGGFLNLRLKKWLRALITRSCAIIPTMIVALVFDTSEDSLDVLNEWLNVLQSIQIP
FALIPLLCLVSKEQIMGTFKIGPILKMVAWLVAALVMVINGYLLLDFFFNEVTGVAFTTVVCGFTGAY
VAFIIYLISRGFTCFSRCCPSKQIEVE

>P. tremula_NRAMP3.1
MPLPEEDPQPLLKDQEETAHDSDGKVLSFGIDYDTESGGSTVVPSFSWRKLWLFTGPGFLMCIAFLDP
GNLEGDLQAGAIAGYSLLWLLLWATAMGLLVQLLSARLGVATGRHLAELCREEYPTWARMILWIM
AELALIGADIQEVIGSAIAIQILSNGVLPLWAGVIITASDCFIFLFLENYGVRKLEAAFGILIGIMAVAFA
WMFADAKPSAPELFLGILIPKLSSKTIKQAVGVVGCIIMPHNVFLHSALVQSREIDCNKKGQVQEALR
YYSIESTAALAISFMINLFVTTIFAKGFHGTELANSIGLVNAGQYLQDKYGGGFFPILYIWGIGLLAAG
QSSTITGTYAGQFIMGGFLNLGLKKWLRALITRSCAIIPTIIVALVFDTSEDSLDVLNEWLNMLQSIQIP
FALIPLLCLVSKEQIMGTFTVGPILKMVSWLVAALVMLINGYLLLDFFSNEVTGVVFTTVVCAFTGAY
VTFIIYLISREVSISTWYCPT

>P. tremula_NRAMP3.2
MSVEENQQPLLQEEEERAYDSDEKVLIIGVDSDTESGGSTVLPPFSWKKLWLFTGPGFLMSIAFLDPG
NLEGDLQAGAIAGYSLLWLLFWATAMGLLVQLLSARLGVATGRHLAELCREEYPTWASMVLWIMA
ELALIGADIQEVIGSAIALKILSNGFLPLWAGVTITACDCFIFLFLENYGVRKLEAVFAVLIGIMAVTFG
WMFADAKPSASELFLGILIPKLSSRTIQQAVGVVGCIIMPHNVFLHSALVQSREIDHNKKGRVQEALR

YYSIESTTALVISFVINLFVTTVFAKGFYGTELANSIGLVNAGQYLQDKYGGGFFPILYIWGIGLLAAG
QSSTITGTYAGQFIMGGFLNLRLKKWLRALITRSCAIIPTMIVALVFDASEDSLDVLNEWLNVLQSIQIP
FALIPLLFLVSKEQIMGTFKIGPILKMVSWLVAALVIVINGYLLLDFFVNEVTGVAFTTVVCGFTGAYV
AFIIYLISRGFKCFSWCCQSKQIEVE

>P. tremuloides_NRAMP3.1
MPLPEEDPQPLLKDQEETAYDSDGKVLSFGIDYDTESGGSTVLPPFSWKKLWLFTGPGFLMSIAFLDP
GNLEGDLQAGAIAGYSLLWLLLWATAMGLLVQLLSARLGVATGRHLAELCREEYPTWARMILWIM
AELALIGADIQEVIGSAIAIQILSNGVLPLWAGVIITASDCFIFLFLENYGVRKLEAAFGILIGIMAVTFA
WMFADAKPSAPELFLGILIPKLSSKTIKQAVGVVGCIIMPHNVFLHSALVQSREIDHNKKGQVQEALR
YYSIESTAALAISFMINLFVTTIFAKGFHGTELANSIGLVNAGQYLQDKYGGGFFPILYIWGIGLLAAG
QSSTITGTYAGQFIMGGFLNLGLKKWLRALITRSCAIIPTIIVALVFDTSEDSLDVLNEWLNMLQSIQIP
FALIPLLCLVSKEQLMGTFTVGPILKMVSWLVAALVMLINGYLLLDFFSNEVTGVLFTTVVCAFTGA
YVTFIIYLISREVSISTWYCPT

>P. tremuloides_NRAMP3.2
MPVEENQQPLLQEEEERAYDSDEKVLIIGVDSDTESGGSTVLPPFSWKKLWLFTGPGFLMSIAFLDPG
NLEGDLQAGAIAGYSLLWLLLWATAMGLLVQLLSARLGVATGRHLAELCREEYPTWASMVLWIMA
ELALIGADIQEVIGSAIALKILSNGFLPLWAGVTITACDCFIFLFLENYGVRKLEAVFAVLIGLMAVTFG
WMFADAKPSASELFLGILIPKLSSRTIQQAVGVVGCIIMPHNVFLHSALVQSREIDHNKKGRVQEALR
YYSIESTTALVISFVINLFVTTVFAKGFYGTELANSIGLVNAGQYLQDKYGGGFFPILYIWGIGLLAAG
QSSTITGTYAGQFIMGGFLNLRLKKWLRALITRSCAIIPTMIVALVFDASEDSLDVLNEWLNVLQSIQIP
FALIPLLCLVSKEQIMGTFKIGPILKMVSWLVAALVIVINGYLLLDFFVNEVAGVAFTTVVCGFTGAY
VAFIIYLISRGFTCFSWCCPSKQIEVE

>P. grandidentata_NRAMP3.1
MPLPEEDPQPLLKDQEETAYDSDGKVLSFGIDYDTESGGSTVVPSFSWRKLWLFTGPGFLMCIAFLDP
GNLEGDLQAGAIAGYSLLWLLLWATAMGLLVQLLSARLGVATGRHLAELCREEYPTWARMILWIM
AELALIGADIQEVIGSAIAIQILSNGVLPLWAGVIITASDCFIFLFLENYGVRKLEAAFGILIGIMAVTFA
WMFADAKPSAPELFLGILIPKLSSKTIKQAVGVVGCIIMPHNVFLHSALVQSREIDHNKKGQVQEALR
YYSIESTAALAISFMINLFVTTIFAKGFHGTELANSIGLVNAGQYLQDKYGGGFFPILYIWGIGLLAAG
QSSTITGTYAGQFIMGGFLNLGLKKWLRALITRSCAIIPTIIVALVFDTSEDSLDVLNEWLNMLQSIQIP
FALIPLLCLVSKEQLMGTFTVGPILKMVSWLVAALVMLINGYLLLDFFSNEVTGVVFTTVVCAFTGA
YVTFIIYLISREVSISTWYCPT

>P. grandidentata_NRAMP3.2
MPVEENQQPLLQEEEERAYDSDEKVLIIGVDSDTESGGSTVLPPFSWKKLWLFTGPGFLMSIAFLDPG
NLEGDLQAGAIAGYSLLWLLLWATAMGLLVQLLSARLGVATGRHLAELCREEYPTWASMILWIMA
ELALIGADIQEVIGSAIALKILSNGFLPLWAGVTITACDCFIFLFLENYGVRKLEAVFAVLIGIMAVTFG
WMFADAKPSASELFLGILIPKLSSRTIQQAVGVVGCIIMPHNVFLHSALVQSREIDHNKKGRVQEALR
YYSIESTTALVISFVINLFVTTVFAKGFYGTELANSIGLVNAGQYLQDKYGGGFFPILYIWGIGLLAAG
QSSTITGTYAGQFIMGGFLNLRLKKWLRALITRSCAIIPTMIVALVFDASEDSLDVLNEWLNVLQSIQIP
FALIPLLCLVSKEQIMGTFKIGPILKMVSWLVAALVIVINGYLLLDFFVNEVAGVAFTTVVCGFTGAY
VAFIIYLISRGFTCFSRCCPSKQIEVE

>P. alba_NRAMP3.1

MPLPEEDPQPLLKDQEETAYDSDGKVLSFGIDYDTESGGSTVVPSFSWRKLWLFTGPGFLMCIAFLDP
GNLEGDLQAGAIAGYSLLWLLLWATAMGLLVQLLSARLGVATGRHLAELCREEYPTWARMILWIM
AELALIGADIQEVIGSAIAIQILSNGVLPLWAGVIITASDCFIFLFLENYGVRKLEAAFGILIGIMAVTFA
WMFADAKPSAPELFLGILIPKLSSKTIKQAVGVVGCIIMPHNVFLHSALVQSREIDHNKKGQVQEALR
YYSIESTAALAISFMINLFVTTVFAKGFHGTELANSIGLVNAGQYLQDKYGGGFFPILYIWGIGLLAAG
QSSTITGTYAGQFIMGGFLNLGLKKWLRALITRSCAIIPTIIVALVFDTSEDSLDVLNEWLNMLQSIQIP
FALIPLLCLVSKEQIMGTFTVGPILQMVSWLVAALVMLINGYLLLDFFSNEVTGVVFTTVVCAFTGAY
VTFIIYLISREVTISTWYCPT

>P. alba_NRAMP3.2
MPVEENQQPLLQEEEERAYDSDEKVLIIGVDSDTESGGSTVLPPFSWKKLWLFTGPGFLMSIAFLDPG
NLEGDLQAGAIAGYSLLWLLLWATAMGLLVQLLSARLGVATGRHLAELCREEYPTWASMVLWIMA
ELALIGADIQEVIGSAIALKILSNGFLPLWAGVTITACDCFIFLFLENYGVRKLEAVFAVLIGIMAVTFG
WMFADAKPSASELFLGILIPKLSSRTIQQAVGVVGCIIMPHNVFLHSALVQSREIDHNKKGRVQEALR
YYSIESTTALVISFVINLFVTTVFAKGFYGTELANSIGLVNAGQYLQDKYGGGFFPILYIWGIGLLAAG
QSSTITGTYAGQFIMGGFLNLRLKKWLRALITRSCAIIPTMIVALVFDTSEDSLDVLNEWLNVLQSIQIP
FALIPLLCLVSKEQIMGTFKIGPILKVSWLVAALVIVINGYLLLDFFVNEVAGVAFTTVLCGFTGAYVA
FIIYLISRGFTCFFWCCQSKQIEVE

>P. cathayana_NRAMP3.1
MPLPEEDPQPLLKDQEETAYDSDGKVLSFGIDYDTESGGSTVVPSFSWRKLWLFTGPGFLMCIAFLDP
GNLEGDLQAGAIAGYSLLWLLLWATAMGLLVQLLSARLGVATGRHLAELCREEYPTWARMILWIM
AELALIGADIQEVIGSAIAIQILSNGVLPLWAGVIITASDCFIFLFLENYGVRKLEAAFGILIGIMAVTFA
WMFADAKPSAPELFLGILIPKLSSKTIKQAVGVVGCIIMPHNVFLHSALVQSREIDHNKKGQVQEALR
YYSIESTAALAISFMINLFVTTVFAKGFHGTELANSIGLVNAGQYLQDKYGGGFFPILYIWGIGLLAAG
QSSTITGTYAGQFIMGGFLNLGLKKWLRALITRSCAIIPTIIVALVFDTSEDSLDVLNEWLNMLQSIQIP
FALIPLLCLVSKEQIMGTFTVGPILQMVSWLVAALVMLINGYLLLDFFSNEVTGVVFTTVVCAFTGAY
VTFIIYLISREVTISTWYCPT

>P. cathayana_NRAMP3.2
MPVEENHQPLLQEEEERAYDSDEKVLIIGVDSDTESGGSTVLPPFSWKKLWLFTGPGFLMSIAFLDPG
NLEGDLQAGAIAGYSLLWLLLWATAMGLLVQLLSARLGVATGRHLAELCREEYPTWASMVLWIMA
ELALIGADIQEVIGSAIAIKILSNGFVPLWAGVTITACDCFIFLFLENYGVRKLEAVFAVLIGIMAVTFG
WMFADAKPSASELFLGILIPKLSSRTIQQAVGVVGCIIMPHNVFLHSALVQSREIDHNKKDRVQEALR
YYSIESTTALVISFVINLFVTTVFAKGFYGTELANSIGLVNAGQYLQDKYGGGFFPILYIWGIGLLAAG
QSSTITGTYAGQFIMGGFLNLRLKKWLRALITRSCAIIPTMIVALVFDTSEDSLDVLNEWLNVLQSIQIP
FALIPLLCLVSKEQIMGTFKIGPILKMVAWLVAALVMVINGYLLLDFFFNEVTGVAFTTVVCGFTGAY
AAFIIYLISRGFTCFSRCCPSKQIEVE

>P. simonii_NRAMP3.1
MPLPEEDPRPLLKDQEETAYDSDGKVLSFGIDYDTESGGSTVVPSFSWRKLWLFTGPGFLMCIAFLDP
GNLEGDLQAGAIAGYSLLWLLLWATAMGLLVQLLSARLGVATGRHLAELCREEYPTWARMILWIM
AELALIGADIQEVIGSAIAIQILSNGVLPLWAGVIITASDCFIFLFLENYGVRKLEAAFGILIGIMAVTFA
WMFADAKPSAPELFLGILIPKLSSKTIKQAVGVVGCIIMPHNVFLHSALVQSREIDHNKKGQVQEALR
YYSIESTAALAISFMINLFVTTVFAKGFHGTELANSIGLVNAGQYLQDKYGGGFFPILYIWGIGLLAAG
QSSTITGTYAGQFIMGGFLNLGLKKWLRALITRSCAIIPTTIVALVFDTSEDSLDVLNEWLNMLQSIQIP
FALIPLLCLVSKEQIMGTFTVGPILQMVSWLVAALVMLINGYLLLDFFSNEVTGVVFTTVVCAFTGAY
VTFIIYLISREVTISTWYCPT

>P. simonii_NRAMP3.2
MPVEENHQPLLQEEEERAYDSDEKVLIIGVDSDTESGGSTVLPPFSWKKLWLFTGPGFLMSIAFLDPG
NLEGDLQAGAIAGYSLLWLLLWATAMGLLVQLLSARLGVATGRHLAELCREEYPTWASMVLWIMT
ELALIGADIQEVIGSAIAIKILSNGFVPLWAGVTITACDFIFLFLENYGVRKLEAVFAVLIGIMAVTFGW
MFADAKPSASELFLGILIPKLSSRTIQQAVGVVGCIIMPHNVFLHSALVQSREIDHNKKDRVQEALRY
YSIESTTALVISFVINLFVTTVFAKGFYGTELANSIGLVNAGQYLQDKYGGGFFPILYIWGIGLLAAGQ
SSTITGTYAGQFIMGGFLNLRLKKWLRALITRSCAIIPTMIVALVFDTSEDSLDVLNEWLNVLQSIQIPF
ALIPLLCLVSKEQIMGTFKIGPILKMVSWLVAALVMVINGYLLLDFFFNEVTGVAFTTVVCGFTGAYV
AFIIYLISRGFTCFSRCCPSKQIEVE

>P. lasiocarpa_NRAMP3.1
MPLPEEDPKPLLKDQEETAYDSDGKVLSFGIDYGTESGGSTVVPSFSWRKLWLFTGPGFLMCIAFLDP
GNLEGDLQAGAIAGYSLLWLLLWATAMGLLVQLLSARLGVATGRHLAELCREEYPTWARMILWIM
AELALIGADIQEVIGSAIAIQILSNGVLPLWAGVIITASDCFIFLFLENYGVRKLEAAFGILIGIMAVTFA
WMFADAKPSAPELFLGILIPKLSSKTIKQAVGVVGCIIMPHNVFLHSALVQSREIDHNKKGQVQEALR
YYSIESTAALAISFMINLFVTTVFAKGFHGTELANSIGLVNAGQYLQDKYGGGFFPIFYIWGIGLLAAG
QSSTITGTYAGQFIMGGFLNLGLKKWLRALITRSCAIIPTIIVALVFDTSEDSLDVLNEWLNMLQSIQIP
FALIPLLCLVSKEQIMGTFTVGPILQMVSWLVAALVMLINGYLLLDFFSNEVTGVVFTTVVCAFTGAY
VTFIIYLISREVTISTWYCPT

>P. lasiocarpa_NRAMP3.2
MPVEENHQPLLQEEEERAYDSDEKVLIIGVDSDTESGGSTVLPPFSWKKLWLFTGPGFLMSIAFLDPG
NLEGDLQAGAIAGYSLLWLLLWATAMGLLVQLLSARLGVATGRHLAELCREEYPTWASMVLWIMA
ELALIGADIQEVIGSAIAIKILSNGFVPLWAGVTITACDWFIFLFLENYGVRKLEAVFAVLIGIMAVTFG
WMFADAKPSASELFLGILIPKLSSRTIQQAVGVVGCIIMPHNVFLHSALVQSREIDHNKKDRVQEALR
YYSIESTTALVISFVINLFVTTVFAKGFYGTELANSIGLVNAGQYLQDKYGGGFFPILYIWGIGLLAAG
QSSTITGTYAGQFIMGGFLNLRLKKWLRALITRSCAIIPTMIVALVFDTSEDSLDVLNEWLNVLQSIQIP
FALIPLLCLVSKEQIMGTFKIDPILKMVSWLVAALVMVINGYLLLDFFFNEVTGVAFTTVVCGFTGAY
VAFIIYLISRGFTCFSRCCPSKQIEVE

>P. maximowiczii_NRAMP3.1
MPLPEEDPQPLLKDQEETAYDSDGKVLSFGIDYDTESGGSTVVPSFSWRKLWLFTGPGFLMCIAFLDP
GNLEGDLQAGAIAGYSLLWLLLWATAMGLLVQLLSARLGVATGRHLAELCREEYPTWARMILWIM
AELALIGADIQEVIGSAIAIQILSNGVLPLWAGVIITASDCFIFLFLENYGVRKLEAAFGILIGIMAVTFA
WMFADAKPSAPELFLGILIPKLSSKTIKQAVGVVGCIIMPHNVFLHSALVQSREIDHNKKGQVQEALR
YYSIESTAALAISFMINLFVTTVFAKGFHGTELANSIGLVNAGQYLQDKYGGGFFPILYIWGIGLLAAG
QSSTITGTYAGQFIMGGFLNLGLKKWLRALITRSCAIIPTIIVALVFDTSEDSLDVLNEWLNMLQSIQIP
FALIPLLCLVSKEQIMGTFTVGPILKMVSWLVAALVMLINGYLLLDFFSNEVTGVVFTTVVCAFTGAY
VTFIIYLISREVTISTWYCPT

>P. maximowiczi_NRAMP3.2
MPVEENHQPLLQEEEERAYDSDEKVLIIGVDSDTESGGSTVLPPFSWKKLWLFTGPGFLMSIAFLDPG
NLEGDLQAGAIAGYSLLWLLLWATAMGLLVQLLSARLGVATGRHLAELCREEYPTWASMVLWIMA
ELALIGADIQEVIGSAIAIKILSNGFVPLWAGVTITACDCFIFLFLENYGVRKLEAVFAVLIGIMAVTFG
WMFADAKPSASELFLGILIPKLSSRTIQQAVGVVGCIIMPHNVFLHSALVQSREIDHNKKDRVQEALR
YYSIESTTALVISFVINLFVTTVFAKGFYGTELANSIGLVNAGQYLQDKYGGGFFPILYIWGIGLLAAG

QSSTITGTYAGQFIMGGFLNLRLKKWLRALITRSCAIIPTMIVALVFDTSEDSLDVLNEWLNVLQSIQIP
FALIPLLCLVSKEQIMGTFKIGPILKMVAWLVAALVMVINGYLLLDFFFNEVTGVAFTTVVCGFTGAY
AAFIIYLISRGFTCFSRCCPSKQIEVE

>P. euphratica_NRAMP3.1
MPLPEEDPQPLLEDQEETAYDSNGKVLSFGIDYDTESGGSTVVPSFSWRKLWLFTGPGFLMCIAFLDP
GNLEGDLQAGAIAGYSLLWLLLWATAMGLLVQLLSARLGVATGRHLAELWREE*PTWARMILWIM
AELALIGADIQEVIGSAIAIQILSNGVLPLWAGVIITASDSFIFLFLENYGVRKLEAAFGILIGIMAVTFAC
MFADAKPSAPELFLGILIPKLSSKTIKQAVGVVGCIIMPHNVFLHSALVQSREIDHNKKGRVQEALRY
YSIESTAALAISFMNNLFVTTVFAKGFHRTELANSIGLVNAGQYLQDKYGGGFFPILYIWGIGLLAAG
QSSTITGTYAGQFIMGGFLNLGLKKWLRALITRSCAIVPTIIVALVFDTSEDSLDVLNEWLNMLQSIQIP
FALIPLLCLVSKEQIMGTFTVGPILKMVSWFVAALVMLINGYLLLDFFSNEVTGVVFITVVCAFTGAY
VTFTIYLISREVTISTWYCPS

>P. euphratica_NRAMP3.2
MPVEENHQPLLQEEEERAYDSDEKVLIIGVDSDTESGGSTVLPPFSWKKLWLFTGPGFLMSIAFLDPG
NLEGDLQAGAIAGYSLLWLLLWATAMGLLVQLLSARLGVATGRHLAELCREEYPTWASMVLWIMA
ELALIGADIQEVIGSAIAIKILSNGFVPLWAGVTITACDCFIFLFLENYGVRKLEAVFAVLIGIMAVTFG
WMFADAKPSASELFLGILIPKLSSRTIQQAVGVVGCIIMPHNVFLHSALVQSREIDHNKKDRVQEALR
YYSIESTTALVISFVINLFVTTVFAKGFYGTELANSIGLVNAGQYLQDKYGGGFFPILYIWGIGLLAAG
QSSTITGTYAGQFIMGGFLNLRLKKWLRALITRSCAIIPTMIVALVFDTSEDSLDVLNEWLNVLQSIQIP
FALIPLLCLVSKEQIMGTFKIGPILKMVAWLVAALVMVINGYLLLDFFFNEVTGVAFTTVVCGFTGAY
AAFIIYLISRGFTCFSRCCPSKQIEVE

>P. ussuriensis_NRAMP3.1
MPLPEEDPQPLLKDQEETAYDSDGKVLSFGIDYDTESGGSTVVPSFSWRKLWLFTGPGFLMCIAFLDP
GNLEGDLQAGAIAGYSLLWLLLWATAMGLLVQLLSARLGVATGRHLAELCREEYPTWARMILWIM
AELALIGADIQEVIGSAIAIQILSNGVLPLWAGVIITASDCFIFLFLENYGVRKLEAAFGILIGIMAVTFA
WMFADAKPSAPELFLGILIPKLSSKTIKQAVGVVGCIIMPHNVFLHSALVQSREIDHNKKGQVQEALR
YYSIESTAALAISFMINLFVTTVFAKGFHGTELANSIGLVNAGQYLQDKYGGGFFPILYIWGIGLLAAG
QSSTITGTYAGQFIMGGFLNLGLKKWLRALITRSCAIIPTIIVALVFDTSEDSLDVLNEWLNMLQSIQIP
FALIPLLCLVSKEQIMGTFTVGPILQMVSWLVAALVMLINGYLLLDFFSNEVTGVVFTTVVCAFTGAY
VTFIIYLISREVTISTWYCPT

>P. ussuriensis_NRAMP3.2
MPVEENHQPLLQEEEERAYDSDEKVLIIGVDSDTESGGSTVLPPFSWKKLWLFTGPGFLMSIAFLDPG
NLEGDLQAGAIAGYSLLWLLLWATAMGLLVQLLSARLGVATGRHLAELCREEYPTWASMVLWIMA
ELALIGADIQEVIGSAIAIKILSNGFVPLWAGVTITACDCFIFLFLENYGVRKLEAVFAVLIGIMAVTFG
WMFADAKPSASELFLGILIPKLSSRTIQQAVGVVGCIIMPHNVFLHSALVQSREIDHNKKDRVQEALR
YYSIESTTALVISFVINLFVTTVFAKGFYGTELANSIGLVNAGQYLQDKYGGGFFPILYIWGIGLLAAG
QSSTITGTYAGQFIMGGFLNLRLKKWLRALITRSCAIIPTMIVALVFDTSEDSLDVLNEWLNVLQSIQIP
FALIPLLCLVSKEQIMGTFKIGPILKMVAWLVAALVMVINGYLLLDFFFNEVTGVAFTTVVCGFTGAY
AAFIIYLISRGFTCFSRCCPSKQIEVE

>P. nigra_NRAMP3.1
MPVPEEDPQPLLKDQEETAYDSDGKVLSFGIDYDTESGGSTVVPSFSWRKLWLFTGPGFLMCIAFLDP
GNLEGDLQAGAIAGYSLLWLLLWATAMGLLVQLLSARLGVATGRHLAELCREEYPTWARMILWIM
AELALIGADIQEVIGSAIAIQILSNGVLPLWAGVIITASDCFIFLFLENYGVRKLEAAFGILIGIMAVTFA

WMFADAKPSAPELFLGILIPKLSSKTIKQAVGVVGCIIMPHNVFLHSALVQSREIDHNKKGQVQEALR
YYSIESTAALAISFMINLFVTTVFAKGFHGTELANSIGLVNAGQYLQDKYGGGFFPILYIWGIGLLAAG
QSSTITGTYAGQFIMGGFLNLGLKKWLRALITRSCAIIPTIIVALVFDTSEDSLDVLNEWLNMLQSIQIP
FALIPLLCLVSKEQIMGTFTVGPILQMVSWLVAALVMLINGYLLLDFFSNEVTGVAFTTVVCAFTGAY
VAFIIYLISREVTISTWYCPT

>P. nigra_NRAMP3.2
MPVEENHQPLLQEEEERAYDSDEKVLIIGVDSDTESGSSTVLPPFSWKKLWLFTGPGFLMSIAFLDPG
NLEGDLQAGAIAGYSLLWLLLWATAMGLLVQLLSARLGVATGRHLAELCREEYPTWASMVLWIMA
ELALIGADIQEVIGSAIAIKILSNGFVPLWAGVTITACDWFIFLFLENYGVRKLEAVFAVLIGIMAVTFG
WMFADAKPSASELFLGILIPKLSSRTIQQAVGVVGCIIMPHNVFLHSALVQSREIDHNKKDRVQEALR
YYSIESTTALVISFVINLFVTTVFAKGFYGTELANSIGLVNAGQYLQDKYGGGFFPILYIWGIGLLAAG
QSSTITGTYAGQFIMGGFLNLRLKKWLRALITRSCAIIPTMIVALVFDTSEDSLDVLNEWLNVLQSIQIP
FALIPLLCLVSKEQIMGTFKIGPTLQMVSWLVAALVMVINGYLLLDFFFNEVTGVAFTTVVCGFTGA
YVAFIIYLISRGFTCFSRCCSSKQIEVE

>P. deltoides_NRAMP3.1
MPLPEEDPQPLLKDQEETAYDSDGKVLSFGIDYDTESGSSTVVPSFSWRKLWLFTGPGFLMCIAFLDP
GNLEGDLQAGAIAGYSLLWLLLWATAMGLLVQLLSARLGVATGRHLAELCREEYPTWARMILWIM
AELALIGADIQEVIGSAIAIQILSNGVLPLWAGVIITASDCFIFLFLENYGVRKLEAAFGILIGIMAVTFA
WMFADAKPSAPELFLGILIPKLSSKTIKQAVGVVGCIIMPHNVFLHSALVQSREIDHNKKGQVQEALR
YYSIESTAALAISFMINLFVTTVFAKGFHGTELANSIGLVNAGQYLQDKYGGGFFPILYIWGIGLLAAG
QSSTITGTYAGQFIMGGFLNLGLKKWLRALITRSCAIIPTIIVALVFDTSEDSLDVLNEWLNMLQSIQIP
FALIPLLCLVSKEQIMGTFTVGPILQMVSWLVAALVMLINGYLLLDFFSNEVTGVVFTTVVCAFTGAY
VTFIIYLISREVTISTWYCPT

>P. deltoides_NRAMP3.2
LQAGAIAGYSLLWLLLWATAMGLLVQLLSARLGVATGRHLAELCREEYPTWASMVLWIMAELALI
GADIQEVIGSAIAIKILSNGFVPLWAGVTITACDCFIFLFLENYGVRKLEAVFAVLIGIMAVTFGWMFA
DAKPSASELFLGILIPKLSSRTIQQAVGVVGCIIMPHNVFLHSALVQSREIDHNKKGRVQEALRYYSIES
TTALVISFVINLFVTTVFAKGFYGTELANSIGLVNAGQYLQDKYGGGFFPILYIWGIGLLAAGQSSTIT
GTYAGQFIMGGFLNLRLKKWLRALITRSCAIIPTMIVALVFDTSEDSLDVLNEWLNVLQSIQIPFALIPL
LCLVSKEQIMGTFKIGPILQMVAWLVAALVMVINGYLLLDFFFNEVTGVAFTTVVCGFTGAYAAFIIY
LISRGFTCFSRCCPSKQIEVE

>S. purpurea_NRAMP3
MSLDENQQPLLQEEEERAYDSDEKVLVIGIDSDAESGGTVLPPFSWKKLWLFTGPGFLMSIAFLDPGN
LEGDLQAGAIAGYSLLWLLFWATAMGLLVQLLSARLGVATGRHLAELCREEYPTWARMILWIMAE
LALIGADIQEVIGSAIAIKILSNGVVPLWAGVTITACDCFIFLFLENYGVRKLEAVFAVLIGVMAVTFG
WMFADAKPSASELFLGILIPKLSSRTIQQAVGVVGCIIMPHNVFLHSALVQSREIDHNNKVQVQEALR
YYSIESTTALVISFVINLFVTTVFAKGFYGTELANSIGLVNAGQYLQDKYGGGFFPILYIWGIGLLAAG
QSSTITGTYAGQFIMGGFLNMRLKKWLRALITRSCAIIPTIIVALIFDTSEDSLDVLNEWLNVLQSIQIPF
ALIPLLCLVSKERIMGTFKIGSILKVVSWLVAALVIVINGYLLLDFFFNEVTGVAFTTVVCTFTAAYAA
FIIYLTSRGVTCSSWRGPPKQIEVE

>S. brachista_NRAMP3

MSLDENQQPLLQEEEERAYDSDQKVLVIGIDSDAESGGTVLPPFSWKKLWLFTGPGFLMSIAFLDPGN
LEGDLQAGAIAGYSLLWLLLWATAMGLLVQLLSARLGVATGRHLAELCREEYPTWARMILWIMAE
LALIGADIQEVIGSAIAIKILSNGVVPLWAGVTITACDCFIFLFLENYGVRKLEAVFAVLIGVMAVTFG
MMFADAKPSASELFLGILIPKLSSRTIQQAVGVVGCIIMPHNVFLHSALVQSREIDHNNKVQVQEALR
YYSIESTTALVISFVINLFVTTVFAQGFYGTELANSIGLVNAGQYLQDKYGGGFFPILYIWGIGLLAAG
QSSTITGTYAGQFIMGGFLNMRLKKWLRALITRSCAIIPTIIVALVFETSEESLDVLNEWLNVLQSIQIPF
ALIPLLCLVSKERIMGTFKIGSILKVVSWLVAALVIIINGYLLLDFFFNEVTGVAYTTAVCTFTAAYAA
FIIYLTSRGVTCSSWRGPPKQNQQIEVE

>S. suchowensis_NRAMP3
MSLDENQQPLLQEEEERAYDSDEKVLVIGIDSDAESGRTVLPPFSWKKLWLFTGPGFLMSIAFLDPGN
LEGDLQAGAIAGYSLLWLLLWATAMGLLVQLLSARLGVATGRHLAELCREEYPTWARMILWIMAE
LALIGADIQEVIGSAIAIKILSNGVVPLWAGVTITACDCFIFLFLENYGVRKLEAVFAVLIGVMAVTFG
WMFADAKPSASELFLGILIPKLSSRTIQQAVGVVGCIIMPHNVFLHSALVQSREIDHNNKVQVQEALR
YYSIESTTALVISFVINLFVTTVFAKGFYGTELANSIGLVNAGQYLQDKYGGGFFPILYIWGIGLLAAG
QSSTITGTYAGQFIMGGFLNMRLKKWLRALITRSCAIIPTIIVALIFDTSEDSLDVLNEWLNVLQSIQIPF
ALIPLLCLVSKERIMGTFKIGSILKVVSWLVAALVIVINGYLLLDFFFNEVTGVAFTTAVCTFTAAYAA
FIIYLTSRGITCSSWRGPPKQIEAE

>S. eriocephala_NRAMP3
MSLDENQQALLQEEEERAYDSDEKVLVIGVYSDAESGGTVLPPFSWKKLWLFTGPGFLMSIAFLDPG
NLEGDLQAGAIAGYSLLWLLLWATAMGLLVQLLSARLGVATGRHLAELCREEYPTWARMILWIMA
ELALIGADIQEVIGSAIAIKILSNGVVPLWAGVTITACDCFIFLFLENYGVRKLEAVFAVLIGVMAVTFG
WMFADAKPSASELFLGILIPKLSSRTIQQAVGVVGCIIMPHNVFLHSALVQSREIDHNNKIQVQEALRY
YSIESTTALVISFVINLFVTTVFAKGFYGTELANSIGLVNAGQYLQDKYGGGFFPILYIWGIGLLAAGQ
SSTITGTYAGQFIMGGFLNMRLKKWIRALITRSCAIIPTIIVALIFDTSEDSLDVLNEWLNVLQSIQIPFA
LIPLLCLVSKERIMGTFKIGSILKVVSWLVAALVMVINGYLLLDFFFNEVTGVAFTTAVCTFTAAYAA
FIIYLTSRGVTCSSWRGPPKQIEVE

>S. sachalinensis_NRAMP3
MSLDENQQPLLQEEEERAYDSDEKVLVIGIDSDAESGGTVLPPFSWKKLWLFTGPGFLMSIAFLDPGN
LEGDLQAGAIAGYSLLWLLLWATAMGLLVQLLSARLGVATGRHLAELCREEYPTWARMILWIMAE
LALIGADIQEVIGSAIAIKILSNGVVPLWAGVTITACDCFIFLFLENYGVRKLEAVFAVLIGVMAVTFG
WMFADAKPSASELFLGILIPKLSSRTIQQAVGVVGCIIMPHNVFLHSALVQSREIDHNNKVQVQEALR
YYSIESTTALVISFVINLFVTTVFAKGFYGTELANSIGLVNAGQYLQDKYGGGFFPILYIWGIGLLAAG
QSSTITGTYAGQFIMGGFLNMRLKKWLRALITRSCAIIPTIIVALIFDTSEDSLDVLNEWLNVLQSIQIPF
ALIPLLCLVSKERIMGTFKIGSILKVVSWLVAALVIVINGYLLLDFFFNEVIGVAFTTAVCTFTAAYAA
FIIYLTSRGITCSSWRGPPKQIEAE

>S. dasyclados_NRAMP3[$]
MAVTFGWMFADAKPSASELFLGILIPKLSSRTIQQAVGVVGCIIMPHNVFLHSALVQSREIDHNNKVQ
VQEAVRYYSIESTTALVISFVINLFVTTVFAQGFYGTELANSIGLVNAGQYLQDKYGGGFFPILYIWGI
GLLAAGQSSTITGTYAGQFIMGGFLNMRLKKWLRALITRSCAIIPTIIVALIFDTSEDSLDVLNEWLNV
LQSIQIPFALIPLLCLVSKERIMGTFKIGSILKVVSWLVAALVIVINGYLLLDFFFNEVTGVAFTTAVCTF
TAAYAAFIIYLTSRGITCSSWRGPPKQIEVE

>S. fargesii_NRAMP3$
MSIAFLDPGNLEGDLQAGAIAGYSLLWLLLWATAMGLLVQLLSARLGVATGRHLAELCREEYPTWARMILWIMAELALIGADIQEVIGSAIAIKILSNGVVPLWAGVTITACDCFIFLFLENYGVRKLEAVFAVLIGVMAVTFGMMFADAKPSASELFLGILIPKLSSRTIQQAVGVVGCIIMPHNVFLHSALVQSREIDHNNKVQVQEALRYYSIESTTALVISFVINLFVTTVFAQGFYGTELANSIGLVNAGQYLQDKYGGGFFPILYIWGIGLLAAGQSSTITGTYAGQFIMGGFLNMRLKKWLRALITRSCAIIPTIIVALVFETSEESLDVLNEWLNVLQSIQIPFALIPLLCLVSKERIMGTFKIGSILKVVSWLVAALVIIINGYLLLDFFFNEVTGVAYTTAVCTFTAAYAAFIIYLTSRGVTCSSWRGPPKQNQQIEVE

>S. viminalis_NRAMP3$
MSLDENQQPLLQEEEERAYDSDEKVLVIGIDSDAESSGTVLPPFSWKKLWLFTGPGFLMSIAFLDPGNLEGDLQAGAIAGYSLLWLLLWATAMGLLVQLLSARLGVATGRHLAELCREEYPTWARMILWIMAELALIGADIQEVIGSAIAIKILSNGVVPLWAGVTITACDCFIFLFLENYGVRKLEAVFAVLIGVMAVTFGWMFADAKPSASELFLGILIPKLSSRTIQQAVGVVGCIIMPHNVFLHSALVQSREIDHNNKVQVQEAVRYYSIESTTALVISFVINLFVTTVFAQGFYGTELANSIGLVNAGQYLQDKYGGGFFPILYIWGIGLLAAGQSSTITGTYAGQFIMGGFLNMRLKKWLRALITRSCAIIPTIIVALIFDTSEDSLDVLNEWLNVLQSIQIPFALIPLLCLVSKERIMGTFKIGSILKVVSWLVAALVIVINGYLLLDFFF*

### Supplementary materials and methods

*Gene expression*

Total RNA from 1-month-old poplars or 1 week-old *A. thaliana* grown *in vitro* on half strength Murashige and Skoog (MS) or ABIS medium, respectively, was extracted using RNeasy Plant Mini Kit (Qiagen) as previously described (Pottier et al., 2015a). Five micrograms of DNA-free RNA were used for reverse transcription by the SuperScript III First-Strand kit (Invitrogen) using random hexamers. Primers were designed using OligoPerfect™ Designer (http://tools.lifetechnologies.com) and their specificity was confirmed by analysis of the melting curves and sequencing of the PCR products. qPCR reactions were performed on a Roche LightCycler 96 using primers listed in supplementary table S7 and Roche reagents, according to the manufacturer's instructions (http://www.roche.com). Relative transcript levels were calculated by normalization to the transcript amount of constitutively expressed genes.

*Yeast strains, transformations and media*

DEY1453 (*fet3fet4*, (Eide et al. 1996), *smf1* (Supek et al. 1996; Thomine et al. 2000) and *smf2* (Cohen et al. 2000) yeast strains were grown at 30ºC on Yeast extract/Peptone/Dextrose (before transformation) or synthetic dextrose -ura (after transformation). For the DEY1453 strain, media were supplemented with 0.2 mM $FeCl_3$. Yeast cells were transformed according to standard procedures (Invitrogen).

*Poplar transformation and regeneration*

The poplar INRA 717-1-B4 clone (*P. tremula* x *P. alba*) was transformed as indicated in supplementary material and methods and using media listed in Supplementary table S8 (Leplé et al. 1992). Briefly, stem explants were excised from *in vitro* grown poplar and incubated for 2 days in the dark on M1 medium. The explants were then co-cultivated for 16 h at 24°C on an orbital shaker (180 rpm) with 150 ml of a suspension ($OD_{600}$ = 0.3) of *A. tumefaciens* C58/pMP90 transformed with pMDC83 constructs. The explants were washed 5 times in sterile water under orbital shaking (140 rpm). After washing, they were transferred on M2 medium containing Ticarpen, cefotaxime and hygromycin B, and incubated at 24°C in darkness for 21 days. The plates were subsequently transferred to light (16 h 130 µE.m$^{-2}$/8h dark) and generated green calli were transferred to M3 agar medium with hygromycin B (Supplementary table S8). The regenerated transformed shoots were then transferred on half strength MS agar medium with hygromycin to allow root regeneration. The transgenic lines were propagated on the same medium until transfer to pots in the greenhouse.
